# Convergent strategies for nanobody-mediated inhibition of an epoxide hydrolase

**DOI:** 10.64898/2026.02.09.704437

**Authors:** Adam R. Simard, Noor M. Taher, Kathryn S. Beauchemin, Akaash K. Mishra, Andrew P. Hederman, Thomas H. Hampton, Natalia Vasylieva, Christophe Morisseau, Margaret E. Ackerman, Dean R. Madden

## Abstract

Secreted by *Pseudomonas aeruginosa*, Cif is an epoxide hydrolase that acts as a virulence factor in the context of cystic fibrosis and thus represents a target for therapeutic inhibition. Here, we present the structures of several high-affinity inhibitory nanobodies, each bound to Cif. Comparison reveals two classes of nanobodies with distinct CDR sequences and convergent recognition strategies. Mimicry between CDR3 and CDR2 loops positions an aromatic residue for insertion through the active-site gate, accessing a cryptic epitope, which sterically blocks substrate access and provides an anchor point for high-affinity engagement. Projection of either inhibitory CDR toward the active-site entrance requires a relative 90° rotation of the core immunoglobulin domain, and yet both classes engage the same set of stereochemical handholds within a highly overlapping shared epitope. The structurally distinct paratopes thus represent fundamentally distinct solutions, reflecting the remarkable capacity of the immune system to solve highly constrained molecular recognition challenges.

## Introduction

Like many successful opportunistic pathogens, *Pseudomonas aeruginosa* (PA) has developed an arsenal of virulence factors allowing it to thrive in a wide range of environments, including the airways of patients with cystic fibrosis (CF) and other diseases that impair mucociliary clearance.^1^ The resulting chronic infections and acute exacerbations lead to progressive tissue damage and patient morbidity and mortality.^2,3^ While highly effective modulator therapies (HEMTs) restore lung homeostasis and improve lung function in CF,^4,5^ chronic PA infections persist.^6–8^ As a result, PA virulence mechanisms remain an important focus of translational research.^9^

The CF transmembrane conductance regulator (CFTR) inhibitory factor (Cif) is a PA virulence enzyme that carries out a two-pronged assault. Once delivered to bronchial epithelial cells by outer membrane vesicles, Cif interferes with endocytic trafficking of CFTR causing its premature degradation in the lysosome,^10–13^ potentially reversing the action of HEMTs,^14^ even if the effect of PA infection on CFTR expression levels may involve additional factors *in vivo.*^15–18^ This has been linked to defective ciliary beating in the mucociliary escalator which in turn enhances airway colonization by PA.^19^ Cif also destroys the signaling molecule 14,15-epoxyeicosatrienoic acid (14,15-EET), which stimulates neutrophil secretion of 15-epi-lipoxin A_4_, a pro-resolving regulatory signal.^18,20,21^ By reducing apical CFTR and promoting a hyperinflammatory environment, Cif compounds the CF phenotype and has the potential to attenuate the therapeutic benefit of HEMT.

Cif is a homodimeric epoxide hydrolase belonging to the α/β-hydrolase superfamily.^22^ Catalytic activity is required for Cif’s virulence effects, as loss of enzymatic function attenuates PA virulence both *in vitro*^23^ and *in vivo.*^18,19^ This has led to a search for inhibitors, using both small molecules and biologics, including development of a panel of Cif-specific nanobodies with inhibitory activity.^24^

Nanobodies are the variable heavy (V_H_) domain of camelid derived heavy-chain only antibodies (also known as VHHs) and represent the smallest binding unit of immunoglobulin-derived fragments.^25–27^ At first approximation, the smaller number of complementarity determining regions (CDRs) available for antigen binding should be detrimental to function, yet nanobodies retain the specificity and high affinity of their larger, conventional antibody (Ab) counterparts. Furthermore, the flat paratopes of conventional Abs present an inherent mismatch to the concavity of enzyme active sites.^28^ Nanobodies often overcome these hurdles through an extra-long CDR3 with exceptional conformational diversity.^29^ With respect to active-site pockets and clefts, this enhances shape complementarity, which is thought to compensate for the missing light-chain contributions.^30–32^ These attributes make nanobodies adept at binding small protein antigens and inhibiting enzymes.^33–35^ Due to facile expression and purification, they can readily serve as reagents for myriad research tools such as immunassays^36–43^

KB2115, a known competitive Cif inhibitor,^44^ was able to displace several of our Cif-specific nanobodies. To further test a hypothesis of competitive inhibition, we leveraged nanobody displacement to develop a small-molecule screen with sufficient sensitivity to delineate the performance of rationally designed KB2115 derivatives.^24^ However, the Cif active site is guarded by a flexible steric “gate” at its entry, raising questions of the nature of the nanobody engagement. In addition, we lacked fine-grained detail for the Cif:nanobody interactions to validate the basis of the screen and to serve as a framework for future applications.

In this study, we determined crystal structures of two Cif-specific nanobodies and five Cif:nanobody complexes to define the mechanisms of nanobody-mediated recognition and inhibition of Cif and to better understand the structural constraints governing the steric interplay between competitive small-molecule and biologic inhibitors. We categorized inhibitory nanobodies into two classes and identified a preferred mechanism for inhibition carried out through structural mimicry between CDR3 and CDR2 loop conformations. We also conclude that despite structurally distinct paratopes, both classes of nanobodies latch onto Cif using nearly identical stereochemical attachment points found throughout a shared core epitope, with implications for molecular recognition and paratope design.

## Materials and Methods

### Cloning and Generation of Stable Protein Production Stocks

All protein coding sequences were cloned into a cytosolic expression system that utilizes an N-terminal deca-histidine tagged SUMO fusion (10xHis-SUMO).^45^ Nanobody sequences were PCR amplified from plasmids isolated from the initial library construction^24^ or were purchased as gBlocks from Integrated DNA Technologies. The coding sequence of mature Cif (UniProtKB accession number A0A0H2ZD27, residues 25 to 319) lacking the secretion signal was PCR amplified from pDPM73.^11^ The resulting amplicons were inserted by Gibson assembly into pCDB24 or pARS6 (a pCDB24 derivative lacking the yeast origin of replication and uracil synthesis cassette) digested by XhoI or XhoI/BamHI, and the resulting product was transformed into chemically competent *Escherichia coli* DH5α by heat shock and recovered on lysogeny broth (LB)^46^ carbenicillin (100 μg/mL) agar plates. Plasmids were isolated using a QIAprep Spin Miniprep Kit from QIAGEN following the manufacturer’s specifications, and fidelity of the insert was verified by Sanger sequencing. pCDB24 was a gift from Dr. Christopher Bahl (https://www.addgene.org/91959/ ; RRID:Addgene_91959).

Once each construct was verified, a stable production glycerol stock was prepared. Plasmids were transformed into chemically competent *E. coli* BL21 (DE3) or *E. coli* BL21 (DE3)-CodonPlus RIL cells and recovered on LB carbenicillin (100 μg/mL) agar plates or LB carbenicillin (100 μg/mL) + chloramphenicol (34 μg/mL) agar plates, respectively. Isolates were inoculated into 5 mL ZYP media^47^ supplemented with 0.8% (*w/v*) dextrose (ZYP-0.8G) and the appropriate antibiotic and then incubated at 37°C overnight. Glycerol stocks were prepared with 9% (*v/v*) glycerol and slowly cooled to -80°C. Small-scale expression tests were performed in Terrific Broth (TB)^48^ and induced with 0.1 mM isopropyl β-D-1-thiogalactopyranoside (IPTG). Whole-cell lysates were assessed for expression by reducing SDS-PAGE, and production stocks for each construct were selected based on band intensity and total biomass as measured by optical density at 600 nm (OD_600_).

### Expression and Purification of Recombinant Proteins

Cif was expressed either as a secreted protein with a C-terminal hexa-histidine tag (Cif-6xHis) or as a cytosolic N-terminal deca-histidine-SUMO fusion protein (10xHis-SUMO-Cif). Cif-6xHis was expressed and purified as previously described.^49^ In brief, *E. coli* TOP10 cells harboring pDPM73 were grown in TB carbenicillin (100 μg/mL) supplemented with 0.2% (*w/v*) L-arabinose at 37°C for 72 h. Supernatant was harvested by centrifugation at 10,000 x *g* at 4°C for 20 min. Recombinant protein was purified from filter-sterilized supernatant by Ni-IMAC followed by size-exclusion chromatography (SEC) on a HiLoad Superdex 200 26/60 PG column as needed. Purified Cif-6xHis was stored at 4°C in buffer [20 mM sodium phosphate, 100 mM NaCl, pH 7.4].

Nanobodies were expressed either as cytosolic N-terminal deca-histidine-SUMO fusion proteins or secreted as C-terminal hexa-histidine-HA fusion proteins. Expression and purification of all recombinant VHH and Cif 10xHis-SUMO fusion proteins were adapted from the procedures outlined below for the 10xHis-SUMO-Cif construct. Different buffer systems were chosen for select nanobodies and are noted below. Recombinant 10xHis-SUMO-Cif from *E. coli* BL21 (DE3) expressed from the pCDB24 plasmid was performed as follows. The frozen production stock was struck to single colonies on LB carbenicillin (100 μg/mL) agar plates and incubated at 37°C overnight. A single colony was used to inoculate 5 mL ZYP-0.8G carbenicillin (100 μg/mL) and incubated at 37°C overnight. The overnight culture was used to seed a propagation culture in TB carbenicillin (100 μg/mL) at a 1:100 dilution and grown at 37°C with vigorous shaking until an OD_600_ of 4 was achieved. The propagation culture was diluted 1:10 into pre-warmed TB carbenicillin (100 μg/mL) and incubated at 37°C with vigorous shaking. Once an OD_600_ of 3 - 4 was reached, cultures were induced with 0.1 mM IPTG, and expression was allowed to proceed at 18°C with vigorous shaking for 16 - 18 h. Cells were harvested by centrifugation at 7,000 x *g* at 4°C for 15 min and resuspended in lysis buffer [50 mM sodium phosphate, 500 mM NaCl, 60 mM imidazole, 2 mM MgCl_2_, 1 mM dithiothreitol (DTT), pH 7.4] supplemented with 25 U/mL Pierce Universal Nuclease and Pierce EDTA-free Protease Inhibitor Tablets per the manufacturer’s specifications. Cells were lysed using a microfluidizer and the lysate was clarified by centrifugation at 40,000 RPM at 4°C for 1 h using a Ti-45 rotor. The recombinant 10xHis-SUMO-Cif was first purified by Ni-IMAC as follows: clarified lysate was loaded onto equilibrated Ni-NTA resin, washed [50 mM sodium phosphate, 500 mM NaCl, 60 mM imidazole, pH 7.4], and eluted by gradient elution with wash buffer with increasing concentrations of imidazole from 60 mM to 500 mM. Fractions containing the 10xHis-SUMO-Cif fusion were pooled and the tag was cleaved by a deca-histidine tagged Cth SUMO protease^50^ during dialysis against buffer containing [20 mM sodium phosphate, 100 mM NaCl, pH 7.4]. Mature Cif was separated by Ni-IMAC and further purified by SEC using a HiLoad Superdex 200 26/60 PG column. Fractions containing pure Cif (assessed by reducing SDS-PAGE) were pooled, concentrated, dialyzed into buffer containing [20 mM sodium phosphate, 100 mM NaCl, pH 7.4], and stored at 4°C. Purification of nanobodies VHH101, VHH113, VHH114, VHH212, VHH214, VHH222, VHH314, and VHH320 were carried out using a sodium phosphate pH 7.4 buffer system with NaCl at concentrations ranging from 100 mM to 500 mM. Purification of VHH108 was carried out using a Tris pH 8.0 buffer system and 100 mM NaCl. Purification of VHH219 was carried out using a Tris pH 8.5 buffer system and 100 mM NaCl.

VHH101 and VHH113 carried on the pComb3x vector or pET28a(+) vector,^24^ respectively, were expressed for crystallization trials as secreted proteins and purified from the culture medium as follows. *E. coli* BL21-CodonPlus (DE3) RIL cells harboring the recombinant protein were grown in TB carbenicillin (100 μg/mL) (VHH101) or TB kanamycin (50 μg/mL) (VHH113) at 37°C until an OD_600_ of 0.6 was reached. Cultures were induced with 0.1 mM IPTG and expressed at 37°C (VHH101) or 18°C (VHH113) for 72 h. Cells were removed by centrifugation at 5,500 x *g* for 30 min at 4°C (VHH101) or 10,100 x *g* for 30 min at 4°C. Residual cell debris was removed from the supernatant using a 0.22 μm filter, and the filtrate was supplemented with imidazole to a final concentration of 5 mM. Recombinant protein was then purified from the filtered culture medium by Ni-IMAC with gradient elution in a Tris buffer system [20 mM Tris, 500 mM NaCl, pH 8.5]. VHH101 was further purified by SEC using a HiLoad Superdex 75 26/60 PG column. Purified nanobodies were then dialyzed against buffer containing [20 mM sodium phosphate, 100 mM NaCl, pH 7.4] and stored at 4°C.

### Crystallization of Nanobodies and Cif:nanobody Complexes

All crystals were grown by vapor diffusion in either 200 nL sitting drops at 293 K or 4 µL hanging drops at 291 K. For all Cif:nanobody complexes, the complex was pre-formed by incubation of Cif or Cif-6xHis with the desired nanobody in a 1.25 molar excess and then purified by SEC using Superdex 200 resin. For Cif:VHH101 and Cif:VHH113, the secreted forms of VHH101 and VHH113 were used. Microseeding was used to obtain quality Cif:VHH219 crystals and was performed using a vibrissa from *Felis silvestris catus* to deposit seed crystals into pre-equilibrated 4 µL hanging drops composed of Cif:VHH219 and well solution. All crystals were cryo-protected prior to flash cooling as follows. Drops containing crystals were pre-conditioned with well solution supplemented with either glycerol or ethylene glycol (termed cryoprotectant) by first pipetting either 0.5 µL of cryoprotectant into 200 nL drops or 1 µL of cryoprotectant into 4 µL drops. Crystals were then transferred to a drop of cryoprotectant, allowed to equilibrate, then flash cooled in liquid nitrogen. A comprehensive description of protein concentrations, buffers, well solution compositions, drop ratios, and cryoprotectants can be found in Table S1.

### Structure Determination and Refinement

Rotation X-ray diffraction data were collected at 100 K at the National Synchrotron Light Source II on the AMX (17-ID-1) beamline using a Dectris Eiger 9M detector or the FMX (17-ID-2) beamline using a Dectris Eiger 16M detector. A total of 180° of data were collected on each crystal with a rotation of either 0.1° or 0.2° per frame. Data were reduced using XDS then scaled and merged with XSCALE.^51,52^ Crystals of the Cif:VHH101 complex grew as two fused crystals and produced low-quality data when processed as a single crystal. High-quality data were obtained by dividing the X-ray diffraction images into two segments to be processed as separate crystals and then merged into a final data set.

For all structures, the R_free_ set was assigned in thin resolution shells using 5% of total reflections using the reflection file editor in PHENIX.^53,54^ Initial phase estimates were obtained by molecular replacement using PHASER-MR.^55^ The publicly available structure of a VHH domain (pdb_00003ezj) was used as a search model for the crystal structure of VHH222 in its free form. A preliminary structure of the VHH113 domain determined in the Cif:VHH113 complex was used as a search model for the crystal structure of VHH113 in its free form. For all Cif:nanobody complexes, a single Cif protomer (pdb_00003kd2) was used as a search model for all Cif chains. The refined structure of VHH113 (pdb_00008e2n) was used as a search model for the VHH chains in the crystal structure of the Cif:VHH101 complex. A preliminary structure of VHH222 was used as a search model for the VHH chains in the crystal structures of the Cif:VHH113 and Cif:VHH219 complexes. The refined structure of VHH222 in its free form (pdb_00008e1c) was used as a search model for the VHH chains in the crystal structures of the Cif:VHH114 and Cif:VHH222 complexes.

Automated refinement was carried out using PHENIX.REFINE^53,54,56^ and manual refinements were performed in Coot.^57^ Anisotropic refinements of protein chains for the Cif:VHH219 complex and Cif:VHH101 complex were carried out with TLS tensors^58,59^ using one group per chain. Full anisotropic refinement of protein chains was performed during automated refinement of the structure of VHH113 in its free form. Torsion-angle NCS restraints were applied during automated refinement of the Cif:VHH101 complex. Positive density peaks were identified and evaluated for water placement using the programs PEAKMAX and WATPEAK from the CCP4 suite,^60,61^ respectively, followed by manual inspection of candidate waters. A 4 σ positive density peak in the Cif active site was present in Cif:VHH219 complex that could not be modeled. Atoms modeled with zero occupancy could not be confidently placed and were selected for zero-occupancy flagging after manual inspection of the 2m*F*_O_-D*F*_C_ map at a 0.5 σ cutoff.

A known artifact of recombinant cytosolic expression of nanobodies is the failure to form the canonical immunoglobulin disulfide bond due to the reducing environment of the cytoplasm.^62,63^ This is also the case for Cys23 and Cys104 in VHH219 (Figure S1). However, the canonical disulfide bond was observed in VHH222, generated using the same expression system, suggesting that disulfide maturation can sometimes overcome the redox hurdle. Regardless, while a reduced disulfide has been shown to decrease thermostability, it appears to have little to no impact on VHH binding affinity, function, or mechanical stability.^64,65^ Furthermore, we do not observe any appreciable differences in the positioning of main-chain atoms flanking either cysteine and therefore do not believe disulfide heterogeneity influences the conformation of either CDR1 or CDR3, which begin shortly after Cys23 and immediately following Cys104, respectively. We therefore observe no evidence that the aberrant sulfhydryl chemistry in recombinant VHH219 has any meaningful effect on the structural basis of the Cif:VHH219 interaction.

### Structural Analyses

All structural alignments were performed in PyMol^66^ using only main-chain atoms. For Cif:nanobody complexes, alignments were performed using main-chain atoms belonging to the Cif chains. Contact analyses were carried out using the CCP4 supported program CONTACT with a 5 Å cutoff. Shape correlation statistics for a single VHH domain bound to a Cif dimer was calculated using the CCP4 supported program SC.^67^ The solvent-accessible surface was calculated using the CCP4 supported program AREAIMOL^68,69^ with a solvent probe radius of 1.4 Å. For clarity, the binding interface was reported as one half of the total solvent-accessible surface area (SASA) sequestered in the complex as follows:

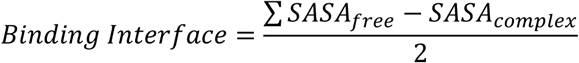

To measure the rotation angle between VHHs bound to Cif, complexes were first aligned by all main-chain atoms belonging to the Cif chain. The VHH chain to be moved was then aligned to the reference VHH chain by main-chain atoms of the framework regions. The resulting coordinates of the moved chain were used in conjunction with the starting coordinates to define the rotation axis and calculate the rotation angle using LSQKAB^70^ with the default settings. Spherical polar coordinates were used as a means to visualize the rotational displacement in PyMol.

Modeling of substrate-nanobody overlap was used to assess direct steric competitive inhibition. Least-squares alignment of all main-chain atoms of Cif was used to individually superimpose existing structures of Cif covalently linked to hydrolysis intermediates of three representative compact substrates (epibromohydrin, 1,2-epoxycyclohexane, and *S*-styrene oxide) (pdb_00004dnf, pdb_00005tnd, and pdb_00005tni, respectively), to the Cif:VHH219 complex. Next, a clash threshold equal to the sum of the van der Waals radii of each atom was used to evaluate the modeled interactions by measuring the shortest interatomic distance (D_min_) between the VHH219 Tyr111 side chain and the hydrolysis intermediate.

### Nanobody Sequence Numbering and Alignments

Nanobody sequences were numbered according to the IMGT unique numbering scheme for V-Domains^71^ using the IMGT/DomainGapAlign server^72^ (Table S2). Sequences were re-gapped as needed based on sequence and structural alignments using the crystal structures reported herein. Multiple sequence alignments of gapped sequences were visualized using ESPript 3.0^73^ with manual annotation. Collier de Perles diagrams were made using the IMGT/Collier-de-Perles web server.^74^

### Epoxide Hydrolysis Assays

A previously established assay for cyano(6-methoxynaphthalen-2-yl)methyl(oxiran-2-ylmethyl) (CMNGC) hydrolysis by Cif^23^ was adapted to test inhibition by α-Cif nanobodies. Prior to the start of the reaction, Cif was incubated with either buffer [20 mM sodium phosphate, 100 mM NaCl, pH 7.4], VHH, or Tiratricol (CAS ID: 51-24-1) at room temperature for 15 min. Reactions consisting of 1.2 µM Cif, 5.5 µM VHH (or buffer), 25 µM CMNGC, 15 µM thesit, and 1% (*v/v*) DMSO were incubated at 37°C for 15 min. Tiratricol was used in control reactions at concentrations of 25 µM, 12.5 µM, and 6.25 µM. The fluorescent product from CMNGC hydrolysis was measured using a TECAN Infinite M1000 plate reader set with an excitation wavelength = 330 nm and emission wavelength = 465 nm. Statistical analyses were performed in R^75^ and assessed by ANOVA using Dunnett’s post-hoc test.

An adrenochrome assay^76^ adapted to measure Cif hydrolysis of 14,15-epoxyeicosatrienoic acid (14,15-EET, CAS ID: 197508-62-6)^18^ was applied to the assessment of Cif inhibition by α-Cif nanobodies. The general reaction contained 20 µM Cif, 1 mM 14,15-EET, 2.5% (*v/v*) DMSO, and buffer [20 mM sodium phosphate, 100 mM NaCl, pH 7.4] and was incubated at 37°C for 30 minutes. Prior to the addition of 14,15-EET, reactions were treated with either buffer, VHH, or Tiratricol and incubated at room temperature for 15 minutes. VHH and Tiratricol were used at a final concentration of 25 µM. The assay was terminated by addition of a half volume of 90% (*v/v*) acetonitrile containing 2 mM sodium periodate and the reaction between periodate and diol was allowed to proceed at room temperature for 30 min. To develop the assay, an excess of epinephrine dissolved in hydrochloric acid was added and allowed to react with residual periodate at room temperature for 5 min. Precipitated protein was removed from the solution by centrifugation at 21,000 x *g* for 5 minutes at 4°C and the absorbance was measured at 490 nm using a TECAN Infinite M1000 spectrophotometer. Statistical analyses were performed in R and assessed by ANOVA using Dunnett’s *post-hoc* test.

### Biolayer Interferometry

Binding kinetics were measured by biolayer interferometry using a Forte Bio Octet system. All binding experiments were conducted in phosphate buffered saline (PBS) [137 mM NaCl, 2.7 mM KCl, 8 mM Na_2_HPO_4_, 2 mM KH_2_PO_4_] supplemented with 0.05% (*v/v*) Tween 20 and 1% (*w/v*) BSA (PBST + 1% BSA). Cif-6xHis was captured on Sartorius HIS1K biosensors until a response of approximately 0.8 nm was achieved. Biosensors with immobilized Cif-6xHis were then dipped into wells containing two-fold dilutions of VHH in PBST + 1% BSA ranging from 400 nM to 6.25 nM; association was allowed to proceed for 300 s, followed by dissociation in PBST + 1% BSA for another 300 s. Biosensors were regenerated by five cycles alternating between 10 mM glycine pH 1.7 and PBST + 1% BSA. All binding experiments were double referenced using a biosensor with immobilized Cif-6xHis exposed to PBST + 1% BSA in place of VHH and with independent reference biosensors that were not used to capture Cif-6xHis. Binding curves were fit using a 1:1 binding model using Forte Bio Data Analysis 7.0 software. *K*_D_ was calculated as the ratio of *k*_off_ to *k*_on_ and reported as the mean of four curves. For VHH101, VHH108, VHH113, and VHH222, the errors exceeded the mean values for the dissociation rates, due to extremely slow dissociation. Using the slowest *k*_off_ value that was confidently fit (1×10^−4^ s^−1^; VHH214) and the experimentally determined *k*_on_ values, we estimated an upper bound on the values of *K*_D_ for these nanobodies, as reported in Table 2 and Fig S11.

**Table 1.**
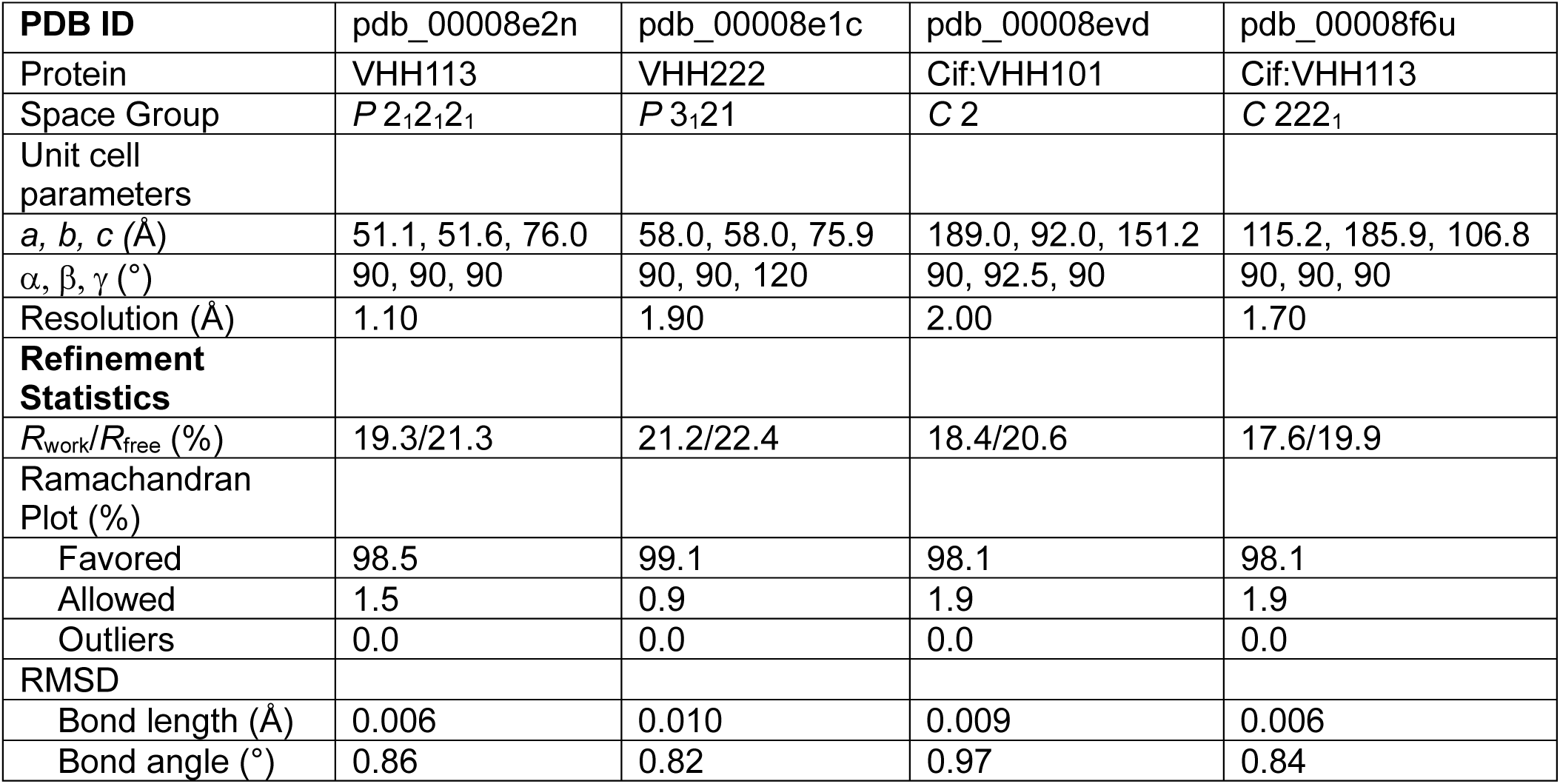
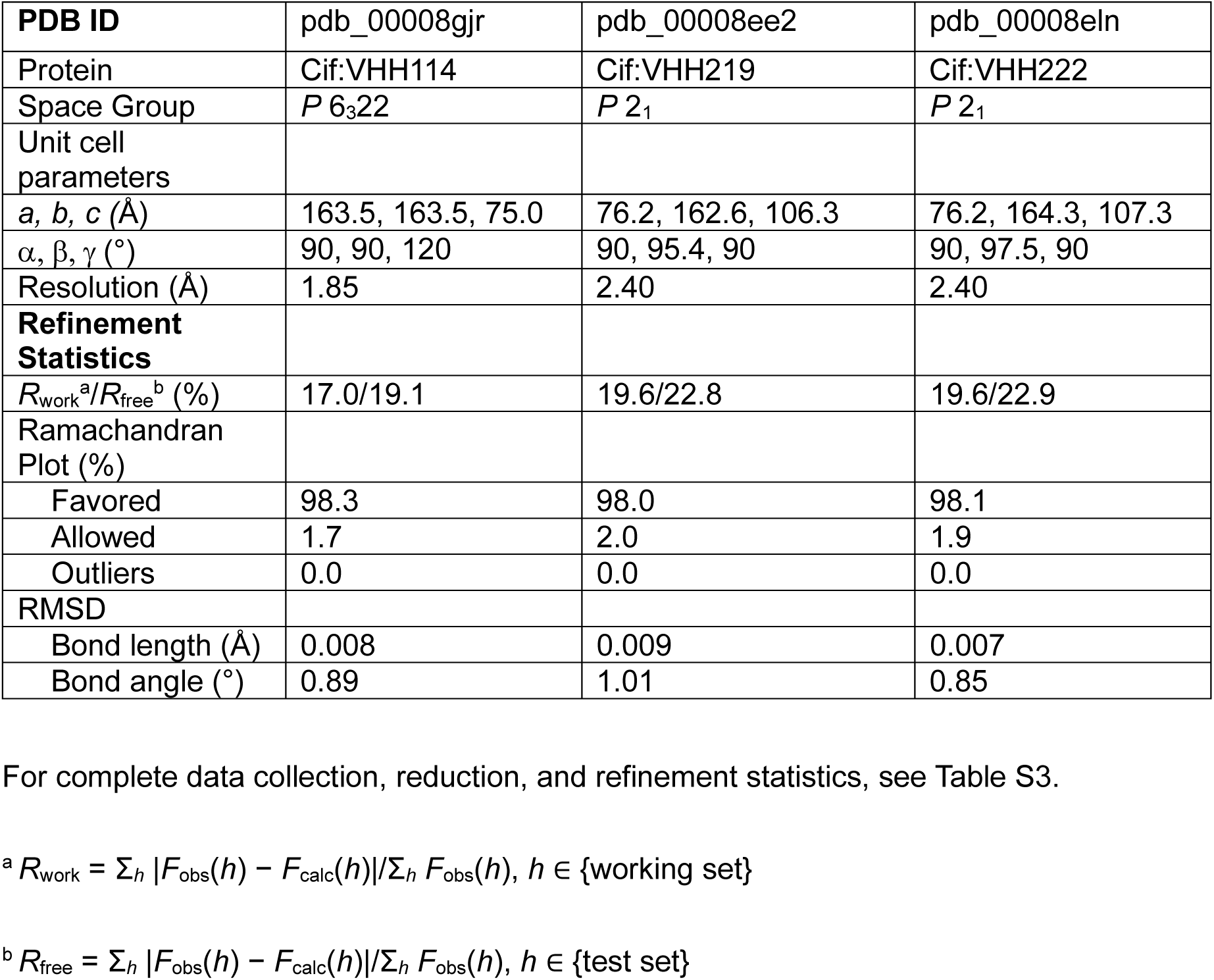
Summarized statistics for structures.

| **PDB ID** | pdb_00008e2n | pdb_00008e1c | pdb_00008evd | pdb_00008f6u |
| --- | --- | --- | --- | --- |
| Protein | VHH113 | VHH222 | Cif:VHH101 | Cif:VHH113 |
| Space Group | $P 2_1 2_1 2_1$ | $P 3_1 21$ | $C 2$ | $C 222_1$ |
| Unit cell parameters |  |  |  |  |
| $a, b, c$ (Å) | 51.1, 51.6, 76.0 | 58.0, 58.0, 75.9 | 189.0, 92.0, 151.2 | 115.2, 185.9, 106.8 |
| $\alpha, \beta, \gamma$ (°) | 90, 90, 90 | 90, 90, 120 | 90, 92.5, 90 | 90, 90, 90 |
| Resolution (Å) | 1.10 | 1.90 | 2.00 | 1.70 |
| **Refinement Statistics** |  |  |  |  |
| $R_{\text{work}}/R_{\text{free}}$ (%) | 19.3/21.3 | 21.2/22.4 | 18.4/20.6 | 17.6/19.9 |
| Ramachandran Plot (%) |  |  |  |  |
| Favored | 98.5 | 99.1 | 98.1 | 98.1 |
| Allowed | 1.5 | 0.9 | 1.9 | 1.9 |
| Outliers | 0.0 | 0.0 | 0.0 | 0.0 |
| **RMSD** |  |  |  |  |
| Bond length (Å) | 0.006 | 0.010 | 0.009 | 0.006 |
| Bond angle (°) | 0.86 | 0.82 | 0.97 | 0.84 |

|  |  |  |  |
| --- | --- | --- | --- |
| **PDB ID** | pdb_00008gjr | pdb_00008ee2 | pdb_00008eln |
| Protein | Cif:VHH114 | Cif:VHH219 | Cif:VHH222 |
| Space Group | $P 6_3 22$ | $P 2_1$ | $P 2_1$ |
| Unit cell parameters |  |  |  |
| $a, b, c$ (Å) | 163.5, 163.5, 75.0 | 76.2, 162.6, 106.3 | 76.2, 164.3, 107.3 |
| $\alpha, \beta, \gamma$ (°) | 90, 90, 120 | 90, 95.4, 90 | 90, 97.5, 90 |
| Resolution (Å) | 1.85 | 2.40 | 2.40 |
| **Refinement Statistics** |  |  |  |
| $R_{\text{work}}^a/R_{\text{free}}^b$ (%) | 17.0/19.1 | 19.6/22.8 | 19.6/22.9 |
| Ramachandran Plot (%) |  |  |  |
| Favored | 98.3 | 98.0 | 98.1 |
| Allowed | 1.7 | 2.0 | 1.9 |
| Outliers | 0.0 | 0.0 | 0.0 |
| RMSD |  |  |  |
| Bond length (Å) | 0.008 | 0.009 | 0.007 |
| Bond angle (°) | 0.89 | 1.01 | 0.85 |

For complete data collection, reduction, and refinement statistics, see Table S3.

**Table 2.** Binding constants of α-Cif nanobodies.

| Protein | $k_{\text{on}}$ ($\text{M}^{-1}\text{s}^{-1}$) | $k_{\text{off}}$ ($\text{s}^{-1}$) | $K_{\text{D}}$ (nM) |
| --- | --- | --- | --- |
| VHH101* | $5.0 \pm 0.68 \times 10^5$ | $4.7 \pm 5.9 \times 10^{-5}$ * | $< 0.2^*$ |
| VHH108* | $3.4 \pm 0.3 \times 10^5$ | $3.2 \pm 3.5 \times 10^{-5}$ * | $< 0.3^*$ |
| VHH113* | $5.5 \pm 0.58 \times 10^5$ | $1.5 \pm 2.2 \times 10^{-5}$ * | $< 0.2^*$ |
| VHH114 | $4.7 \pm 0.75 \times 10^5$ | $2.1 \pm 0.62 \times 10^{-4}$ | $0.44 \pm 0.084$ |
| VHH212 | $4.2 \pm 0.83 \times 10^5$ | $6.7 \pm 0.14 \times 10^{-4}$ | $1.6 \pm 0.19$ |
| VHH214 | $5.7 \pm 0.79 \times 10^5$ | $1.0 \pm 0.38 \times 10^{-4}$ | $0.17 \pm 0.051$ |
| VHH219 | $4.6 \pm 5.6 \times 10^5$ | $1.8 \pm 1.1 \times 10^{-4}$ | $0.39 \pm 0.21$ |
| VHH222* | $6.1 \pm 1.1 \times 10^5$ | $3.5 \pm 6.0 \times 10^{-5}$ * | $< 0.2^*$ |
| VHH314 | $4.1 \pm 0.76 \times 10^5$ | $8.7 \pm 1.4 \times 10^{-4}$ | $2.2 \pm 0.48$ |
| VHH320 | $3.8 \pm 0.54 \times 10^5$ | $3.7 \pm 2.6 \times 10^{-5}$ | $0.11 \pm 0.098$ |

Biolayer interferometry was used to measure binding kinetics of each nanobody to immobilized Cif-6xHis. Four nanobody concentrations (100 nM, 50 nM, 25 nM, and 12.5 nM) were used to calculate mean binding constants for each VHH. The $K_{\text{D}}$ was calculated as the $k_{\text{off}}/k_{\text{on}}$ ratio. For nanobodies marked with an asterisk (\*), the error in the measurement of the dissociation phase exceeded that of the mean and could not be confidently determined, due to extremely slow dissociation. Estimates for these nanobodies were calculated using the slowest $k_{\text{off}}$ value that could be fit with confidence ( $1.0 \times 10^{-4} \text{ s}^{-1}$ ; VHH214) as an upper bound.

### Data Visualization

All structural figures were rendered using PyMol as part of the SBGrid software environment.^77^ Graphs were generated using R with the following packages: tidyverse,^78^ plater,^79^ DescTools,^80^ ggsignif,^81^ ggprism,^82^ and ggplot2.^78^

### Accession Numbers

Coordinate files and structure factor files for all structures are available at the Protein Data Bank^83^ using the following codes: pdb_00008e1c (VHH222), pdb_00008e2n (VHH113), pdb_00008eln (Cif:VHH222), pdb_00008ee2 (Cif:VHH219), pdb_00008gjr (Cif:VHH114), pdb_00008evd (Cif:VHH101), and pdb_00008f6u (Cif:VHH113). X-ray diffraction image files were uploaded to the SBGrid Data Bank and are searchable by PDB codes. The dataset for Cif:VHH114 crystallized in conditions without citrate is available at the SBGrid Data Bank as SBGrid Dataset Number 1057, doi:10.15785/SBGRID/1057.

## Results and Discussion

### Structures of the Cif:VHH219 and Cif:VHH222 Complexes

To identify the molecular basis for nanobody recognition and inhibition of Cif and to explore the stereochemical determinants of our nanobody-displacement ELISA, we focused our initial efforts on VHH219 and VHH222.^24^ After co-crystallizing each nanobody with Cif, structures were determined for both complexes at 2.4 Å with excellent refinement statistics (Tables 1 and S3). In each case, the biological assembly contains one Cif homodimer and two VHH molecules in a 2:2 stoichiometry, *i.e*., one VHH associates with one Cif monomer and the second VHH associates with the second Cif monomer (Figs 1A and 1C). Both complexes crystallized with three biological assemblies in the asymmetric unit (ASU).

**Figure 1.**
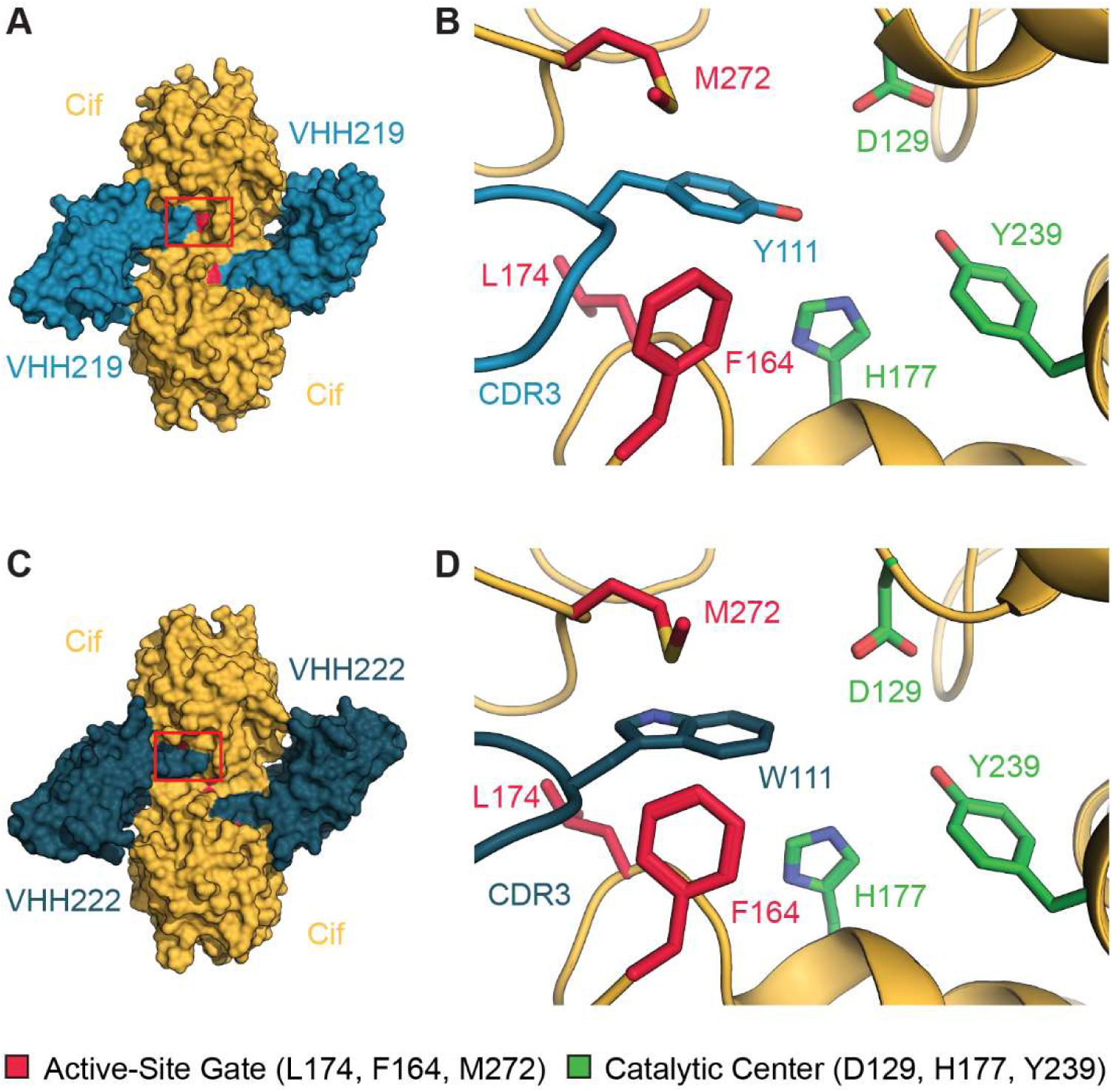
A shared equatorial binding mode occludes entry into the active site. Co-crystal structures of Cif with VHH219 and VHH222 reveal a 2:2 VHH:protomer binding stoichiometry and overlap of the VHH with a portion of the active-site gate. The VHHs interact primarily with the proximal Cif protomer and make some peripheral interactions with the distal protomer. The Cif:VHH219 (A) and Cif:VHH222 (C) complexes are illustrated as surface representations with Cif in orange, VHH219 in blue, and VHH222 in dark blue. Both VHHs bind near the dimer interface, and the red box highlights a portion of CDR3 covering the active-site entrance (red) of the upper protomer of Cif. The active-site entrance is occupied by Tyr111 of VHH219 CDR3 (B, blue sticks) and Trp111 of VHH222 CDR3 (D, dark blue sticks). Select residues in the catalytic center (nucleophile Asp129, ring-opening pair His177 and Tyr239) are shown as green sticks for orientation. Residues of the gate (Phe164, Leu174, and Met272) are shown as red sticks. Non-carbon atoms are colored by type: oxygen, red; nitrogen, blue; sulfur, yellow.

The interaction between Cif and each nanobody occurs near the Cif dimer interface (Figs 1A and 1C). After aligning all protomers (defined as a Cif monomer and its associated VHH molecule) in each ASU by all main-chain atoms in Cif, no differences exceeding maximum-likelihood coordinate error were observed except in a few loop regions of the nanobody that are distal to the binding face. As these regions do not contact Cif, we conclude that there are no meaningful differences at the binding interface among molecules related by non-crystallographic symmetry and have restricted all subsequent analyses to a single protomer unless otherwise stated. Similarly, as discussed in the *Methods* section, some heterogeneity in the formation of the canonical immunoglobulin disulfide bond was detected for different nanobodies (Fig S1). Consistent with studies of other nanobodies obtained by cytoplasmic bacterial expression,^84–86^ structural changes are localized to the immediate vicinity of the conserved cysteine residues and thus unlikely to affect the interaction with Cif. The conformations of Cif in the apo and nanobody-bound states were also assessed, and no structural changes greater than the estimated coordinate error were observed except in the gate-keeper residues, which are described in detail below.

### VHH219 and VHH222 Compete Sterically with Cif Substrates

Inhibitory nanobodies have been shown to operate through different mechanisms including direct steric competitive inhibition and allosteric modulation.^30,33,34,87–89^ As outlined below, this pair of co-crystal structures clearly demonstrate direct steric competition.

Each Cif monomer consists of a core domain together with a cap domain, which mediates dimerization and substrate selectivity.^90,91^ The active site is sandwiched between these domains and is only accessible by a tunnel whose entrance is regulated by the gate-keeping residues Phe164, Leu174, and Met272.^23,49^ In the apo state, these residues form a (presumably dynamic) barricade between the active site and bulk solvent. Each nanobody sits at the junction of the cap and core domains with significant overlap of the active-site gate by CDR3 (Figs 1A and 1C). Unexpectedly, each nanobody inserts an aromatic residue on CDR3 into the tunnel that leads to the catalytic center: either Tyr111 of VHH219 or Trp111 of VHH222 (blue and dark blue sticks, Figs 1B and 1D, respectively). In each case, the gate-keeper residues are forced open (Fig 2) assuming conformational states that resemble the open states seen in previous co-crystal structures of Cif bound to small-molecule inhibitors that extend through the gate.^23,44^

**Figure 2.**
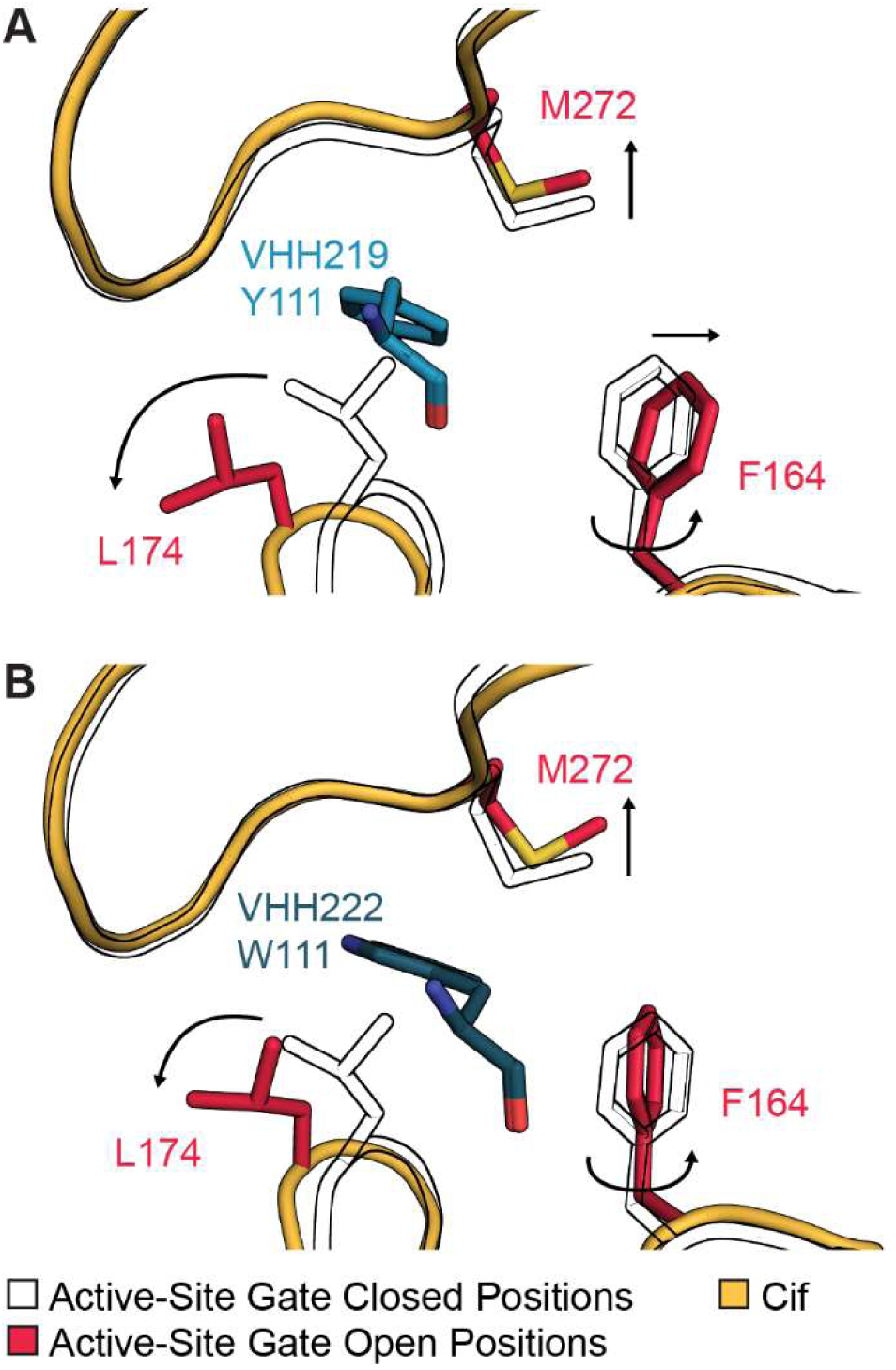
Active-site gate rearrangements are necessary for VHH binding. Cif:VHH co-crystal structures were aligned to apo Cif (pdb_00003kd2) by all main-chain atoms of a Cif protomer. A view into the pocket shows the active-site gate in the open conformation with VHH219 Tyr111 of CDR3 (A) and VHH222 Trp111 of CDR3 (B) inserted through the gate. Black arrows illustrate rearrangements in the gate residues from the (closed) Cif apo to the (open) Cif:VHH bound states. Side chains of the active-site gate are shown as black outlines (closed conformation) or red sticks (open conformation). This view is roughly orthogonal to that shown in Figure 1. The side chains of VHH219 and VHH222 are depicted as sticks colored blue and dark blue, respectively. Non-carbon atoms are colored by type: oxygen, red; nitrogen, blue; sulfur, yellow.

Nanobodies VHH219 and VHH222 were originally identified as inhibitors of Cif hydrolysis using CMNGC, a fluorogenic reporter,^23,24^ and we recapitulated these findings (Fig S2A). The structures also suggest that CDR3 intrudes upon the steric volume required for the docking of 14,15-EET, a known physiological substrate of Cif with implications for neutrophil recruitment and inflammatory resolution.^18,49^ Consistent with this observation, each nanobody prevents Cif hydrolysis of 14,15-EET (Fig S2B). To visualize the steric occlusion directly, each Cif:nanobody complex was superimposed on the previously reported crystal structure of Cif adducted to the 14,15-EET hydrolysis intermediate. Due to its extended geometry, the α-carboxylic acid of 14,15-EET protrudes through the gate creating an extensive and irreconcilable steric overlap with the bound position of CDR3 of VHH219 or VHH222 (Figs 3A and 3C). We therefore conclude that steric competition drives inhibition of 14,15-EET hydrolysis by VHH219 and VHH222.

**Figure 3.**
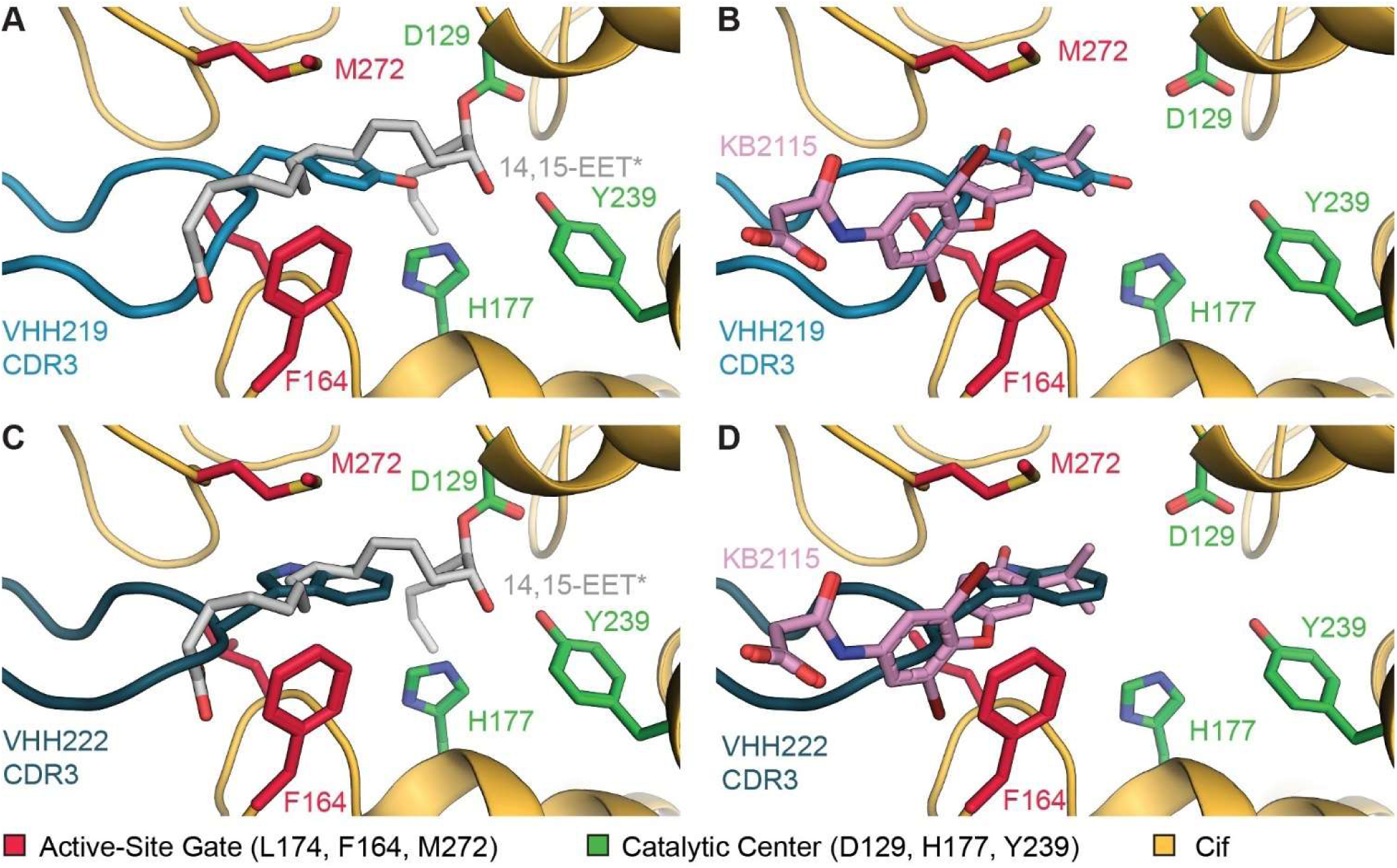
Structural evidence for inhibition by direct steric competition. To model interactions between nanobodies and 14,15-EET or inhibitors, Cif co-crystal structures were aligned by all main-chain atoms of a Cif monomer. These alignments strongly suggest sterically exclusive binding to Cif by VHH219, VHH222, a representative substrate (14,15-EET), and a representative inhibitor (KB2115). (A and C) Cif containing the adducted 14,15-EET hydrolysis intermediate (14,15-EET*, gray carbons) (pdb_00005jyc) aligned to Cif:VHH219 (blue) (A) or Cif:VHH222 (dark blue) (C). Both alignments show considerable overlap between CDR3 and the α-carboxylic acid of the 14,15-EET hydrolysis intermediate, suggestive of competitive inhibition. (B and D) Cif co-crystallized with inhibitor KB2115 (pink carbons) (pdb_00005hkb) aligned to Cif:VHH219 (blue) (B) and Cif:VHH222 (dark blue) (D). KB2115 overlaps with CDR3 of both VHHs. Colored as in previous figures with non-carbon atoms colored by type: oxygen, red; nitrogen, blue; sulfur, yellow.

Given the diversity of previously identified Cif substrates, we were curious if these findings could be generalized to much smaller xenobiotic substrates. Unlike 14,15-EET, these substrates can be fully contained within the active-site cavity and do not extend through the gate when adducted.^49,92^ Modeling confirmed that the shortest non-hydrogen interatomic substrate-nanobody distance (*D*_min_) is 1.8 Å or less, nearly half the minimum value necessary to avoid van der Waals collisions, for all substrates (Fig 4). In contrast with inhibition of 14,15-EET hydrolysis, which also would collide with main-chain atoms of CDR3 (Fig 3A), steric obstruction of small xenobiotic substrates is restricted to sequences with aromatic side chains, which make van der Waals contact with the attacking oxygen of the Asp129 nucleophile. A similar interaction would be mediated by Trp111 in VHH222. This further demonstrates the importance of the bulky hydrophobic side chains at position 111 in CDR3 in the exclusive occupancy of the Cif active site by either VHH219 or a substrate. Collectively, these data strongly suggest these nanobodies act as competitive inhibitors of Cif for all known substrates.

**Figure 4.**
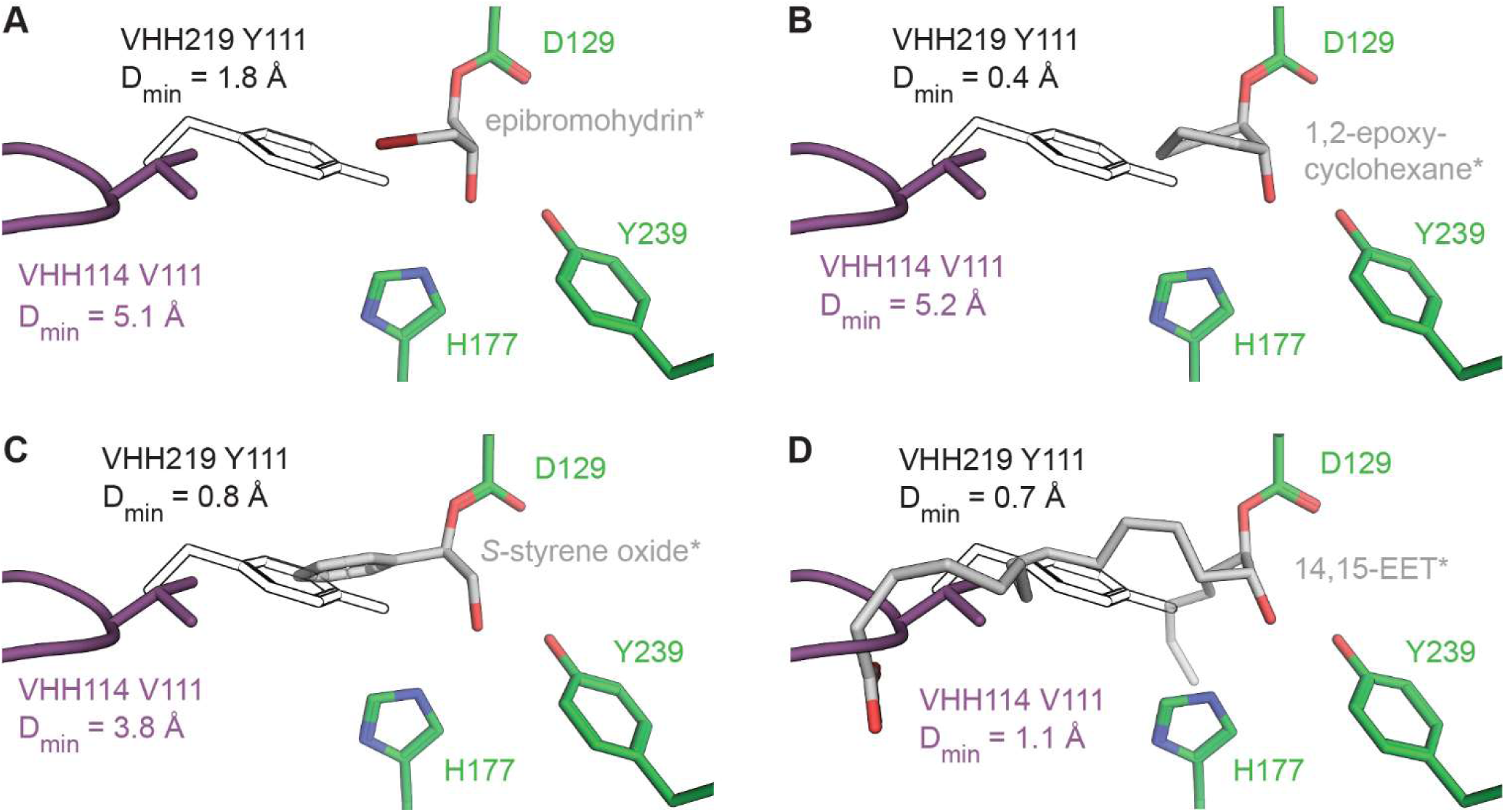
Nanobody inhibition involves variable degrees of steric overlap with different substrates. Potential clashes between the VHH114 or VHH219 CDR3s and a series of adducted substrates were evaluated for each Cif:VHH co-crystal structure and crystal structures of Cif covalently adducted to hydrolysis intermediates following alignments by main-chain atoms of the Cif chain. The shortest interatomic distance (D_min_) between VHH114 Val111 or VHH219 Tyr111 and the hydrolysis intermediate is shown. (A-C) Modeling suggests Cif can bind to VHH114 while maintaining van der Waals spacing with small substrates epibromohydrin (pdb_00004dnf) (A), 1,2-epoxycyclohexane (pdb_00005tnd) (B), and *S*-styrene oxide (pdb_00005tni) (C), consistent with non-competitive inhibition by VHH114, but in contrast to competitive inhibition by VHH219. (D) Overlap between 14,15-EET and VHH114 Val111/VHH219 Tyr111 is consistent with direct steric competitive inhibition for both nanobodies. For simplicity, only Cif residues from structures of hydrolysis intermediates are shown. The nucleophile (Asp129) and ring-opening pair (His177 and Tyr239) are shown as green sticks and the adducted hydrolysis intermediates (*) are shown as gray sticks. VHH114 CDR3 is shown in purple. For visual simplicity, only the side chain of Tyr111 of VHH219 CDR3 is shown and rendered as a black and white outline. Non-carbon atoms are colored by type: oxygen, red; nitrogen, blue.

### The CDR3 Hairpin in VHH222 is Dynamic in its Unbound State

We also solved the crystal structure of nanobody VHH222 at 1.9 Å to assess the conformational state of the CDRs prior to Cif binding. Following exclusion of a conformationally diverse loop in framework region (FR) 2, the remaining structure of VHH222 in the absence of Cif superimposed well to all VHH222 chains in the complex, with a root-mean square deviation (RMSD) not exceeding 0.75 Å (Fig S3A). Although the positions of CDR1 and CDR2 are equivalent in both states, a region of CDR3 spanning residues 111:113 could not be resolved in the unbound state, indicating potential conformational heterogeneity (Fig S3B). In contrast, this CDR3 region is stabilized at the Cif:VHH222 interface and forms an extended β-hairpin with a slightly distorted Type I reverse turn-like structure, with Trp111 at the *i*+1 position (Fig S3C). In the Cif:VHH219 complex, VHH219 CDR3 adopts an analogous structure with Tyr111 at position *i*+1. However, VHH219 includes a main-chain bulge following the turn to accommodate an additional CDR3 residue absent in VHH222 (Fig S4). The solvent-accessible surface area (SASA) of Trp111 was then calculated to gauge how effectively the CDR3 conformation positions the side chain to insert deeply into the active site. The result is similar to that of a free tryptophan side chain, confirming that the reverse turn maximizes the exposure of the *i*+1 side chain. Conversely, 98.4% of Trp111’s SASA becomes buried during complexation with Cif; this event accounts for 24.4% of the total buried interface. Side chains at this position within the turn are sterically unencumbered, allowing Trp111 of VHH222, and by extension, Tyr111 of VHH219, to engage the gating mechanism, which in turn blocks access to the active site like a cork plugging a bottle.

### Nanobodies and Inhibitors Also Bind in Overlapping Sites

To gain insight into the interplay between nanobodies, small-molecule inhibitors, and Cif, we leveraged the existing structure of Cif bound to the small-molecule inhibitor KB2115.^44^ Akin to the aromatic sidechains at position 111, KB2115 extends through the gating mechanism, although it is anchored within the active site, rather than on the external surface of Cif. Nevertheless, there is definitive overlap of the nanobody epitopes and the binding site of KB2115 in the vicinity of the gate. After superimposing Cif:VHH219 with Cif:KB2115, KB2115 and the reverse turn of CDR3 both clearly occupy the same region. Specifically, the isopropyl- or hydroxyl-substituted benzene rings of KB2115 overlap with Tyr111. An analogous observation is made with the Trp111 ring system in the Cif:VHH222 complex (Figs 3B and 3D). This explains the earlier observation that nanobodies VHH219 and VHH222 can be displaced from Cif by small-molecule competitive inhibitors that target the Cif active site, providing a structural explanation for the aforementioned nanobody-displacement ELISA.^24^

### Mapping the Shared Epitopes of VHH219 and VHH222

We next sought to establish a molecular basis for each Cif:nanobody interaction. While the paratope is generally convex, the extended CDR3 loops of VHH219 and VHH222 create protrusions that engage Cif. The resulting binding interfaces cover 1003 Å^2^ and 1081 Å^2^, with shape correlation statistics (Sc) of 0.67 and 0.71, respectively, comparable to other reported nanobody:antigen interfaces.^31,93^ Although the majority of the interactions occur between the VHH and the proximal Cif protomer, a few van der Waals contacts are observed between the VHH framework regions and the distal Cif protomer. However, interaction distances are variable and at times absent in other protomers in the ASU, leading us to conclude that these distal interactions are of a transient nature and may be influenced by lattice packing. The epitope analysis was therefore restricted to the proximal interactions using a 5 Å cutoff. This highlighted 35 Cif residues interacting with VHH219 and 39 residues interacting with VHH222. Strikingly, the nanobodies share 34 Cif contacts despite limited sequence conservation (46% identity) in the CDRs (Fig S5).

Subsequent analysis of the structures and sequences in tandem provided insight into how two different nanobodies interact with highly similar epitopes. After surveying the interactions at the Cif:nanobody proximal interface for both complexes, several shared contacts were identified (Fig 5) that map to conserved residues in the CDRs (Fig 6). CDR1 from each nanobody interacts with Cif His269 via two hydrogen bonds. The first involves the VHH main-chain carbonyl at position 35, while the second engages the hydroxyl of the Thr37 side chain. For both nanobodies, Arg58 of CDR2 extends toward the carbonyl oxygens of Cif Lys205 and Ala208. Several interactions between CDR3 and Cif likely contribute to the positioning of the inhibitory aromatic residue within the active-site gate. In both cases, Leu109 of CDR3 intercalates between Gln170 and Leu174 of Cif and is accompanied by two hydrogen bonds along the main chain leading into the reverse turn-like structure. At the corner of the turn is the defining hydrophobic interaction responsible for inhibition: VHH219 Tyr111/VHH222 Trp111 has been inserted through the gating mechanism.

**Figure 5.**
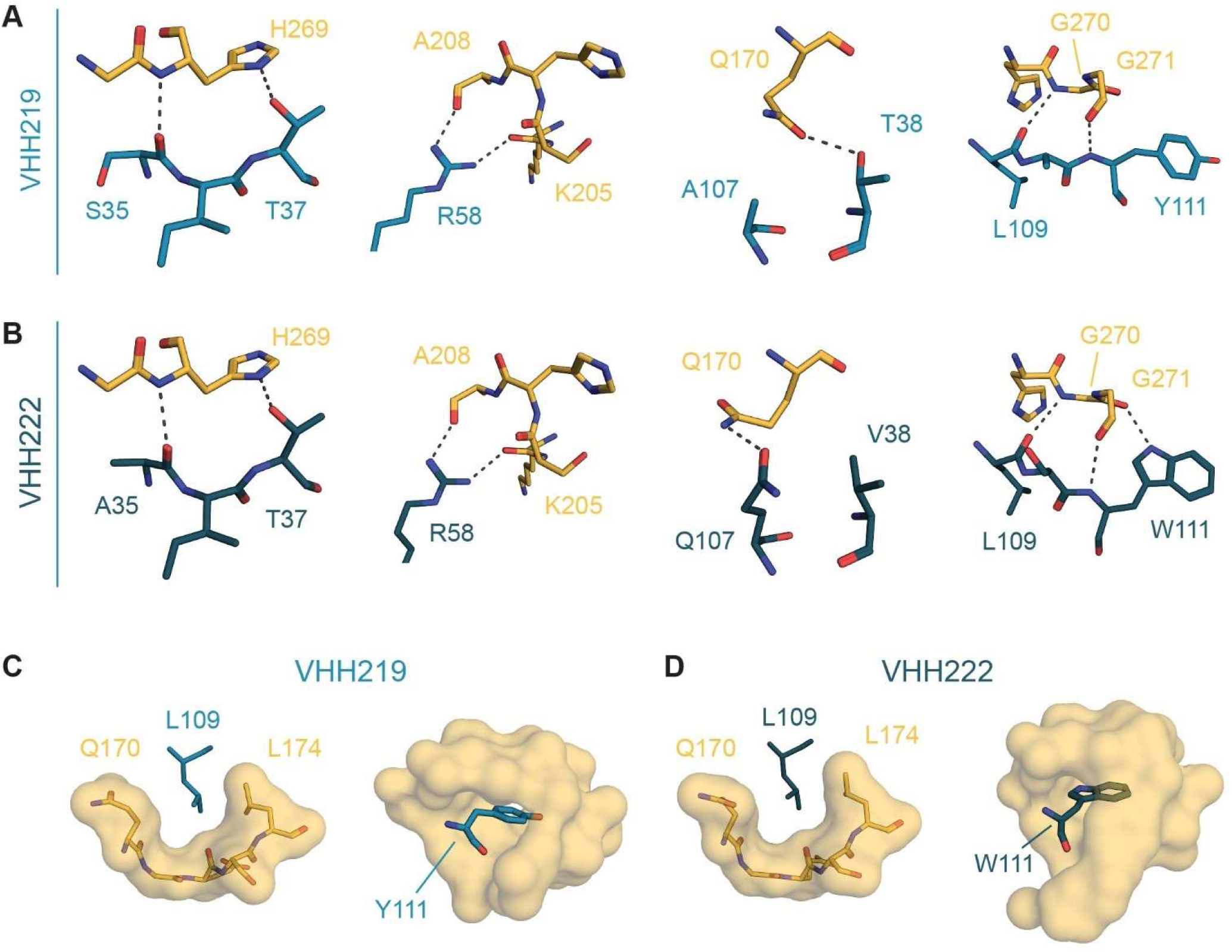
Cif:VHH interactions share striking similarity. Canonical stereochemical interactions are shown at the Cif:VHH219 and Cif:VHH222 interface. The Cif:VHH219 (A) and Cif:VHH222 complexes (B) are in part driven by near-identical hydrogen-bond interactions, with variation only observed with Cif Gln170 and an additional hydrogen bond between Cif and the indole ring of VHH222 Trp111. Key hydrophobic interactions between Cif and CDR3 of VHH219 (C) and VHH222 (D) are also shown. Leu109 intercalates between the gate-keeper residue Leu174 and aliphatic skeleton of Gln170, while the aromatic side chain of Tyr111/Trp111 is buried deep in the active-site entryway. Colors are shown as in previous figures with non-carbon atoms colored by type: oxygen, red; nitrogen, blue.

**Figure 6.**
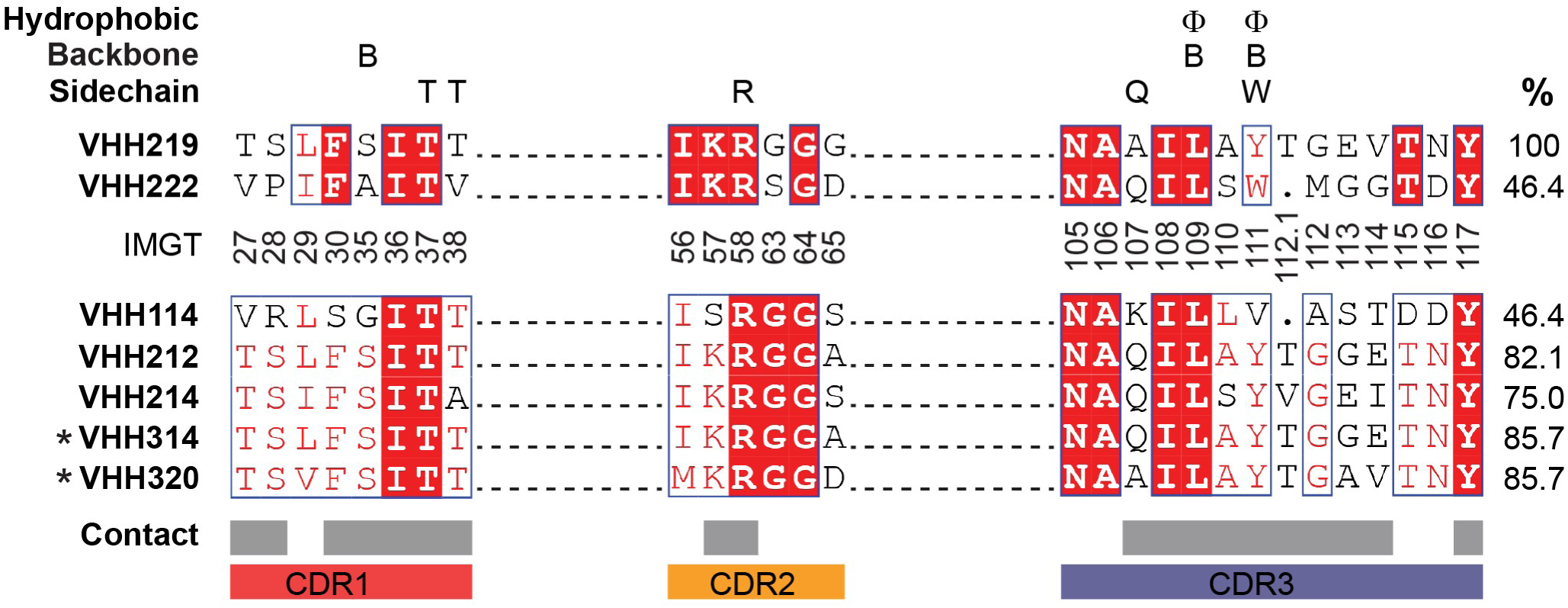
Cif:VHH contacts are preserved despite sequence variation. Pairwise alignment of VHH219 and VHH222 in the top two lines highlight conserved CDR residues. The interactions found at each Cif:VHH interface (Figure 5) were categorized and then mapped to the alignment. Categories are as follows: *Hydrophobic* indicates the burial of hydrophobic VHH side chains into specific pockets located on Cif; *Backbone* indicates hydrogen bonds involving main-chain atoms of residues in a VHH CDR; and *Sidechain* indicates hydrogen bonds involving side-chain atoms of residues in a VHH CDR. The column on the right displays sequence identity of CDR residues to those of VHH219. Beneath the alignment are the CDR positions as determined by IMGT unique numbering with gapped positions appearing as a dot in the alignment. Below the IMGT positions are alignments of additional VHHs with requisite residues to maintain these interactions. Asterisks (*) denote VHHs that did not show Cif inhibition during preliminary assessment ^24^. Gray bars indicate residues identified through surface contact analysis. CDRs are annotated by colored bars as follows: CDR1 – red, CDR2 – orange, and CDR3 – purple.

There are, however, some differences in the points of engagement that nonetheless stabilize highly congruent interactions. For example, VHH222 forms an additional hydrogen bond between the indole nitrogen of Trp111 and carbonyl oxygen of Cif Gly270 not seen with VHH219. A hydrogen bond is found between Cif Gln170 and VHH219 CDR1 residue Thr38 but is absent in the Cif:VHH222 complex due to a valine substation at the equivalent position. In VHH222, contact with Cif Gln170 instead involves a rotamer shift of Gln170 that brings it into contact with CDR3 Gln107. Considering these results, we conclude that antigen recognition by VHH219 and VHH222 occurs through related interactions in nearly identical, overlapping epitopes dominated by a shared subset of Cif residues.

### A Sequence Motif Encodes a Shared Structural Basis for CDR3 Inhibition of Cif

Having uncovered the stereochemical basis for Cif inhibition by VHH219 and VHH222, these findings were extrapolated to identify similar nanobodies within the Cif-specific panel and to highlight additional sequences for further study. We first selected candidate nanobodies with CDR sequences containing amino acids at positions that are compatible with the stereochemical attachment points seen in the Cif epitope outlined in Fig 5. A secondary filter was then applied to further sort sequences from the panel that contain a tryptophan or tyrosine at position 111. This strategy identified four additional inhibitory nanobodies (VHH212, VHH214, VHH314, and VHH320, Figs 6 and S6) that may act through a similar mechanism as described above. A fifth nanobody (VHH114) was also identified that satisfies most of the motif characteristics with a prominent exception: it has a small valine side chain at position 111, in place of a larger aromatic residue.

All five candidates were assayed. Three of the predicted nanobody sequences (including VHH114) had previously been flagged as Cif inhibitors and were confirmed on retest. In addition, the structure-guided sequence analysis identified two nanobodies (VHH314 and VHH320, Fig 6) that had not shown inhibitory activity in the earlier screen.^24^ These nanobodies were re-evaluated using two different biochemical assays and were shown to inhibit Cif with high efficacy (Fig S2). These results led us to reclassify VHH314 and VHH320 as inhibitory nanobodies and corroborated the predictive value of the sequence motif analysis.

### An Aromatic “Stopper” is not Required for Inhibition

While the degree of steric overlap would likely be reduced, we hypothesized that VHH114 would recognize the same core epitope and that CDR3 of VHH114 would adopt a similar Type I reverse turn-like structure with Val111 at position *i*+1. Crystallization of the Cif:VHH114 complex led to high-quality diffraction data in spacegroup *P* 6_3_22; structure refinement was straightforward (Tables 1 and S3). A shift in the Cif main chain-spanning residues 183 to 186 was observed, but is influenced by two nearby citrate molecules and is unlikely to have biological significance based on comparison with electron density from a citrate-free data set of comparable resolution (SBGrid Data Bank: dataset 1057) (Fig S7). As with VHH219, the VHH114 canonical disulfide bond is reduced (Fig S8).

The Cif:VHH114 co-crystal structure confirmed our predictions: it is very similar to those of Cif:VHH219 and Cif:VHH222. With a binding interface of 873 Å^2^ and an S_c_ value of 0.71, VHH114 recognizes a smaller area of Cif than VHH219 and VHH222, consistent with the smaller buried side chain, but nonetheless footprints to approximately the same region (Fig S9a). We also confirmed that key interactions identified in the Cif:VHH219 and Cif:VHH222 complexes are also present at the Cif:VHH114 interface (Fig S10). Importantly, VHH114 CDR3 adopts an extended β-hairpin with Val111 at the analogous *i*+1 position and reaches towards the Cif gating mechanism, which is sterically blocked and held in the open conformation (Fig S9b).

While the stubby side chain of Val111 does not reach as deeply into the active site as the corresponding sidechains in VHH219 and VHH222, VHH114 still sterically overlaps with 14,15-EET, which extends through the gate and would collide with Val111 as well as main-chain atoms of CDR3 (Fig 4D). In contrast, Val111 does not sterically overlap with small substrates. As for the aromatic side chains in VHH219 and VHH222, the interactions were modeled to determine D_min_ for VHH114 and the same set of substrates (Fig 4). Epibromohydrin and 1,2-epoxycyclohexane both have a D_min_ exceeding 5 Å, which should easily allow simultaneous binding of either substrate with VHH114. For *S*-styrene oxide, the D_min_ is 3.8 Å, which is greater than the threshold for carbon-to-carbon clashing. Modeling of the 14,15-EET interaction shows significant overlap with main-chain atoms of VHH114 CDR3 (Fig 4D). We therefore propose that the nanobodies possessing a Tyr or Trp at CDR3 position 111 (VHH212, VHH214, VHH219, VHH222, VHH314, and VHH320) act as obligate competitive inhibitors, while VHH114 may conditionally operate as a competitive or non-competitive inhibitor depending on substrate size.

### Inhibition by Nanobodies of Different Sequence

So far, all of the inhibitory nanobodies presented use a conserved set of binding interactions and operate through a mechanism analogous to that of VHH219 and VHH222, reflected in a shared sequence motif. The analysis implemented above thus explains the mechanism behind the majority of inhibitory nanobodies in the Cif-specific library. However, the sequences of three nanobodies originally flagged as candidate inhibitors (VHH101, VHH108, and VHH113) do not conform to the paratope motifs defined above and also share little overall sequence identity with VHHs described thus far (Fig S6). All three nanobodies demonstrate at least low nanomolar binding (Table 2, Fig S11). However, even at a concentration experimentally confirmed to saturate binding, VHH108 fails to demonstrate an inhibitory effect in our standard Cif hydrolysis assays. However, we were able to recapitulate potent inhibition by VHH101 and VHH113 (Fig S2), confirming their original characterization.^24^

### A CDR2-Based Alternative Paratope

While VHH113 and VHH101 share nearly identical CDR2 and CDR3 sequences with each other (89% identity), they have distinct CDR1 sequences (38% identity, Fig S12A). As such, both were selected for structural studies in complex with Cif to capture any important differences in inhibition strategies. Crystals yielding high-resolution diffraction data were obtained for both complexes, and the resulting structures exhibit comparable resolution, refinement statistics, and biological assemblies (Tables 1 and S3). For each structure, no meaningful differences were noted between Cif and the associated CDRs for all protomers within each ASU, as main-chain RMSDs are at or below estimated maximum-likelihood coordinate error.

A detailed analysis of the Cif:VHH113 and Cif:VHH101 interfaces was performed to assess the impact of the different CDR1 sequences on Cif binding. The complexes share similar quaternary structures, in which VHH113 and VHH101 each bind near the dimer interface and overlap with the active site of the proximal Cif protomer (Figs 7A and S13A). VHH113 recognizes a binding interface of 1077 Å^2^, with an S_c_ value of 0.65, while VHH101 forms an interface of 1062 Å^2^, with an S_c_ value of 0.68, consistent with other nanobodies, including those described above.

**Figure 7.**
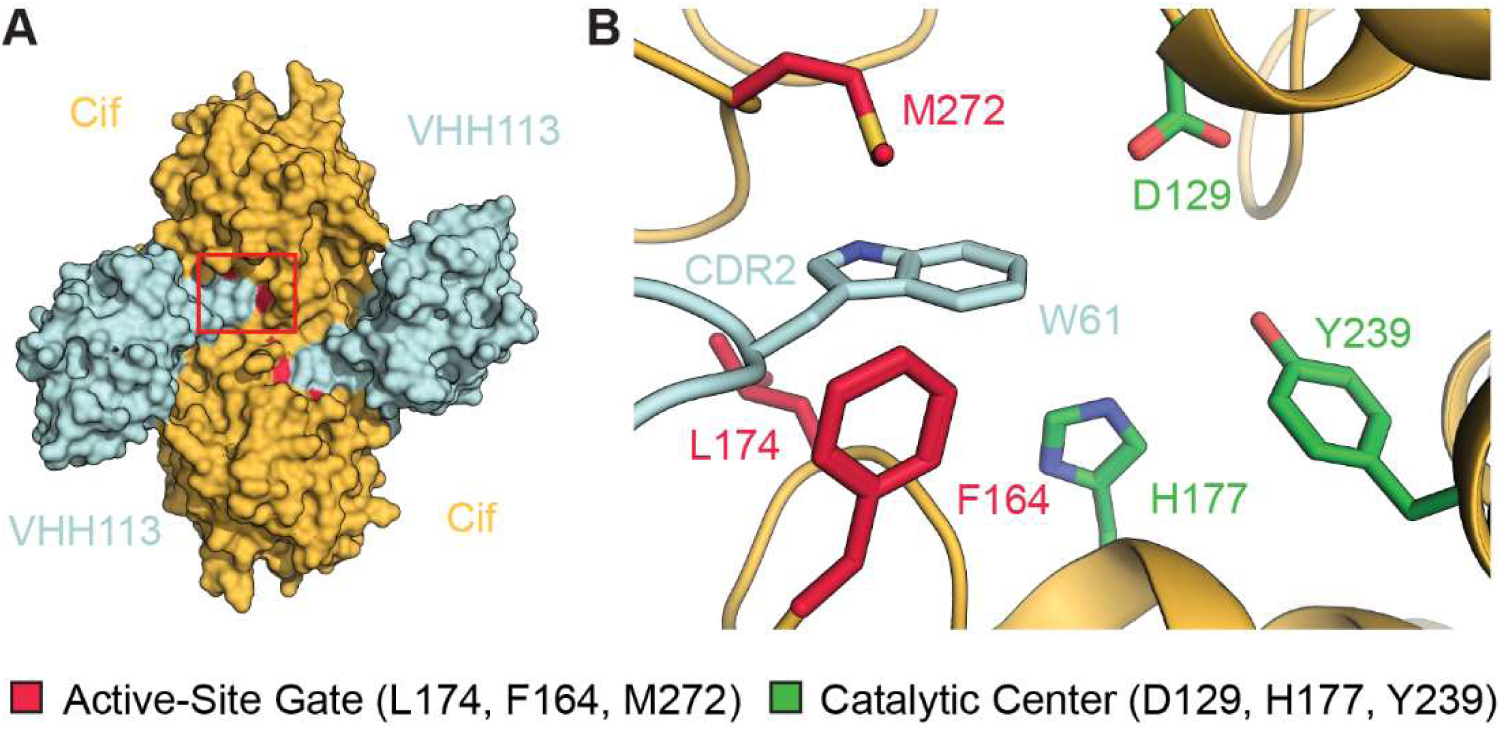
VHH113 CDR2 recapitulates the mechanism of competitive inhibition by direct steric occlusion seen with CDR3 inhibitors. The quaternary structure of the Cif:VHH113 complex (A) closely resembles that of Cif:VHH219 and Cif:VHH222 (Figures1A and 1C, respectively). Furthermore, VHH113 implements an analogous mechanism of inhibition (B). (A) The Cif:VHH113 complex is illustrated as surface representations, with Cif in orange and VHH113 in faded denim. The red box highlights a portion of CDR2 covering the active-site entrance of the upper Cif protomer (red). (B) Detailed view of the region within the red box at a slightly different angle. Residues of the active-site gate (Phe164, Leu174, and Met272) are depicted as red sticks and found in the open conformation, with VHH113 Trp61 of CDR2 inserted through the gating mechanism. Select residues in the catalytic center (nucleophile Asp129, ring-opening pair His177 and Tyr239) are shown as green sticks for orientation. Non-carbon atoms are colored by type: oxygen, red; nitrogen, blue; sulfur, yellow.

In addition to the CDR2 interactions, which are described in more detail below (Figs 8A and S14A), numerous hydrogen-bond interactions are common to both of the Cif:VHH113 and Cif:VHH101 complexes (Figs 8B and S14B). The Cif:VHH113 complex has two additional hydrogen bonds between Cif and CDR1 not found in the Cif:VHH101 complex: one between VHH113 Arg28 and Cif Gly294 and the other between VHH113 Glu27 and Cif Thr211 (Fig 8C). This difference arises from the lack of sequence conservation and structural homology between these CDR1 loops (Fig S12) which may reflect the expanded regions of hypervariability at the FR1-CDR1 junction in germline VHH alleles.^94^

**Figure 8.**
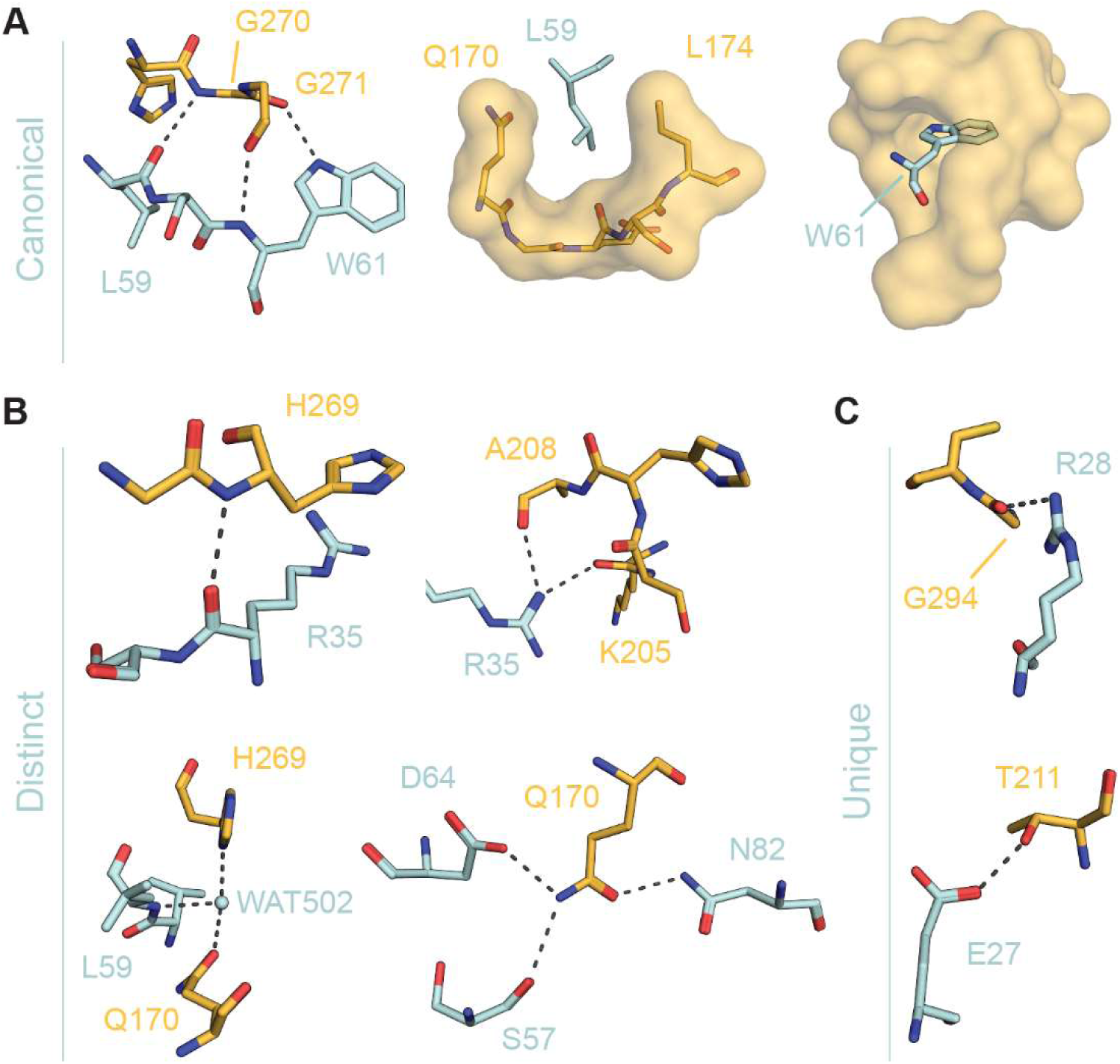
VHH113 is an alternative solution to conceptually identical means of Cif inhibition. VHH113 and VHH222 both inhibit Cif through direct steric occlusion of the active-site gate by a tryptophan side chain, but the Cif:VHH113 interface is composed of a mixture of canonical, distinct, and unique interactions. (A) The inhibitory VHH113 CDR2 forms interactions with Cif analogous to those found between Cif and the inhibitory VHH222 CDR3 (see last panel in Figure 5B as well as Figure 5D). (B) A cluster of interactions between VHH113 and Cif are distinct and not present in the Cif:VHH222 complex but are found in the Cif:VHH101 complex (see Figure S14). (C) VHH113 Interactions with Cif that are absent in the Cif:VHH101 complex (see Figure S14) TOP – A VHH113 interaction that recapitulates an interaction found at the Cif:VHH114 interface (see Figure S10B). BOTTOM – a unique VHH113 interaction. VHH113, faded denim; Cif, orange. Non-carbon atoms are colored by type: oxygen, red; nitrogen, blue.

Contrary to most published nanobody:antigen complexes, as well as those described above, in both VHH113 and VHH101 the CDR3 loops make minimal contact with Cif and do not directly contribute to inhibition. Instead, CDR2 constitutes the bulk of the paratope. Importantly, the CDR2 loops of both VHH113 and VHH101 recapitulate the previously described corking action by inserting a tryptophan side chain through the active-site gate (Figs 7B and S13B). VHH113 and VHH101 have nearly identical CDR2 sequences (Fig S12A and S15) and exert identical mechanisms of inhibition. Thus, these two inhibitory nanobodies represent a distinct class of CDR2-based Cif inhibitors.

### A CDR2 Reverse Turn Mimics Cif Inhibition by CDR3

To streamline comparisons between nanobody classes, the Cif-bound complexes with either VHH222 or VHH113 were selected as representatives of CDR3 inhibitors and CDR2 inhibitors, respectively. This pair was chosen because both nanobodies insert a tryptophan side chain through the gating mechanism and both reflect shared characteristics of homologous nanobodies.

Overall, the CDR sequences of VHH113 bear no meaningful similarity to those of VHH222 (Fig S16), which is reflected in the nanobody structures (Figs 9A and 9B). Given these differences, we were surprised to observe that the co-crystal structures of Cif:VHH113 bore a striking overall resemblance to Cif:VHH222 (compare Figs 1C and 7A). In particular, beyond blocking the gate (compare Figs 1D and 7B), the interactions between Cif and VHH113 CDR2 closely mirror those found between Cif and VHH222 CDR3 (compare Figs 8A, 5B – right panel, and 5D). VHH113 CDR2 and VHH222 CDR3 both form Type 1 reverse turn-like structures, and a sequence alignment of VHH113 CDR2 to VHH222 CDR3 highlights a key inhibitory motif of five amino acids. This pentad at the top of the loop shows close superposition of main-chain atoms following alignment of each complex by main-chain atoms of Cif (RMSD = 0.6 Å) (Figs 9C, 9D and S16).

**Figure 9.**
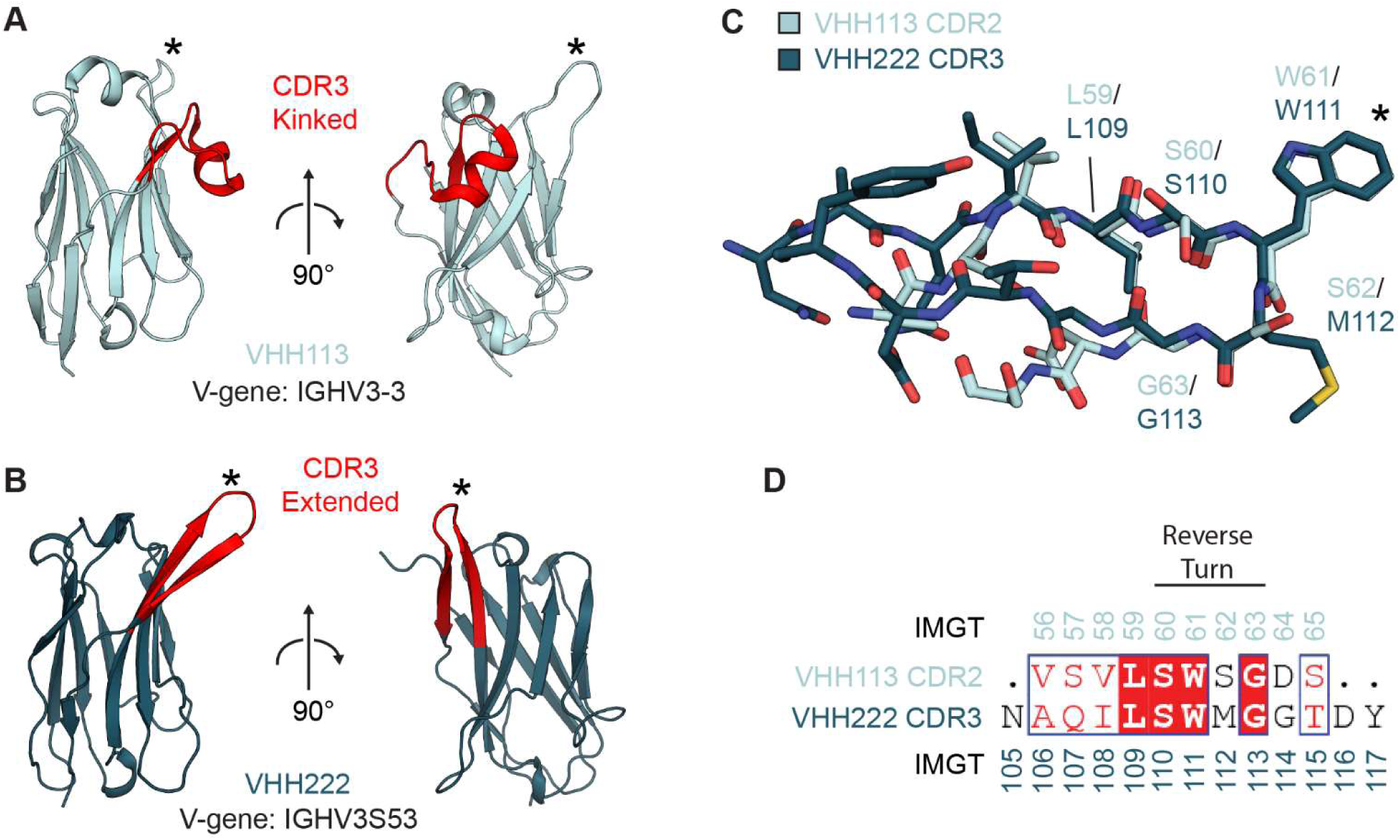
Structural mimicry of reverse turns in different CDR loops engenders a convergent mechanism of Cif inhibition by VHHs originating from different V-gene alleles. Steric occlusion of the Cif active-site entrance by a tryptophan in a reverse-turn like structure by both VHH113 and VHH222 is enabled by structural mimicry and sequence convergence in CDR2 and CDR3, respectively. Two viewing angles of VHH113 (faded-denim) (A) and VHH222 (dark blue) (B) are shown with CDR3 highlighted in red. The approximate location of the tryptophan side chain involved in blocking the active-site entrance is marked with an asterisk (*). Each is labeled with both the V-gene allele identified by the IMGT/BlastSearch tool and the CDR3 conformation. (C) Alignment of each co-crystal structure by main-chain atoms of the Cif chain shows the regions spanning the reverse turn-like structure of the inhibitory CDR superimpose well, RMSD = 0.6 Å. However, the path of the main chain diverges as it is followed away from the reverse turn back to the framework regions. The tryptophan that plugs the active-site entrance is marked with an asterisk (*). Non-carbon atoms are colored by type: oxygen, red; nitrogen, blue; sulfur, yellow. (D) A pairwise sequence alignment of VHH113 CDR2 and VHH222 CDR3 highlights a short region of sequence similarity corresponding to the reverse turn and the preceding leucine.

However, outside of the short segment of sequence identity within the Type 1 reverse turn, the structural alignment diverges as the β-strands veer to rejoin the framework regions of their respective core immunoglobulin domains (Fig 9C). While the VHH222 CDR3 forms an archetypal anti-parallel β-sheet along its length, the β-sheet of the VHH113 CDR2 is instead redirected by a kink and accompanied by a shift in main-chain hydrogen-bonding register (Fig 10). Owing to main-chain phi and psi angles incommensurate with β-strand hydrogen bonding, Ser57 is rotated so the side-chain hydroxyl moiety assumes the approximate position of the carbonyl in a typical β-sheet and serves as a hydrogen-bond acceptor to the amide group of Ser65 on the adjacent strand. This same hydroxyl functions as a donor in a second hydrogen bond with the carbonyl of the adjacent Val58 residue (Fig 10A). It is worth noting that CDR2 of VHH113 is rather peculiar due to its extended length as well as its unique conformation.^29,95^ Neither is represented in the canonical H2-13 (heavy chain CDR2, length 13) loop clusters^†^,^96,97^ although despite low (40%) sequence identity, a similar kinked conformation with highly similar hydrogen-bond pattern and register shift has been observed in an unrelated alpaca VHH in the H2-14 (heavy chain CDR2, length 14) loop cluster (Fig S17).^98^

**Figure 10.**
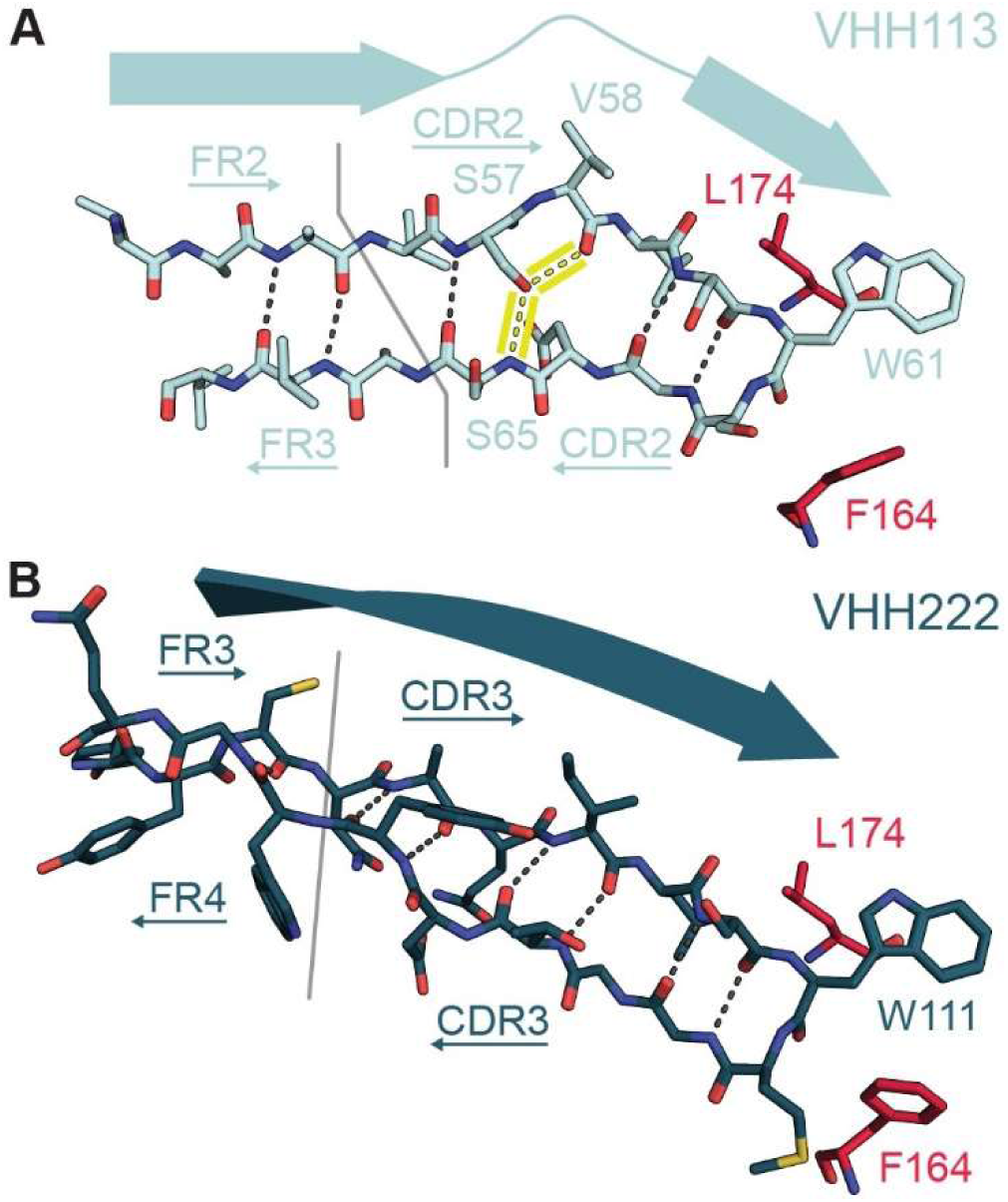
Conformations of CDRs implicated in Cif inhibition. The secondary structure of VHH113 CDR2 is kinked compared to VHH222 CDR3 in order to perform analogous functions. The loops of both CDRs form Type I reverse turn-like structures. Hydrogen bonds are depicted as gray dashes. Active-site gate resides Phe164 and Leu174 are shown for orientation (red sticks). (A) VHH113 CDR2 forms a kinked β-strand containing an atypical hydrogen-bond pattern highlighted in yellow. (B) VHH222 CDR3 resembles a common β-strand. VHH113, faded denim; VHH222, dark blue. Non-carbon atoms are colored by type: oxygen, red; nitrogen, blue; sulfur, yellow.

To evaluate if this feature is hard-wired by the CDR2 sequence or a possible result of induced fit during antigen binding, we solved the structure of VHH113 in its free state. While in a slightly twisted conformation, with a main-chain RMSD of 2.9 Å (Fig S18A and S18B) and a possible influence of lattice packing (Fig S19), the structural organization of CDR2 remains similar to the Cif-bound complex, with an atypical hydrogen-bonding pattern, kinked architecture, and general Type I reverse turn character (Fig S18C).

### An Axial Rotation Relates Bound CDR2- and CDR3-based Nanobodies

We next investigated how both classes were able to recapitulate highly similar mechanisms of inhibition given the different locations of the inhibitory CDR in the nanobody sequences (Figs S4, S15, and S16). Consistent with their shared immunoglobulin fold, the main-chain atoms of the VHH framework regions by themselves can be fitted to one another with a main-chain RMSD of 0.8 Å. If instead the Cif domains in each complex are superimposed, the main-chain RMSD between the same VHH atomic pairs rises to 13.0 Å. This is because each Cif-bound nanobody pivots by 89.8° around a rotation axis near CDR1. The resulting rotation places VHH113 CDR2 in a similar position with respect to the Cif active-site gate as VHH222 CDR3, accounting for the localized superposition of reverse-turn mimics (Fig 11). Put in other words, if one were to rotate VHH113 about this axis using its Cif-bound state as a starting orientation, the positioning of its CDRs would resemble the positions of the corresponding CDRs in VHH222. Because CDR1 lies near the rotation axis, it approximately rotates in place and thus localizes to similar footprints in each complex. This results in an alternate orientation of the α-helix of CDR1.

**Figure 11.**
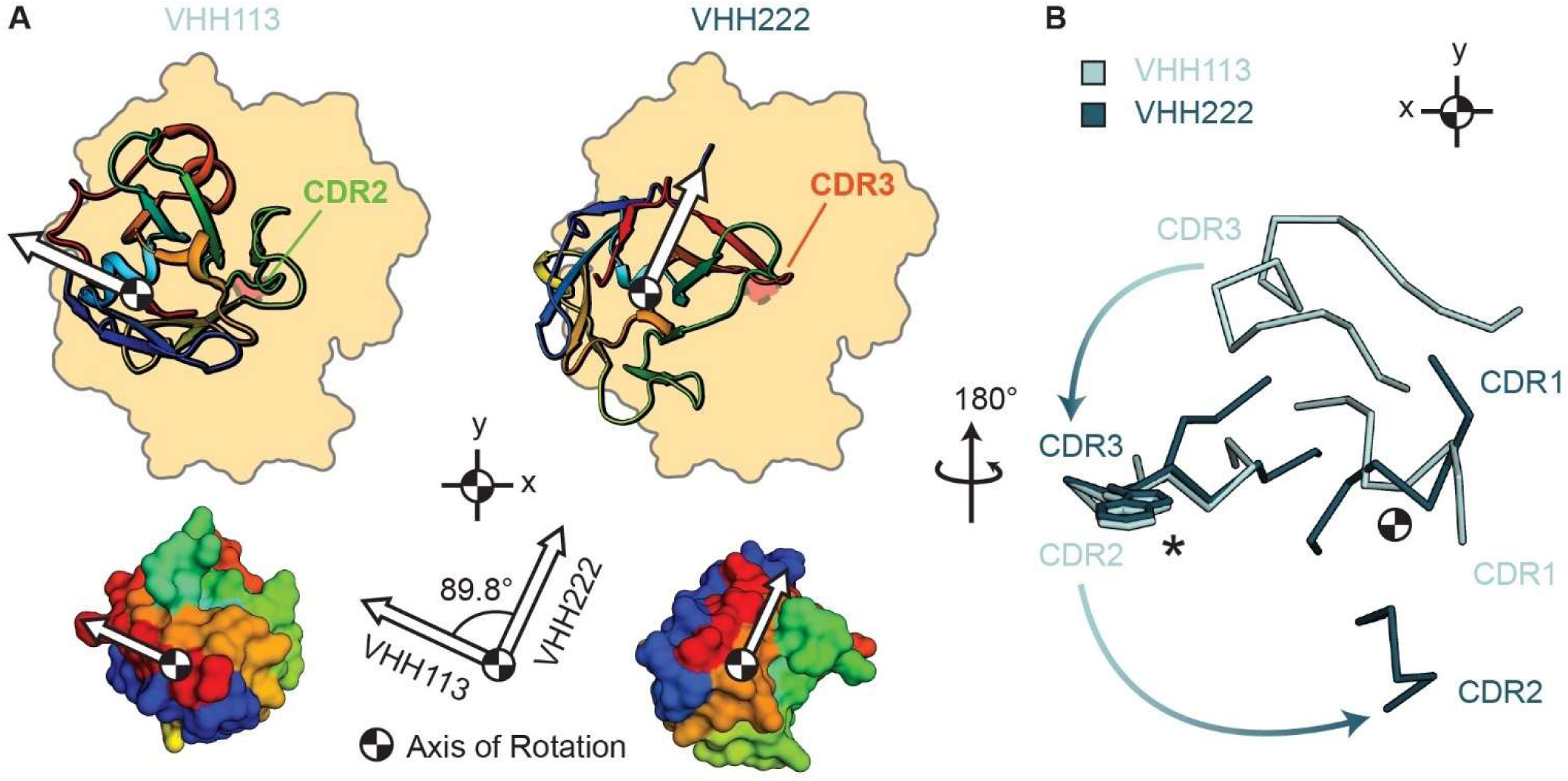
Different VHH binding orientations align different CDRs over the Cif active-site entrance. VHH113 and VHH222 footprint to similar regions of Cif but in different orientations. (A - TOP) Schematic showing relative rotation of each VHH viewed along the rotation axis (black and white pinwheel) from the back of the VHH looking down onto the epitope on Cif. VHH113 is rotated 89.8° along this rotation axis relative to VHH222. VHHs are colored rainbow and the CDR involved in inhibition is labeled. For simplicity, Cif is shown as an orange-filled outline with the active-site cavity shown in red. An arbitrary white orientation vector extends from the rotation axis and acts as a visual aid to help track rotation. (A - BOTTOM) Surface representations of each VHH colored and oriented as above to serve as an alternate depiction of the domain rotation. The angle of rotation is shown between the orientation vectors in the diagram between the rainbow-colored surface models. (B) The spatial organization of the CDRs with respect to Cif changes as a result of VHHs binding to Cif in different orientations as viewed from Cif looking “out” toward the VHH paratopes (view in A rotated 180° around the y-axis). Rotation of the immunoglobulin domain alters the spatial organization of CDRs relative to one another and accounts for superposition of the VHH113 CDR2 and VHH222 CDR3 reverse turns. The tryptophan side chain that inserts through the gate is shown in sticks and marked with an asterisk (*). VHH113, faded denim; VHH222, dark blue.

### Shared Stereochemical Attachment Points are Used by Both Nanobody Classes

In consideration of the sharply different spatial positions of CDRs within the paratope due to the domain rotation, we hypothesized that the respective classes might recognize distinct stereochemical handholds within their overlapping epitope footprints, and that outside of the active-site gate and substrate-entry tunnel, they might be comprised of largely disjoint sets of Cif residues. However, this was not the case. Instead, the majority of Cif contact residues participate in each Cif:VHH pairing. This “core epitope” was found to encompass between 73% - 86% of the total epitope for each co-crystal structure and is extensively utilized by both CDR3-centric and CDR2-centric inhibitory nanobodies (Fig 12).

**Figure 12.**
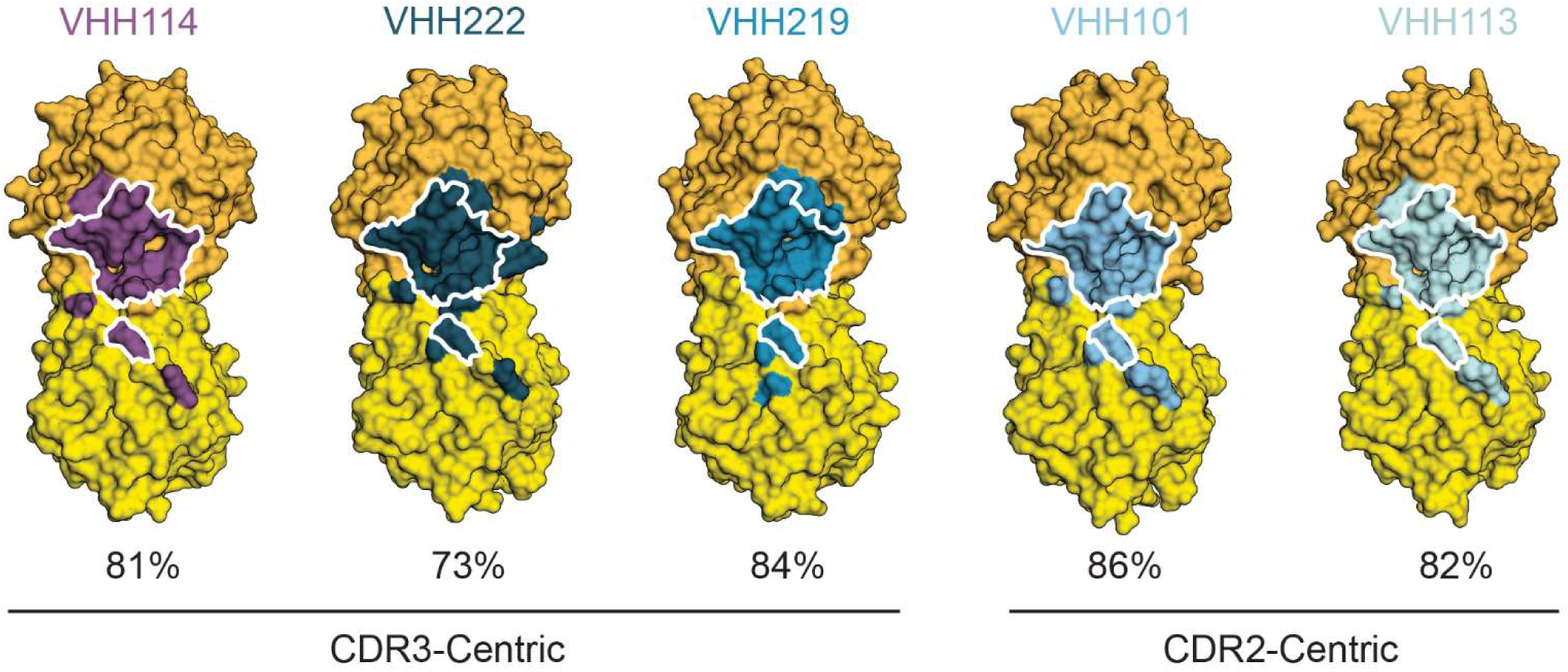
VHHs from both classes preserve a shared binding footprint. All five VHHs footprint to the same regions of Cif and recognize shared core epitopes demarcated by a white outline. The percentage contribution of residues located in the core epitope to the total epitope for each VHH is displayed below the epitope maps. A two-tone color scheme for Cif was chosen to differentiate epitopes on the proximal Cif monomer (orange) and the distal Cif monomer (yellow). Epitopes are colored according to the nanobody (VHH114, purple; VHH222, dark blue; VHH219, blue; VHH101, light teal; VHH113, faded denim).

To better understand the molecular recognition strategies exploited by these two structurally distinct paratopes, a pairwise side-by-side comparison of the individual stereochemical interactions in the epitope:paratope interface of the VHH222 (CDR3) and VHH113 (CDR2) nanobodies was performed. A single relatively conserved interaction is found. Positioned near the axis of rotation, the amide nitrogen of Cif His269 can hydrogen bond with the carbonyl oxygen atom at position 35 of CDR1 from either VHH113 or VHH222 (Fig 13A, left-hand panel), although the angle of interaction differs.

**Figure 13.**
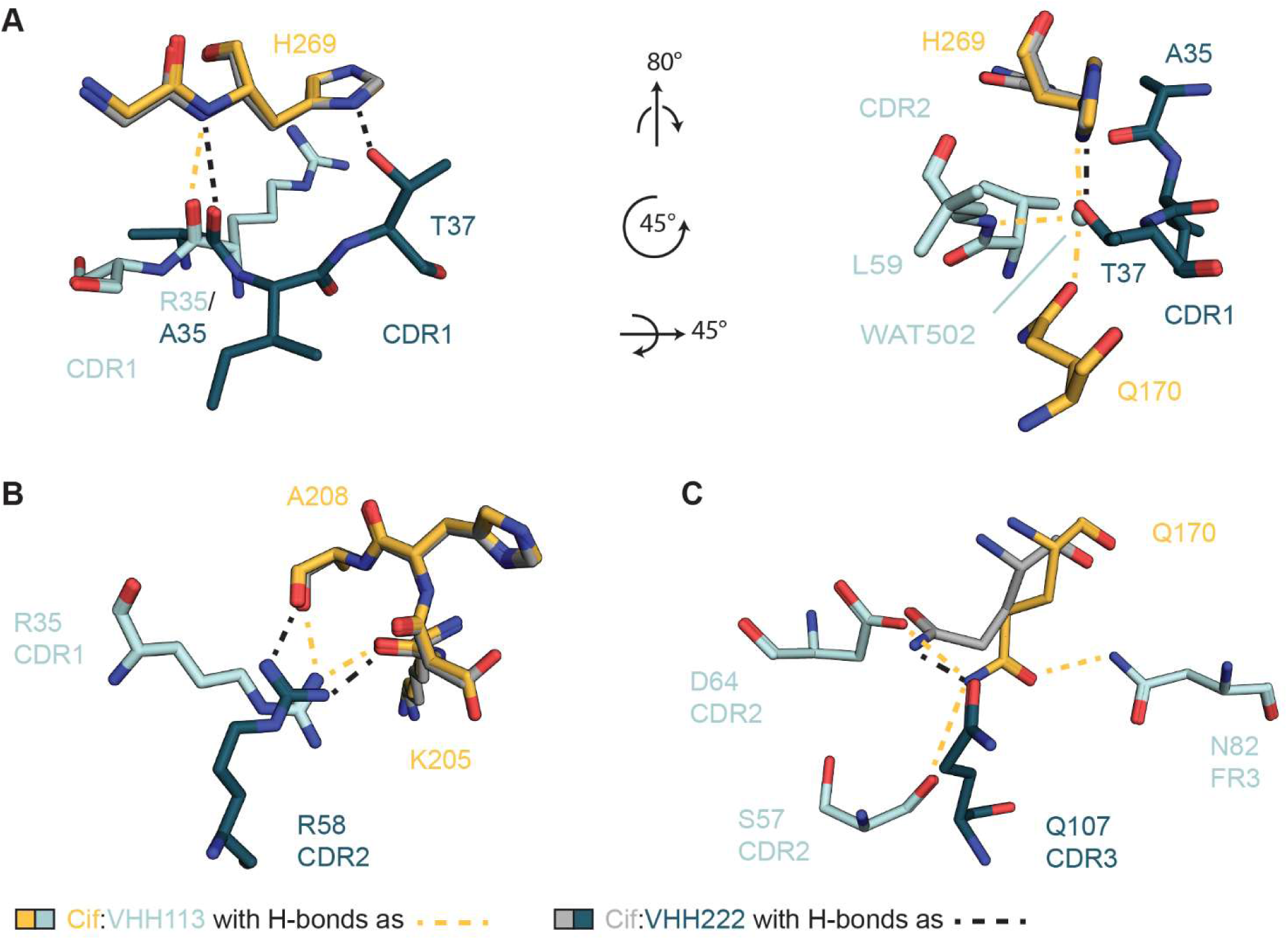
VHH113 and VHH222 make different interactions with the same Cif atoms underpinning a shared footprint. Despite having structurally unique paratopes of differing amino-acid composition, VHH113 and VHH222 interact with a common set of Cif residues. (A) A hydrogen bond between VHH222 CDR1 Thr37 and Cif His269 is maintained by structural water WAT502 and Leu59 of CDR2 at the Cif:VHH113 interface. (B) Arginine side chains extending from CDR1 (VHH113) or CDR2 (VHH222) hydrogen bond with the same carbonyl oxygens of Cif. (C) VHH222 uses a side chain on CDR3 to form a hydrogen bond with Cif Gln170. VHH113 forms three hydrogen bonds with the same Cif residue using two residues from CDR2 and one from FR3. Cif:VHH113 complex, orange/faded denim; Cif:VHH222 complex, grey/dark blue; hydrogen bonds between Cif and VHH113, orange dashed line; hydrogen bonds between Cif and VHH222, black dashed line. Non-carbon atoms are colored by types: oxygen, red; nitrogen, blue.

With the exception of this example, in general, the nanobody side chains forming each paratope are quite different between the two classes (Figs 5 and 8), as may be expected for independent antigen combining sites even with a common target. In contrast, essentially identical sets of Cif residues within the core epitope provide the specific stereochemical anchors for nanobody attachment in both classes. Again using VHH222 to represent CDR3 inhibitors and VHH113 to represent CDR2 inhibitors, the ways in which different paratopes engage a shared epitope become clear.

A prime example involves the reverse turn that in each paratope engages the active-site gate, as seen in a comparison of Fig 8A (leftmost panel) and Fig 5B (rightmost panel) or of Fig 8A [remaining panels] and Fig 5D. While the targets on Cif are identical, the reverse turn is formed in VHH113 and VHH222 by distinct CDRs (CDR2 and CDR3, respectively), as described in detail above. Surrounding the active-site gate, other examples are found. There is a hydrogen bond between the indole ring of Cif His269 and VHH222 Thr37. In complex with VHH113, a water molecule (WAT502) occupies the location corresponding to the VHH222 threonine hydroxyl group, where it mediates an indirect hydrogen bond between Cif His269 and VHH113 Leu59 (Fig 13A, right-hand panel). As another example, the carbonyl oxygens of Cif Lys205 and Ala208 form hydrogen bonds in each interface, but do so by engaging the side chain of either Arg35 (VHH113) or Arg58 (VHH222), which approach from different directions (Fig 13B). Cif Gln170 also participates in each interface, forming either a single hydrogen bond with Gln107 when bound to VHH222, or else a network that includes VHH113 residues Asp64, Ser57, and Asn82 (Fig 13C), as well as the water molecule (WAT502) that engages Cif His269, as noted above (Fig 13A, right-hand panel). In total, all of the Cif side chains involved in electrostatic, hydrogen-bond, or van der Waals interactions at the interface of Cif:VHH222 are also engaged in the Cif:VHH113 complex, even though they often engage non-corresponding VHH side chains located in different CDR loops when binding each of the two classes of nanobodies.

### Summary and concluding remarks

The five Cif:nanobody co-crystal structures presented here provide mechanistic insights regarding nanobody-mediated inhibition of a PA virulence factor implicated in the worsening of CF lung pathophysiology. Mapping of the Cif:VHH interactions to the CDR sequences revealed a class of nanobodies that share binding characteristics common to VHH219 and VHH222. A defining feature of this class is an extended CDR3 conformation (Fig 9B) that reaches toward the gating mechanism (Fig 1), where a Type I reverse turn-like structure plugs the active-site entrance with a hydrophobic side chain (Trp/Tyr/Val) at the *i*+1 position. These findings were structurally validated using VHH114 as a case study and further assessed biochemically through competition hydrolysis assays that identified two previously unrecognized sequences (VHH314 and VHH320) as Cif inhibitors. Based on steric considerations, we propose these nanobodies act as strictly competitive inhibitors, with the exception of VHH114 which may conditionally act as a non-competitive inhibitor for a handful of small substrates.

These inhibitory nanobodies interact with surfaces that are inaccessible in the apo Cif structure. Although such cryptic epitopes are not unique,^99–101^ they have implications for the kinetics of the Cif interaction. Unless held in the open conformation by some type of interacting molecule (substrates or inhibitors), the Cif active-site gate has consistently been found in a closed conformation.^23,49,90^ So far, this is the only known entry point to the active site, suggesting that access to the active site is controlled in part by the conformational dynamics of Phe164 and Leu174 of the cap domain and Met272 of the core domain. These so-called gate-keeper residues are thought to play a pivotal role in regulating the movement of molecules in and out of the active site in the broader class of α/β-hydrolases with immobile cap domains.^102^ When these side chains splay apart (Fig 2), a tunnel is revealed, connecting the exterior to the active site.^49,91^ As the hydrophobic side chains of the nanobodies make surface contacts with portions of this tunnel, these gate-keeper residues serve a secondary purpose in concealing a portion of a functionally important antigenic surface. In effect, the entirety of the epitope may be only transiently accessible to these inhibitory nanobodies, when Cif adopts an open conformation.

A second class of inhibitory nanobodies uses CDR2 in place of CDR3, underscoring the versatility of immunoglobulin scaffolds and variable CDR loops in addressing molecular recognition challenges. Although often shorter and less conformationally diverse than CDR3,^29,32^ CDR2 can nonetheless drive enzyme inhibition.^34^ In our particular case, the non-canonical CDR2 of both nanobodies forms the centerpiece of the paratope while CDR3 makes minimal contact with Cif in an apparent role inversion.

In this class of nanobodies, the inhibitory CDR2 reported here encodes a pentad motif shared with the CDR3-centric nanobodies, leading to a nearly identical mechanism of Cif inhibition, as opposed to an independent “orthogonal” strategy, as has been reported in some cases.^103,104^ In other studies, shared sequence motifs in antibody CDRs appear to have been driven by convergence towards substrate mimicry.^105,106^ That does not appear to be the case here. Although similar to each other, the CDR pentads lack stereochemical similarity to known Cif substrates or inhibitors, and thus represent a departure from traditional forms of ligand _mimicry.35,107,108_

Consistent with findings in other cases,^109,110^ the extended vs. kinked CDR3 conformations observed in the Cif:VHH structures (Figs 9A and 9B) are associated with different V-gene alleles. Motif convergence was thus presumably mediated by ensuing somatic hypermutation and selection during affinity maturation. The resulting nanobodies of both classes favor a reverse-turn L(S/A)(W/Y)XG pentad motif with a hydrophobic (often aromatic) residue at the corner position *i*+1, which enhances the ability of its side chain to access the cryptic active site of Cif. Tryptophan and tyrosine residues also tend to be highly abundant in paratopes^31,32,111–113^ and in some instances, exhibit preferential placement into specific pockets.^99,104^ In the case of Cif, this preference may complement an active site designed to hydrolyze hydrophobic substrates. By targeting the active-site tunnel, the Cif-specific nanobodies benefit from the physical proximity of stereochemical “handles” within a typical enzyme catalytic site. Indeed, this feature may be a strategy that specifically benefits nanobodies that inhibit enzymes, as Akiba *et al.* have reported a similarly high density of binding energies between a nanobody and the active site of lysozyme.^84^

Overall, both classes of inhibitory nanobodies footprint to the same region of Cif. The overlap of epitopes is not surprisingly *per se*: clusters of overlapping epitopes have been identified in other antigen:nanobody and antigen:antibody complexes,^41,107,114–119^ likely reflecting the propensity of select antigenic surfaces to provoke robust affinity maturation. Within the shared epitope, the two classes of nanobodies exhibit a nearly 90° relative macroscopic domain reorientation around a central pivot near CDR1, which is required to position the convergent pentads of either the CDR3 or CDR2 loops, respectively, above the active-site gate. However, this relative paratope reorientation also fundamentally disrupts the majority of the individual paratope:epitope interactions that contribute to either class interface. In addition, the paratopes of these classes are stereochemically distinct. Nonetheless, both classes of nanobodies manage to engage nearly identical sets of handholds on the Cif epitope. A side-by-side comparison of the pairwise interactions reveals how each class of inhibitory nanobody forms distinct compensatory interactions with this common set of stereochemical anchor points (compare Figs 5 to 8 and see Fig 13).

In effect, each class of nanobodies thus represents an independent paratope solution to the highly constrained structural challenge of recognizing each stereochemical moiety within a large, shared epitope. A similar challenge is posed by efforts to achieve *de novo* design of VHHs to target a user-specified therapeutic epitope, an area that has seen dramatic recent advances.^120^ Nanomolar affinities can be realized by coupling state-of-the-art computational design pipelines with mutational strategies and HTS libraries to emulate affinity maturation.

However, such workflows bring forth the added challenge of defining an ideal epitope. In the absence of a tacit solution supplied by the immune system, inadvertent selection of sub-optimal epitopes may impede *in silico* design efforts from the beginning, leaving downstream computational designs trapped in low-affinity constructions. The availability of experimental examples of multiple, independently derived nanobody paratopes engaging with shared epitopes leverages the molecular creativity of the immune system to expand our understanding of strategies to create high-affinity binding interactions targeting a defined epitope, and may lead to the improvement of *in silico* design methods.

The inhibitory Cif nanobodies demonstrate this concept. Utilization of nearly identical sets of epitope atoms on Cif is unlikely to be a coincidence, but rather a reflection of a naturally occurring optimal grip that has been developed through affinity maturation of separate nanobody lineages. To this point, nanobodies belonging to each class both exhibit low-nanomolar (or better) affinities. In addition, the Cif nanobodies underscore the importance of recognizing the connection between the binding pose of the nanobodies and the sequence-structure relationship of the CDR loops. For VHH113 and VHH101, the interaction of the reverse turn with the active-site gate is made possible not only by the relative pivot of the domain, but also by a kink in the non-canonical CDR2. Indeed, there may be value in the systematic pursuit of reference enzyme:nanobody structures for sets of inhibitory nanobodies with sequences that have distinct allelic origins. While allosteric inhibition mechanisms exist, the selection for potent enzyme inhibitors will enhance the likelihood that many of the epitopes will be centered around active-site entrances. At the same time, affinity maturation should enable different allelic starting points to converge on distinct solutions to the binding challenge at hand. While such studies are time consuming, a relatively small number of datasets could provide invaluable training sets for more refined computational models of paratope:epitope recognition, and more generally, for the sequence-structure relationships that underlie their formation. In parallel, they would test the generality of the observations made here with Cif and would generate versatile reagents for the study of the target enzymes in their physiological systems.

## Acknowledgements

Research reported in this publication was supported in part by NIH awards R01-AI091699, T32-AI007519 (A.R.S.), R35-ES030443 (NIEHS RIVER Award; C.M.), and U19-AI45825 (A.P.H.). Additional core support was provided by the Institute for Biomolecular Targeting (P20-GM113132), the Dartmouth CF Research Center (P30-DK117469 and Cystic Fibrosis Foundation award BOMBER24R0), the Center for Quantitative Biology (P20-GM130454), the Dartmouth Cancer Center Genomics and Molecular Biology Shared Resources (RRID: SCR_021293; P30-CA023108) and the Center for BioMolecular Structure (P30-GM133893 and DOE Office of Biological and Environmental Research award KP1605010). This research used the AMX (17-ID-1) and FMX (17-ID-2) beamlines of the National Synchrotron Light Source II, a U.S. Department of Energy (DOE) Office of Science User Facility operated for the DOE Office of Science by Brookhaven National Laboratory under Contract No. DE-SC0012704. We would also like to thank Dr. Kai Diederichs for consultation for data reduction of rotation X-ray diffraction data collected from the macroscopically twinned Cif:VHH113 crystal (pdb_00008f6u), Dr. Stacie Balsam for assisting with crystallization trials of VHH222 in its free form, and Dr. Nicholas Gill for consultation during refinement of CMNGC hydrolysis assays and helpful suggestions for data visualization. This study is dedicated to the memory of Dr. Bruce D. Hammock: husband, father, scientist, mentor, collaborator, friend (1947-2026).

## Author Contributions

A.R.S., N.M.T., and D.R.M. conceptualized the study and designed the experiments; A.R.S., N.M.T., A.K.M., K.S.B., and A.P.H. conducted experiments; A.R.S., A.P.H., T.H.H., and D.R.M. analyzed the data; T.H.H. and A.R.S. wrote R scripts for data wrangling and visualization; N.V., C.M., M.E.A., and D.R.M. provided laboratory resources; D.R.M. and M.E.A. were responsible for funding acquisition; A.R.S. and D.R.M. wrote the initial drafts of the manuscript. All authors edited and approved the final manuscript.

## Footnotes

† The VHH113/VHH101 CDR2 loop is 10 residues long following IMGT nomenclature but 13 residues long following the convention used by North *et al*.^97^ in the updated canonical CDR loop conformations.

## Notes

https://www.rcsb.org/structure/8E2N

https://www.rcsb.org/structure/8E1C

https://www.rcsb.org/structure/8EVD

https://www.rcsb.org/structure/8F6U

https://www.rcsb.org/structure/8GJR

https://www.rcsb.org/structure/8EE2

https://www.rcsb.org/structure/8ELN

### Competing Interest Statement

The authors have declared no competing interest.

### Summary of Updates

A correction was made in the acknowledgements section: spelling and attributing help during the data reduction of X-ray dataset for pdb_00008evd (incorrectly listed as pdb_00008f6u).

