## Supplementary Material for "Convergent strategies for nanobody-mediated inhibition of an epoxide hydrolase"

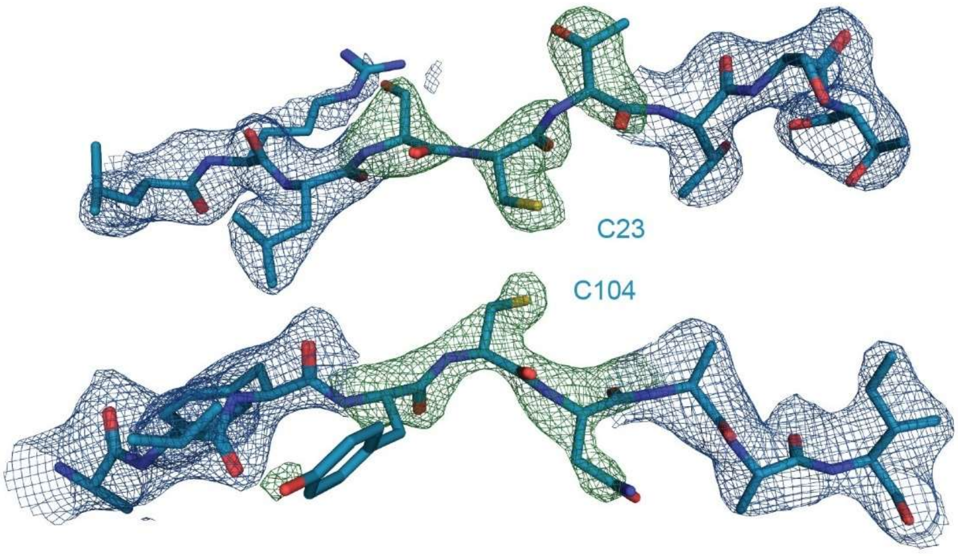

**Supplementary Figure 1. Canonical disulfide bond did not form in VHH219.** 2m$F_O$-D$F_C$ (blue mesh) and m$F_O$-D$F_C$ (green mesh) electron density maps were calculated after molecular replacement with cysteines and flanking residues omitted. The resulting omit map is contoured at 3 $\sigma$ and shows no indication of a disulfide bond. Fitment of the refined VHH219 structure with cysteines modeled in the reduced form are in good agreement with the omit map. VHH219, blue. Non-carbon atoms are colored by type: oxygen, red; nitrogen, blue; sulfur, yellow.

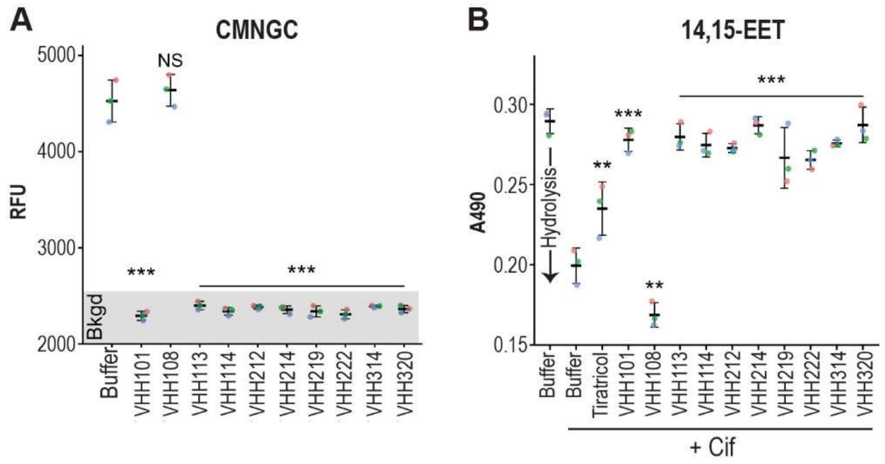

**Supplementary Figure 2. Inhibition of Cif by Nanobodies.** Cif hydrolysis was measured in the presence of saturating concentrations of each nanobody to ascertain inhibitory potential. (A) Nanobodies were tested for inhibitory activity using the fluorogenic reporter substrate CMNGC. Reactions consisted of 1.2 µM Cif, 5.5 µM VHH (or buffer control), 25 µM CMNGC, 15 µM thesit, and 1% (*v/v*) DMSO and were incubated at 37°C for 15 min. (B) Nanobodies were tested for inhibitory activity using 14,15-EET. Reactions consisted of 20 µM Cif, 25 µM VHH (or buffer control), 1 mM 14,15-EET, and 2.5% (*v/v*) DMSO. The Cif inhibitor Tiratricol was used as a positive control for inhibition and was included at 25 µM. All reactions were incubated at 37°C for 60 min. Replicates are shown as colored dots and the means are depicted as horizontal bars with vertical lines equal to one standard deviation. Significance was assessed by ANOVA using Dunnett's post-hoc test, $n$ = 3.

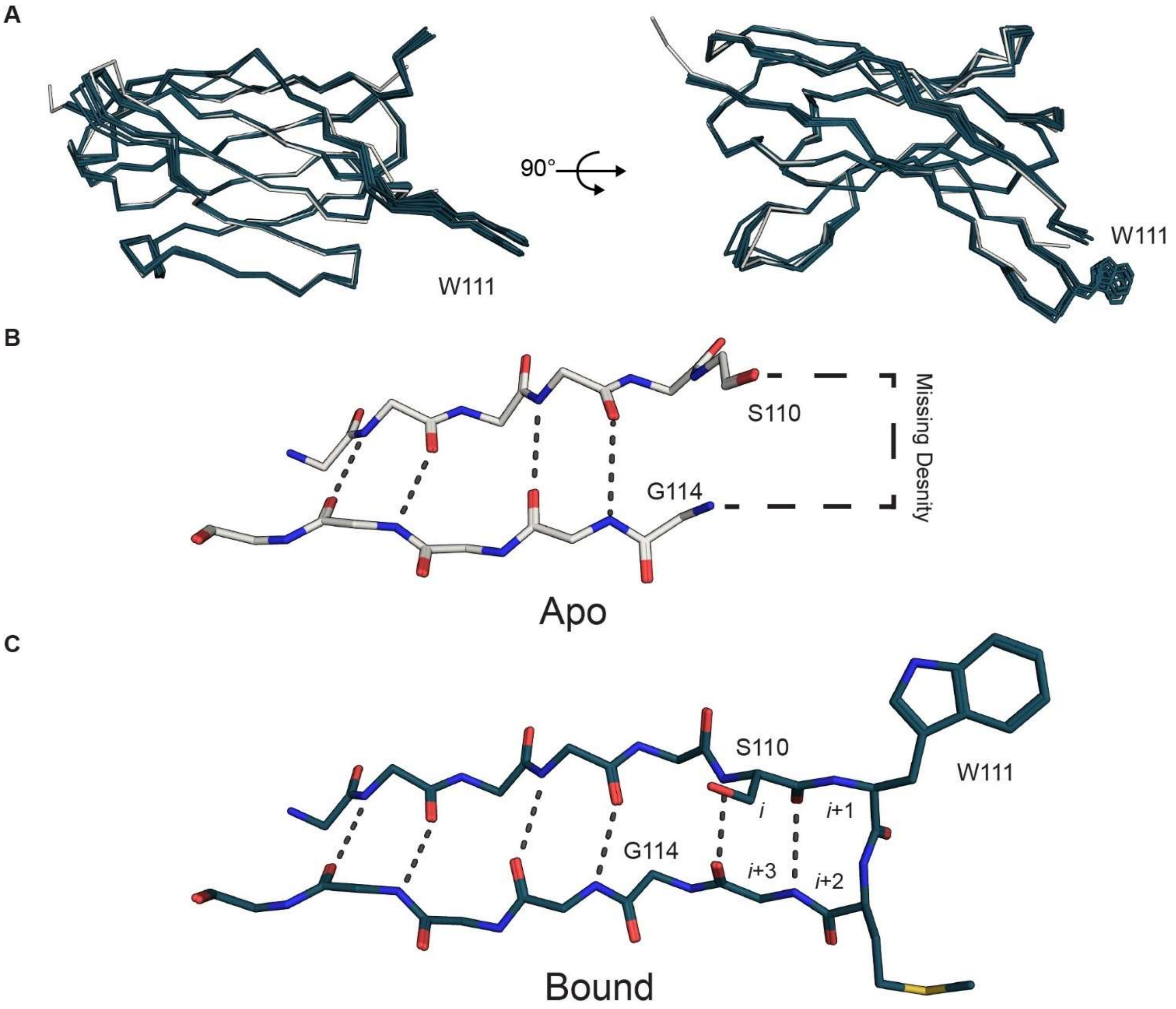

**Supplementary Figure 3. Comparison of VHH222 in free and bound states.** (A) Main-chain alignment of VHH222 in its free form (bone) and all VHH222 chains in the Cif:VHH222 co-crystal structure (dark blue). The side chain of Trp111 is depicted as sticks for orientation. (B) The reverse turn of VHH222 CDR3 was unresolved in the apo structure. (C) CDR3 is stabilized upon binding to Cif, and residues 110:113 form a Type 1 reverse turn-like structure with Trp111 at *i*+1. Non-carbon atoms are colored by type: oxygen, red; nitrogen, blue; sulfur, yellow.

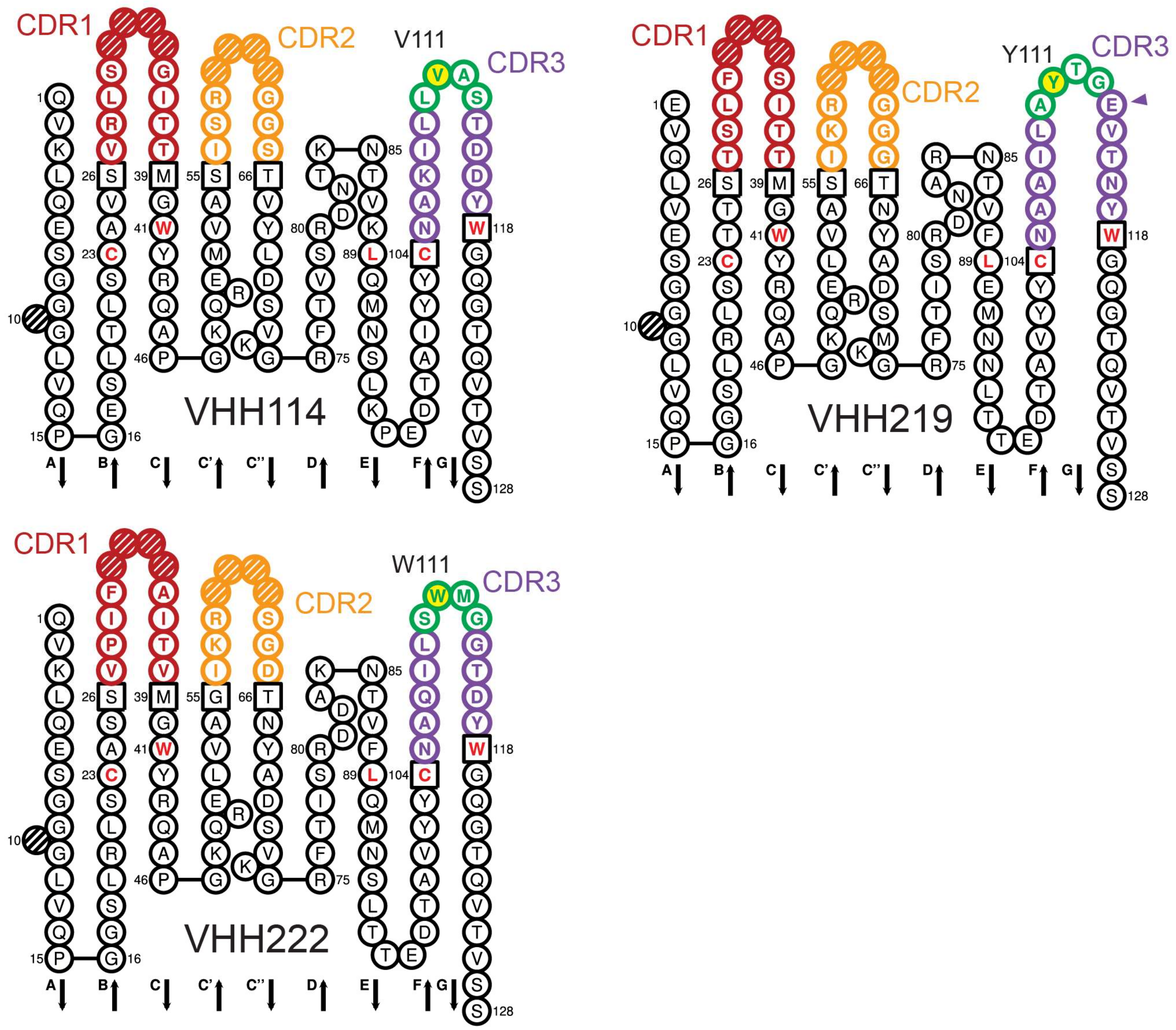

**Supplementary Figure 4.** ***Collier de Perles*** **diagrams highlight sequence variation among CDR3 loops involved in Cif inhibition**. Although each VHH uses CDR3 to inhibit Cif via direct steric occlusion via insertion of a hydrophobic side chain through the active-site gate, a surprising degree of variation is observed in the sequences composing the Type I reverse turn-like structure. Despite these differences, each loop forms highly similar structures underscoring the convergence of a structural motif to carry out inhibition that can tolerate some degree of sequence diversity. Notably, VHH114 has a small hydrophobic side chain while VHH219 and VHH222 have bulky hydrophobic side chains at the equivalent positions. The CDR3 of VHH219 also has an extra amino acid at the top of the loop following the turn (purple triangle). *Collier de Perles* diagrams of VHH114, VHH219, and VHH222 were made using the IMGT/Collier-de-Perles web server [74,121-123] and re-colored to highlight the location of the Type 1 reverse turn (green/white) and the hydrophobic sidechain at position 111 (green/yellow). CDRs are colored as follows: CDR1 (red), CDR2 (orange), CDR3 (purple). Gapped positions are shown with hatch marks. Conserved FR residues are shown in red text and anchor positions appear in black boxes.

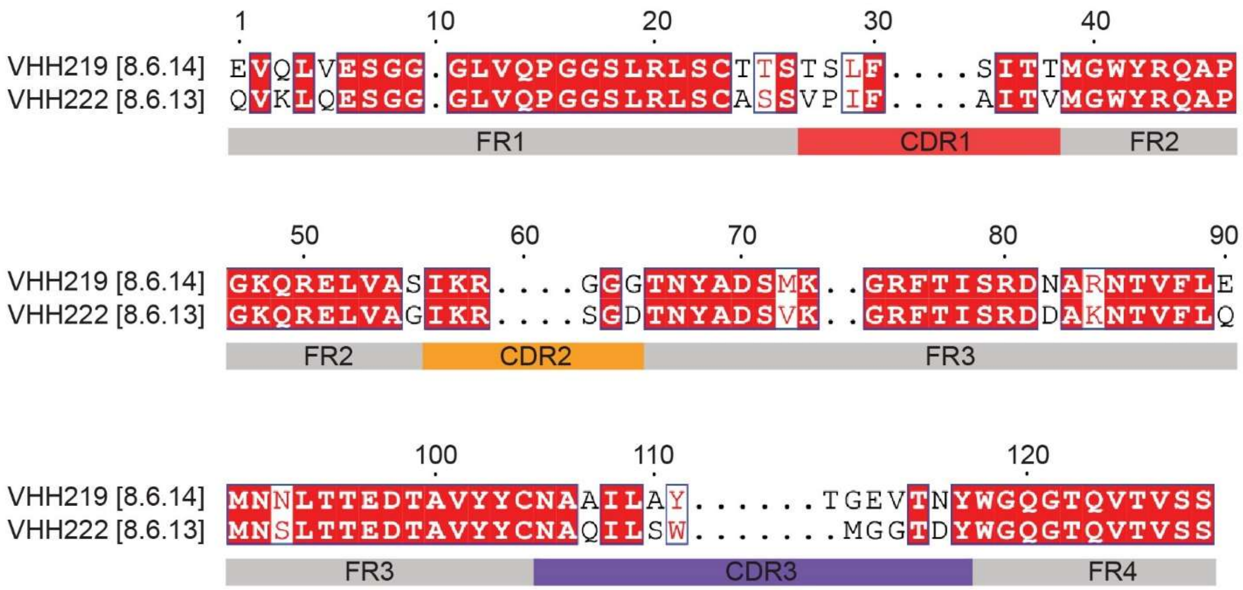

**Supplementary Figure 5. VHH219 and VHH222 pairwise alignment.** Sequence conservation between VHH219 and VHH222 is highest in the framework (FR) regions with most of the variation occurring in the CDRs. In total, only 13 out of 28 CDR positions are conserved. The sequence of CDR2 has more conserved positions (4 out of 6) than CDR1 (3 out of 8) or CDR3 (6 out of 14). Sequence numbering and annotation follows guidelines established by IMGT unique numbering with dots representing gapped positions. CDR lengths are provided in brackets as follows: [CDR1.CDR2.CDR3].

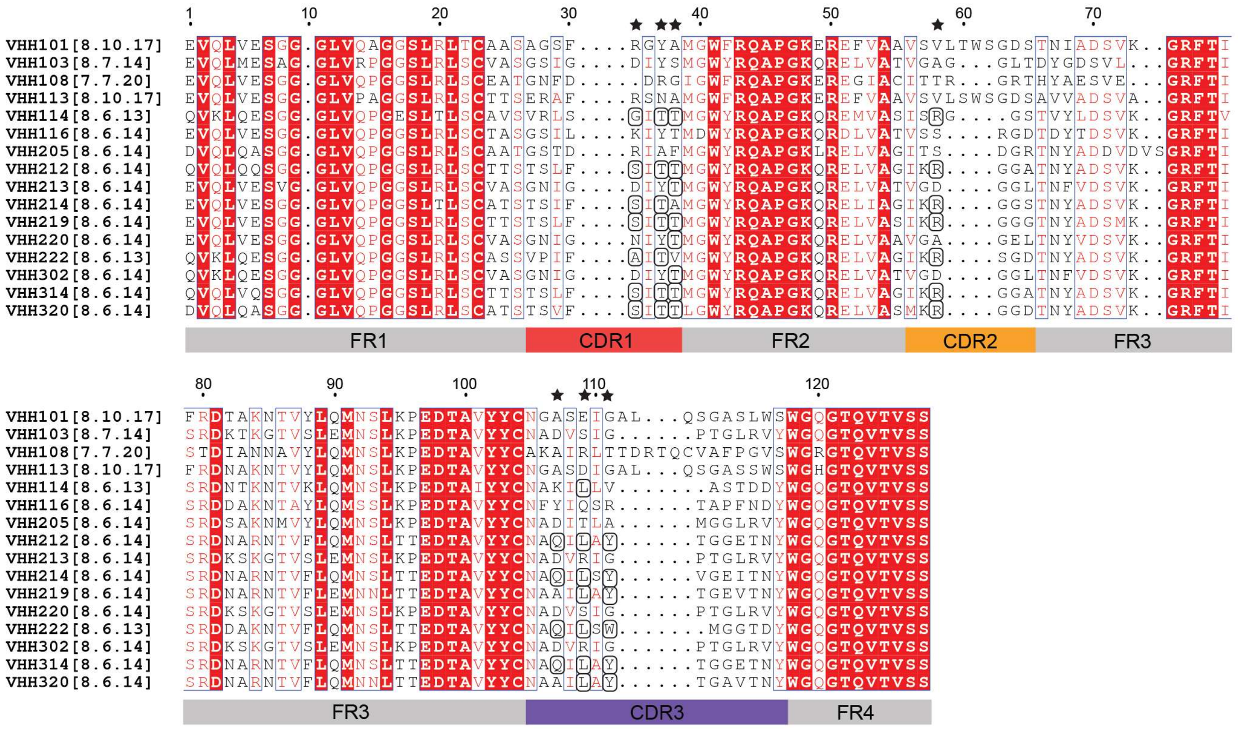

**Supplementary Figure 6. Multiple sequence alignment of nanobody library.** Interactions identified at the Cif:VHH219 and Cif:VHH222 interfaces were used to identify related nanobodies. Sequence numbering and annotation follows guidelines established by IMGT unique numbering with dots representing gapped positions. CDR lengths are provided in brackets as follows: [CDR1.CDR2.CDR3]. Black stars mark CDR positions involved in interactions found at the Cif:VHH219 and Cif:VHH222 interface. Residues predicted to maintain the interaction are circled. See Figure 6 for specifics regarding interactions.

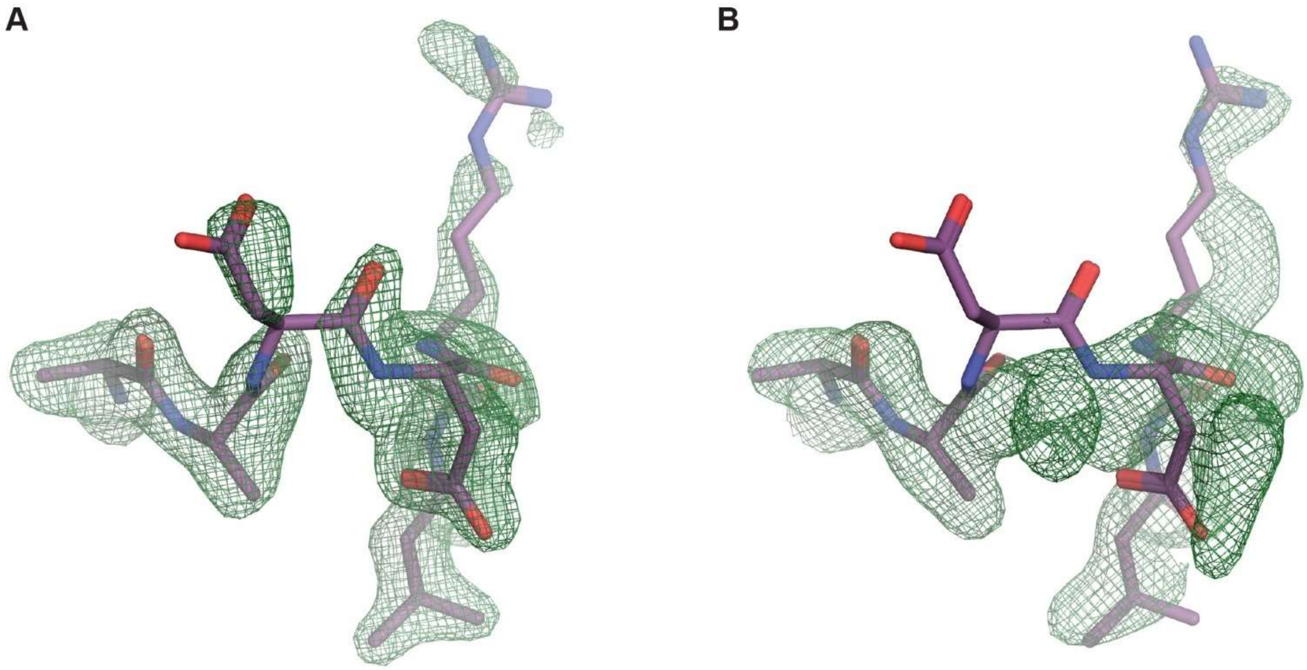

**Supplementary Figure 7. Main-chain rearrangements in the crystal structure of Cif:VHH114 are artefacts of crystallization.** The published structure of Cif:VHH114 depicts a shift in the main chain due to the presence of two nearby citrate ions from the crystallization buffer. After phasing, omit maps were calculated for the deposited coordinates (A) and for a second dataset in an alternate crystal system that grew without citrate in the crystallization buffer (B) (SBGrid Data Bank: dataset 1057). The resulting m$F_O$-D$F_C$ maps are contoured at 3 $\sigma$ and the coordinates of the deposited structure (pdb_00008gjr) are displayed in each map. Notably, the omit maps are different in this region. Non-carbon atoms are colored by type: oxygen, red; nitrogen, blue.

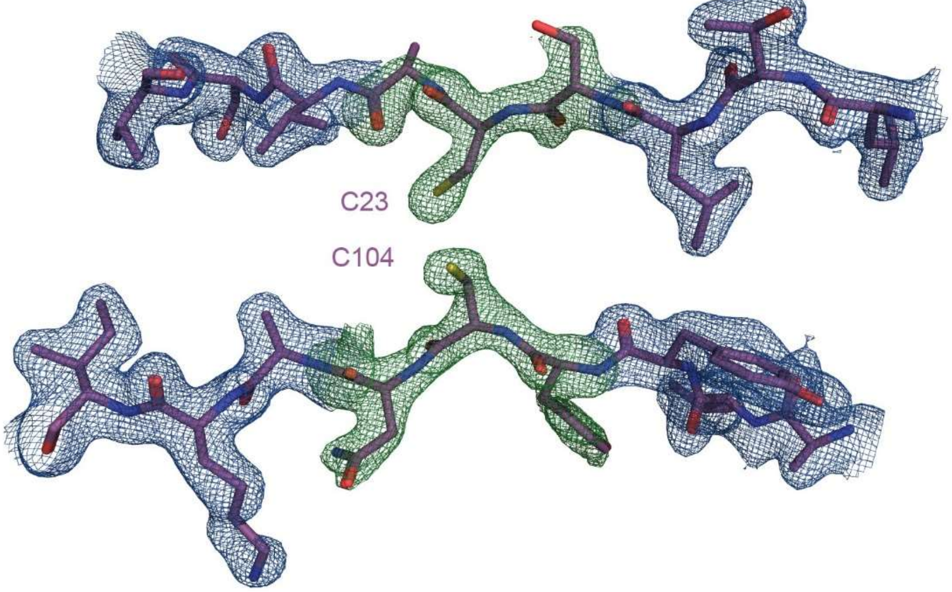

**Supplementary Figure 8. Canonical disulfide bond did not form in VHH114.** 2m$F_O$-D$F_C$ (blue mesh) and m$F_O$-D$F_C$ (green mesh) electron density maps were calculated after molecular replacement with cysteines and flanking residues omitted. The resulting omit map is contoured at 3 $\sigma$ and shows no indication of a disulfide bond. Fitment of the refined VHH114 structure with cysteines modeled in the reduced form are in good agreement with the omit map. VHH114, purple. Non-carbon atoms are colored by type: oxygen, red; nitrogen, blue; sulfur, yellow.

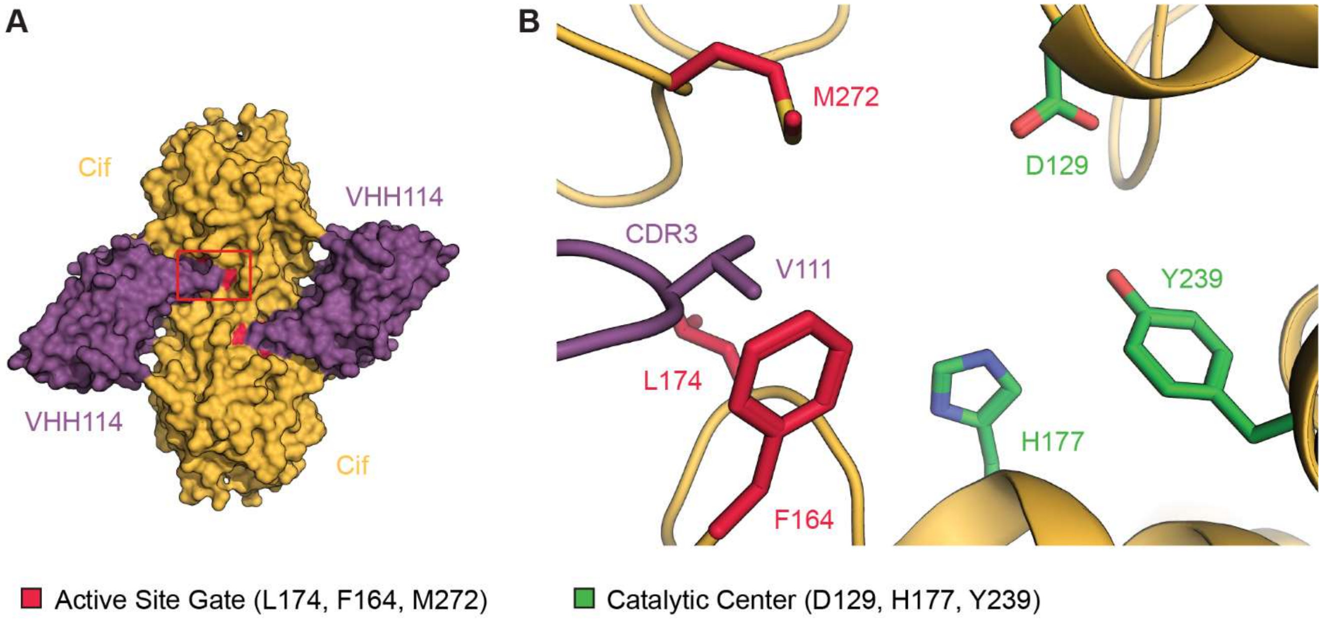

**Supplementary Figure 9. Co-crystal structure of Cif:VHH114 reveals a valine functionally replaces the aromatic "stopper."** The Cif:VHH114 complex bears strong resemblance to that of Cif:VHH219 and Cif:VHH222 in terms of 2:2 stoichiometry and equatorial binding of VHH114 near the Cif dimer interface (compare to Figure 1). (A) The Cif:VHH114 complex is illustrated as surface representations with Cif in orange and VHH114 in purple. The red box highlights a portion of CDR3 covering the active-site entrance (red) of the upper protomer of Cif. (B) The entryway is occupied by Val111 of CDR3. Although Val111 occupies a smaller steric volume than Tyr111 and Trp111 of VHH219 and VHH222, respectively, the valine side chain is sufficient to block the active-site entrance and cause similar perturbations in the gate-keeper residues (red sticks). Select residues in the catalytic center (nucleophile Asp129, ring-opening pair His177 and Tyr239) are shown as green sticks for orientation. Non-carbon atoms are colored by type: oxygen, red; nitrogen, blue; sulfur, yellow.

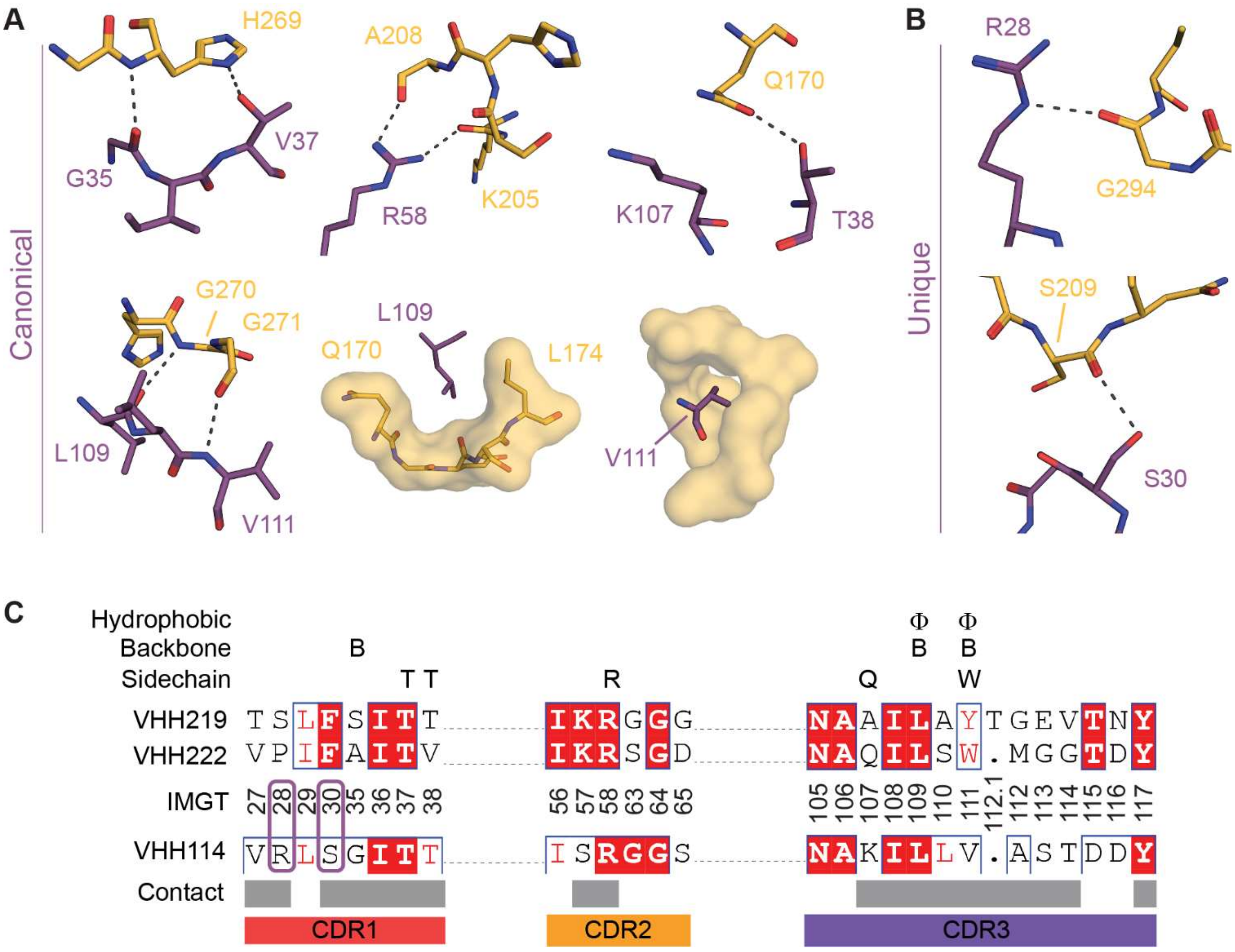

**Supplementary Figure 10. Contact analysis of Cif:VHH114 interface.** Interactions found at the Cif:VHH114 interface were identified, categorized, and compared to those identified at the Cif:VHH219 and Cif:VHH222 interfaces. (A) Cif:VHH114 interactions also found at the interface of Cif:VHH219 and/or Cif:VHH222 and referred to as canonical interactions (compare to Figure 5). (B) VHH114 makes two unique interactions with Cif that are not present in the Cif:VHH219 or Cif:VHH222 complexes. (C) Pairwise alignment of CDRs from VHH219, VHH222, and VHH114. TOP – Reference interactions identified at the Cif:VHH219 and Cif:VHH222 interface. BOTTOM – Comparison of interactions identified at the Cif:VHH114 interface. Purple ovals highlight VHH114 amino acids involved in distinct interactions displayed in (B). Sequence numbering and annotation follows guidelines established by IMGT unique numbering with dots representing gapped positions. Categories are as follows: *Hydrophobic* indicates the burial of hydrophobic VHH side chains into specific pockets located on Cif; *Backbone* indicates hydrogen bonds involving main-chain atoms of residues in a VHH CDR; *Sidechain* indicates hydrogen bonds involving side-chain atoms of residues in a VHH CDR.  Gray bars indicate residues identified through surface contact analysis. CDRs are annotated as colored bars as follows: CDR1 – red, CDR2 – orange, and CDR3 – purple. Cif, orange; VHH114, purple. Non-carbon atoms are colored by type: oxygen, red; nitrogen, blue.

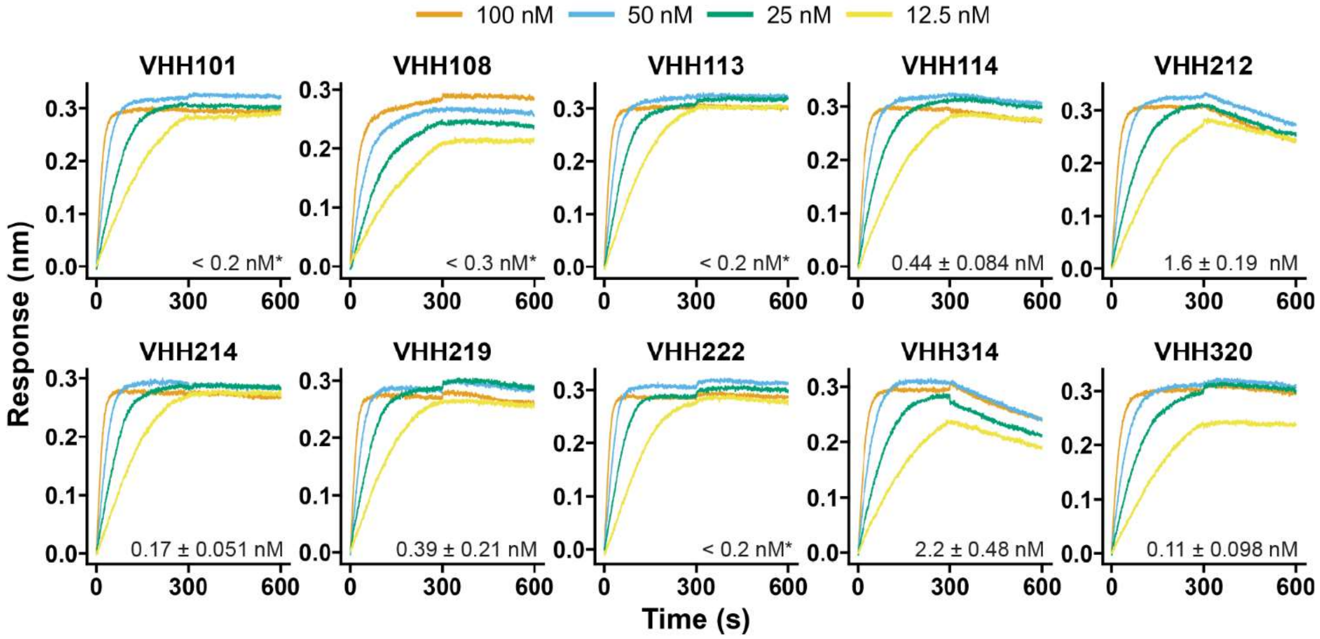

**Supplementary Figure 11. Biolayer interferometry sensograms for binding of nanobodies to Cif.** The affinity for each nanobody was measured by biolayer interferometry to track kinetics of association (300 s) and dissociation (300 s) using a Forte Bio Octet system with immobilized Cif-6xHis. Modeling of the association and dissociation kinetics yielded good fits ($R^2$ > 0.95) with the exception of VHH214, VHH219, and VHH222 tested at 400 nM, which had $R^2$ values of 0.86, 0.88, and 0.87, respectively. For nanobodies marked with an asterisk (*), the error exceeded the estimate of the dissociation phase; for these curves an upper bound was estimated for $K_D$, using the slowest $k_{off}$ value that could be confidently fit ($10^{-4}$ $s^{-1}$). See Methods for details.

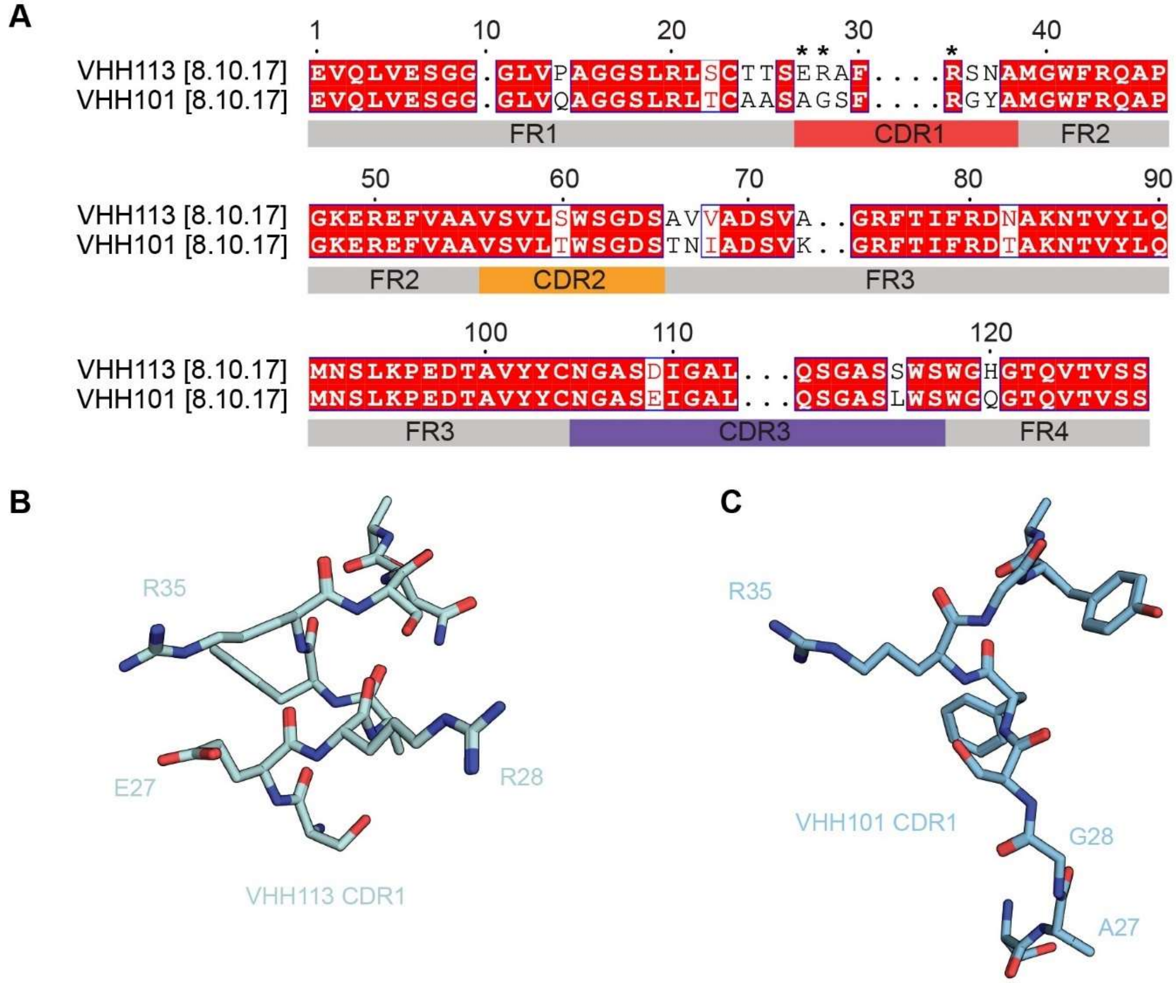

**Supplementary Figure 12. Variation between CDR1 loops of VHH113 and VHH101.**

VHH113 and VHH101 are near identical except for CDR1. (A) A pairwise alignment highlights CDR1 as the only region with substantial differences. In total, 27 out of 35 CDR positions are conserved. CDR1 only conserves 3 out of 8 positions. In contrast, CDR2 conserves 9 out of 10 positions and CDR3 conserves 15 out of 17 positions. Key CDR1 positions are marked with an asterisk (*). CDR lengths following IMGT nomenclature are provided in brackets as follows: [CDR1.CDR2.CDR3]. VHH113 CDR1 (B) and VHH101 CDR1 (C) are shown with key CDR1 positions labeled. The only CDR1-Cif interaction found in both VHHs is mediated by Arg35 due to overlap of the side-chain guanidinium group despite each CDR1 adopting different tertiary structures. Notably, VHH113 Glu27 and Arg28 project toward Cif and participate in H-bonds that are unique to the Cif:VHH113 complex. VHH113, faded denim; VHH101, light teal. Non-carbon atoms are colored by type: oxygen, red; nitrogen, blue.

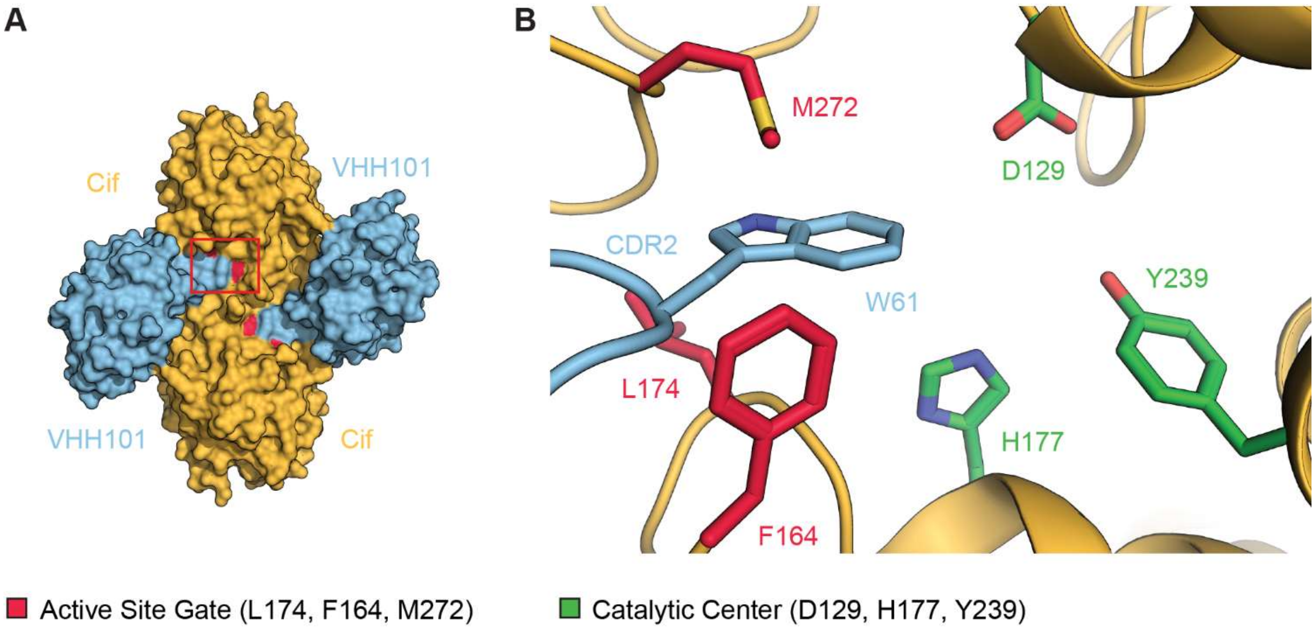

**Supplementary Figure 13. VHH101 is closely related to VHH113 and implements identical mechanisms of Cif inhibition.** VHH101 is a sister nanobody to VHH113 (77% CDR sequence identity, see Figure S12A) and forms a nearly identical complex as the Cif:VHH113 complex (Figure 7) (A) The Cif:VHH101 complex is illustrated as a surface representation with Cif in orange and VHH101 in light teal. (B) VHH101 sterically occludes the active-site entrance using Trp61 of CDR2 in the same manner as VHH113. Active-site gate, red sticks; catalytic center, green sticks. Non-carbon atoms are colored by type: oxygen, red; nitrogen, blue; sulfur, yellow.

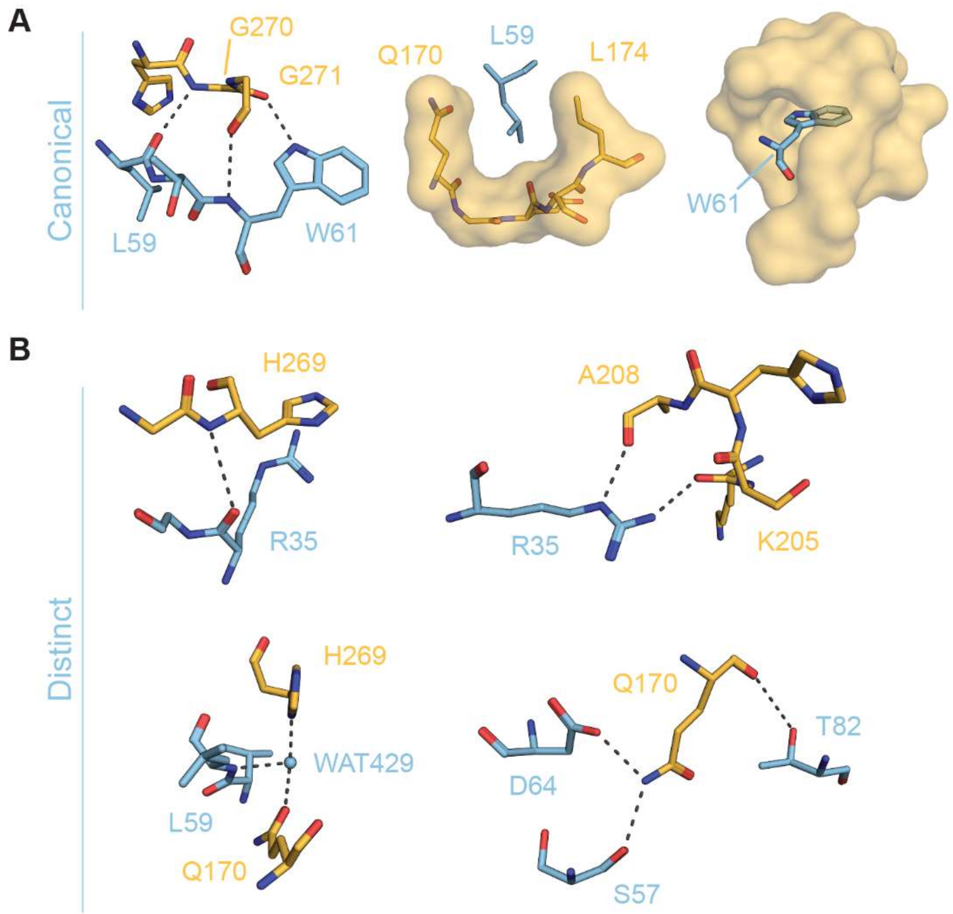

**Supplementary Figure 14. VHH101 makes near identical contacts as VHH113.** Like VHH113, VHH101 exhibits a mixture of canonical and unique interactions that mediate Cif recognition (compare to Figure 8). (A) Interactions in VHH101 CDR2 that perfectly recapitulate those created by VHH113 CDR2 and mimic the interactions of VHH222 CDR3. (B) Cif:VHH101 interactions that are not found at the Cif:VHH222 interface. These are near identical to those found in the Cif:VHH113 complex excluding the VHH113 unique interactions (Figure 8C). Cif, orange; VHH101, light teal. Non-carbon atoms are colored by type: oxygen, red; nitrogen, blue.

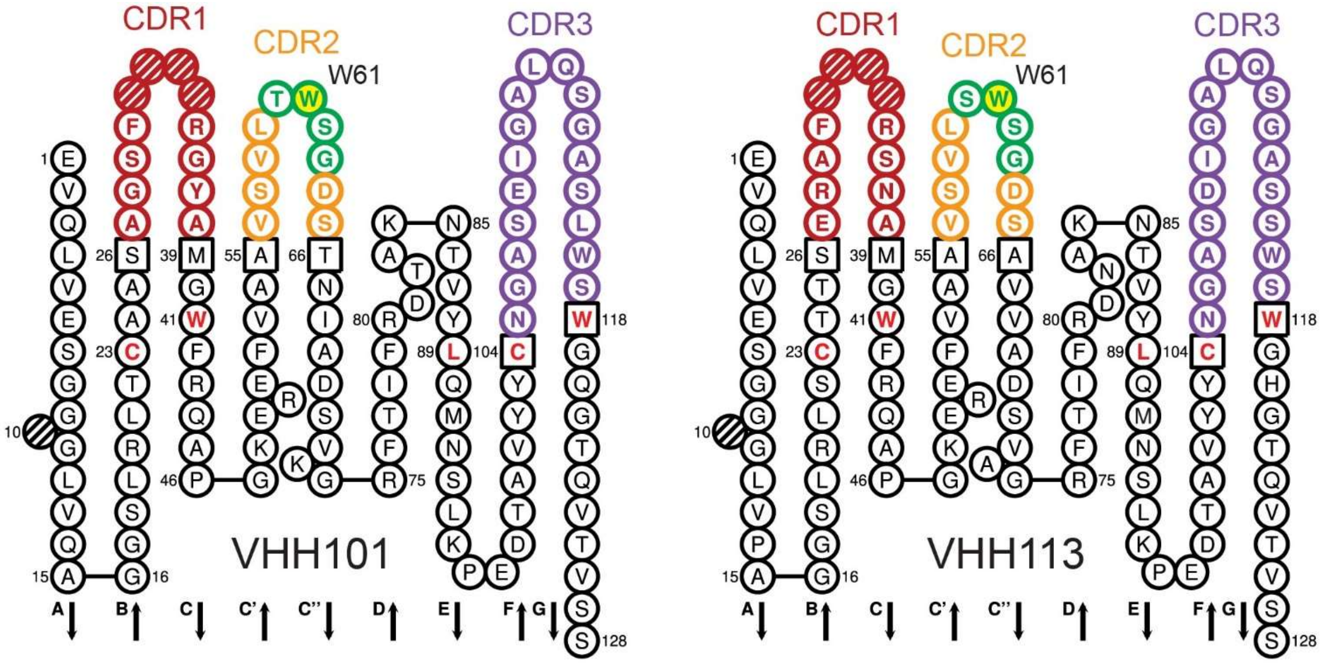

**Supplementary Figure 15**. ***Collier de Perles* diagrams highlight sequence similarities among CDR2 loops involved in Cif inhibition**. While CDR2 and CDR3 are similar, CDR1 demonstrates greater sequence variation that accounts for two additional interactions only found in the Cif:VHH113 complex. *Collier de Perles* diagrams of VHH101 and VHH113 were made using the IMGT/Collier-de-Perles web server [74,121-123] and re-colored to highlight the location of the Type 1 reverse turn (green/white) and Trp61 (green/yellow). CDRs are colored as follows: CDR1 (red), CDR2 (orange), CDR3 (purple). Gapped positions are shown with hatch marks. Conserved FR residues are shown in red text and anchor positions appear in black boxes.

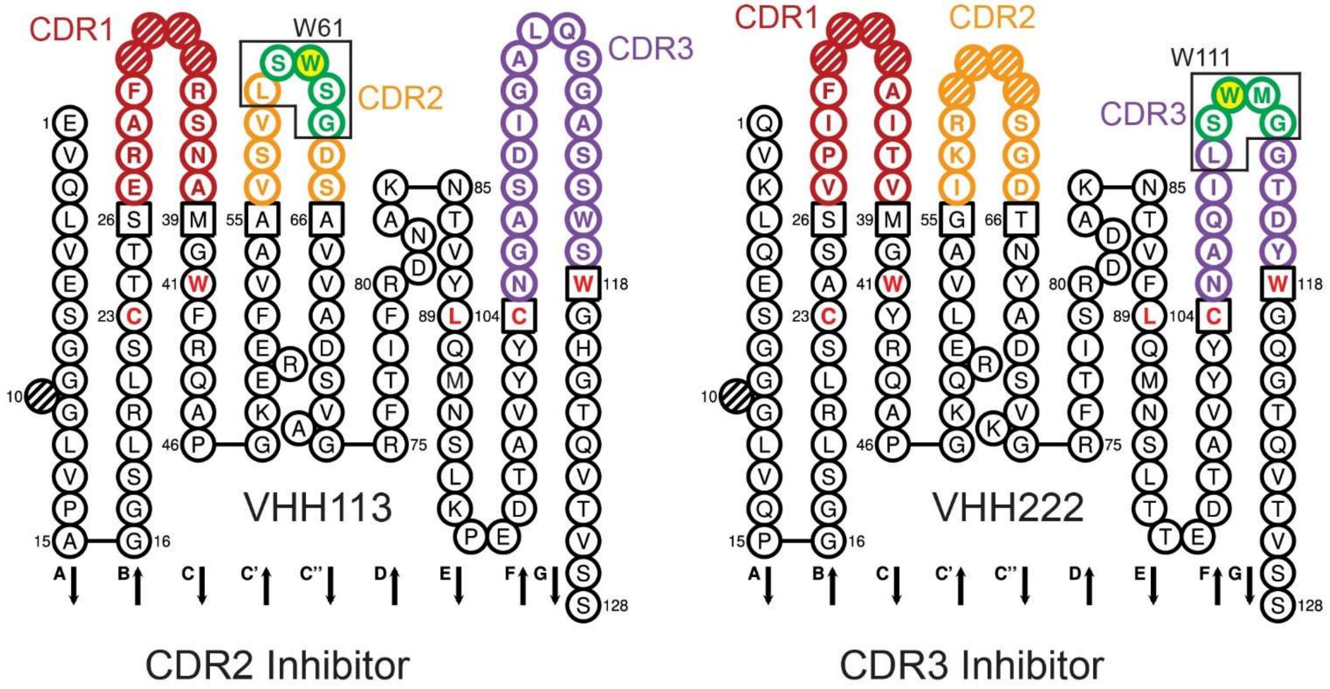

**Supplementary Figure 16. *Collier de Perles* diagrams highlight alternate locations of inhibitory loop and sequence variation in analogous CDRs between nanobody classes.** Analogous CDRs are markedly different between each class yet both recognize highly similar and overlapping Cif epitopes. However, the LSWXG motif (black box) containing the inhibitory Trp is found in VHH113 CDR2 or VHH222 CDR3. *Collier de Perles* diagrams of VHH113 and VHH222 were made using the IMGT/Collier-de-Perles web server [74,121-123] and re-colored to highlight the location of the Type 1 reverse turn (green/white) and the hydrophobic sidechain at position 111 (green/yellow). CDRs are colored as follows: CDR1 (red), CDR2 (orange), CDR3 (purple). Gapped positions are shown with hatch marks. Conserved FR residues are shown in red text and anchor positions appear in black boxes.

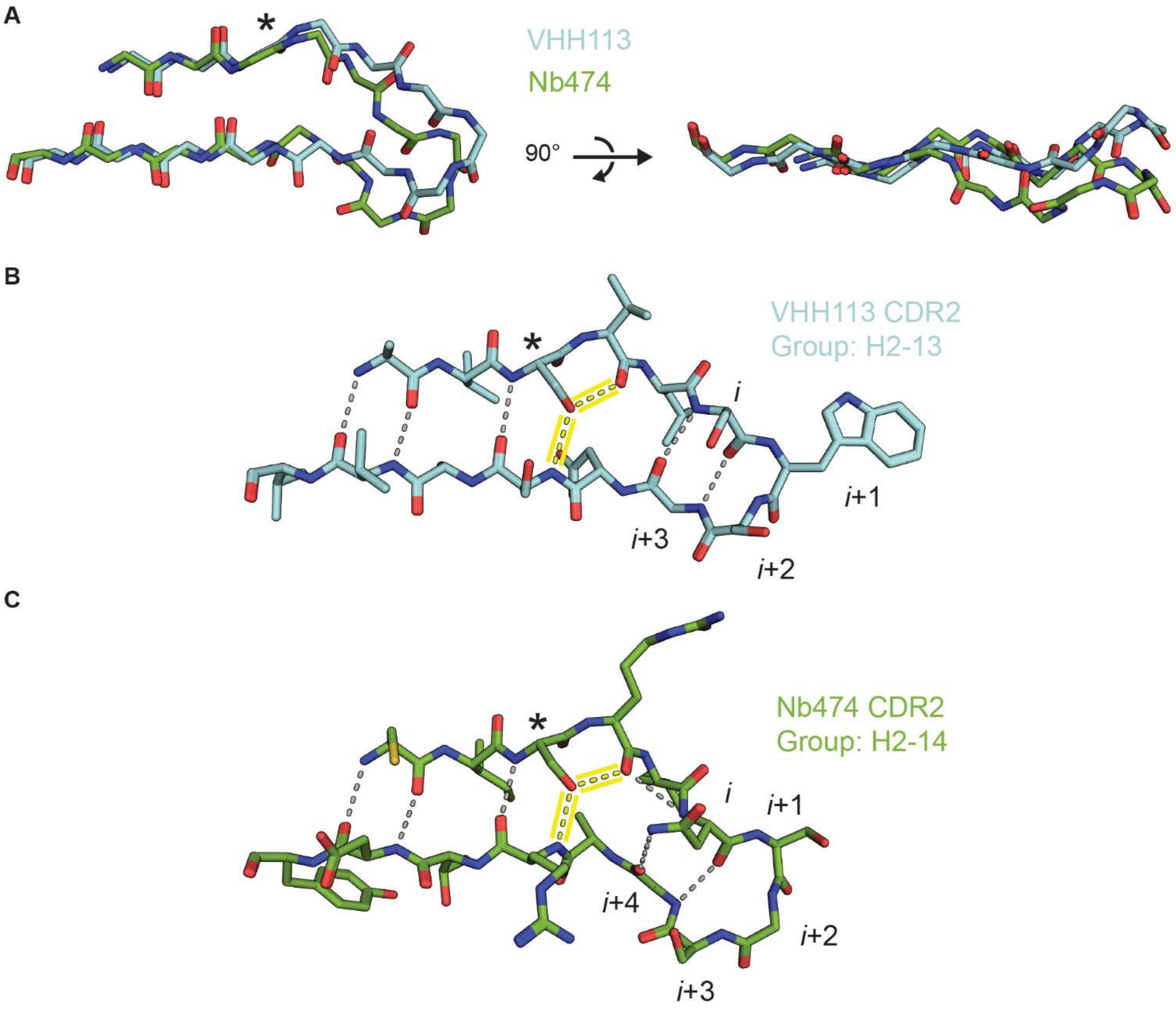

**Supplementary Figure 17. VHH113 CDR2 shares secondary-structure characteristics with Nb474 CDR2.** Following the convention established by [97], VHH113 CDR2 falls into the H2-13 loop cluster but is structurally dissimilar to representatives within the group. Instead, there is a strong resemblance to the CDR2 of Nb474 (pdb_00005o0w) which is in the H2-14 loop cluster. (A) Alignment of each CDR2 by main-chain atoms reveals a similar kink in the β-strand architecture. (B and C) In both structures, the serine at the third position in CDR2 (*) rotates toward the adjacent strand positioning the hydroxyl group just above the position where the main-chain carbonyl would be positioned in an archetypal anti-parallel β-strand. The atypical hydrogen-bond pattern (yellow highlight) is also accompanied with a register shift. (B) VHH113 CDR2 maintains a Type I reverse turn-like structure. (C) Nb474 CDR2 contains an extra amino acid resulting in a three-residue β-bulge. Residues in this loop are labeled as in a reverse turn for ease of comparison with VHH113 CDR2. VHH113, faded denim; Nb474, green. Non-carbon atoms are colored by type: oxygen, red; nitrogen, blue.

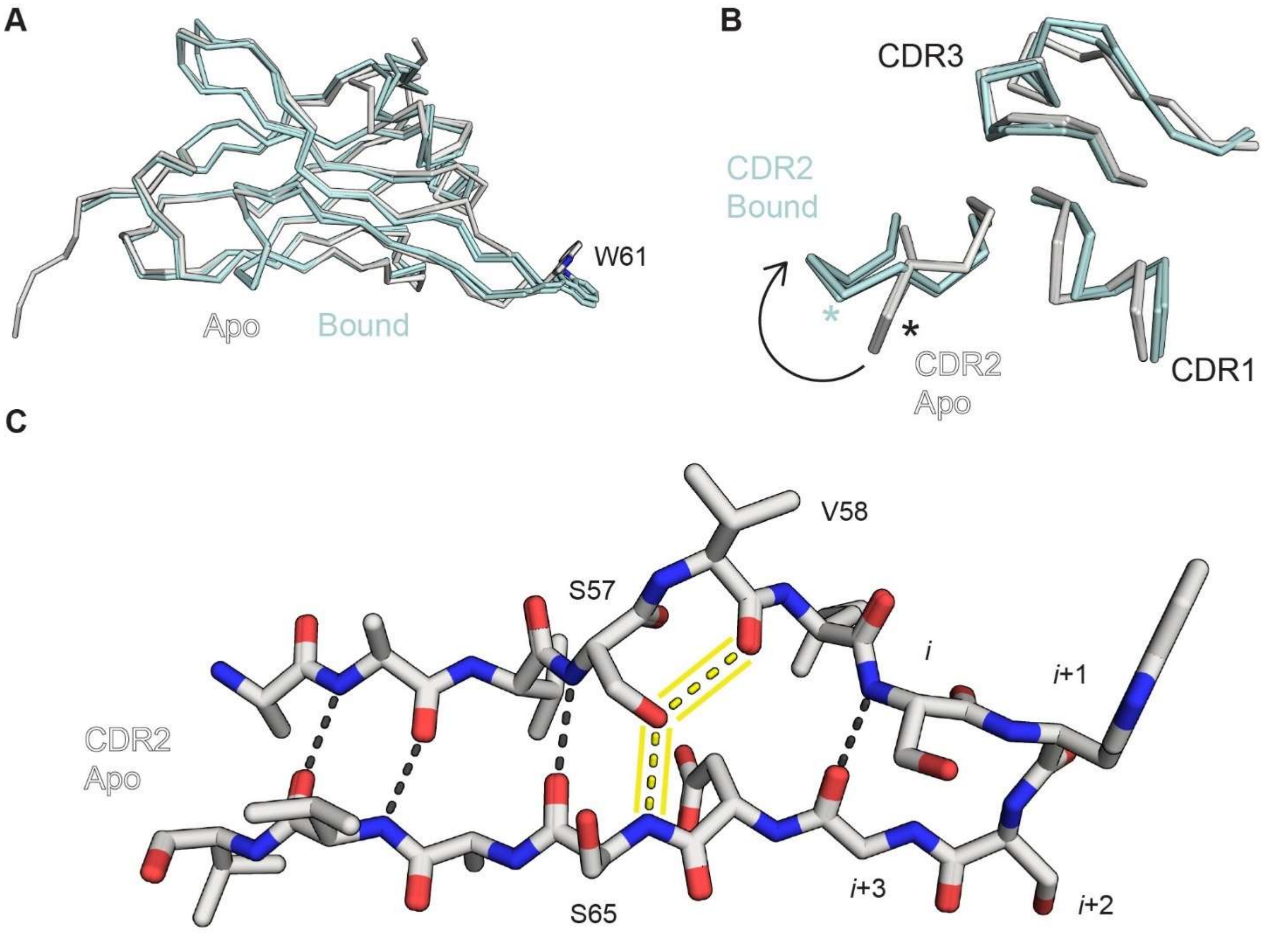

**Supplementary Figure 18. VHH113 CDR2 conformations in apo and Cif-bound states.** Structural alignment of the crystal structure of VHH113 in the apo state (pale nimbus white) to the Cif-bound state (faded denim) reveals different positions of CDR2 and the presence of the kinked β-strand architecture in the unbound state. (A) Side-view of VHH113 states with inhibitory Trp61 of CDR2 shown for orientation. (B) Face-on view of VHH113 CDRs highlighting alternate conformations of CDR2 in the free and bound structures, main-chain RMSD = 2.9 Å. Asterisks (*) denote the $C_\alpha$ position of Trp61. (C) Hydrogen bond pattern of VHH113 CDR2 in apo state with atypical hydrogen-bond pattern highlighted yellow. Non-carbon atoms are colored by type: oxygen, red; nitrogen, blue.

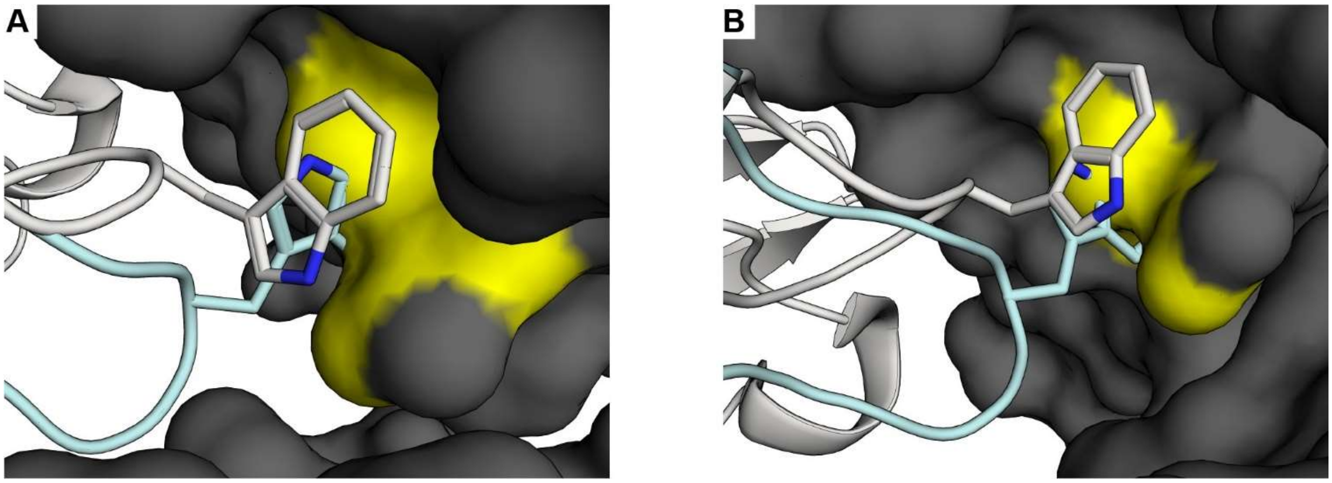

**Supplementary Figure 19. Potential influence of lattice packing on the VHH113 free form.** VHH113 crystallized with two molecules in the ASU. CDR2 extends outward and in the free form (pale nimbus white), it packs tightly against symmetry-related molecules in the crystal lattice (dark gray surface representation). The conformation of CDR2 in the Cif-bound state (faded denim) would clash with symmetry-related molecules (yellow). (A) Molecule A of the ASU. (B) Molecule B of the ASU. The inhibitory Trp61 that plugs the active-site entrance is shown as sticks. Non-carbon atoms are colored by type: nitrogen, blue.

**Supplementary Tables Below**

**Supplementary Table 1. Conditions for the crystallization of proteins.**

| Protein | Buffer | Concentration (mg/mL) | Well Solution | Drop Size (μL) | Drop Ratio (protein: well solution) | Cryoprotectant |
|---|---|---|---|---|---|---|
| VHH113 | 20 mM sodium phosphate, 20 mM NaCl, pH 7.4 | 5.4 | 17% (*w/v*) PEG 3350, 150 mM magnesium formate | 0.2 | 1:1 | 20% (*v/v*) glycerol |
| VHH222 | 20 mM Tris, 50 mM NaCl, pH 8.5 | 10 | 100 mM succinic acid/sodium phosphate monoboasic/glycine 2:7:7 (SPG), pH 6.0, 25% (*w/v*) PEG 1500 | 0.2 | 1:1 | 25% (*v/v*) glycerol |
| Cif:VHH101 | 20 mM sodium phosphate, 20 mM NaCl, pH 7.4 | 4.1 | 15% (*v/v*) isopropanol, 1 M ammonium citrate/ammonium hydroxide pH 8.5 | 0.2 | 1:1 | 25% (*v/v*) glycerol |
| Cif:VHH113 | 20 mM sodium phosphate, 20 mM NaCl, pH 7.4 | 3 – 3.5 | 200 mM potassium bromide, 200 mM potassium thiocyanate, 100 mM sodium acetate pH 5, 3% (*w/v*) gamma-PGA (Na+ form, LM), 10% (*w/v*) PEG 2000 MME | 0.2 | 1:1 | 25% (*v/v*) glycerol |
| Cif:VHH114 | 10 mM HEPES, 20 mM NaCl, pH 7.4 | 4.55 | 13.5 % (*w/v*) PEG6K, 2.5 % (*v/v*) ethylene glycol, 100 mM sodium citrate pH 3.5 | 4 | 1:1 | 22.5% (*v/v*) ethylene glycol |
| Cif:VHH219 | 20 mM Tris, 20 mM NaCl, pH 8.5 | 2.1 | 14.5% (*w/v*) PEG3350, 375 mM sodium malonate pH 4, 5 mM cobalt (II) chloride | 4 | 1:1 | 20% (*v/v*) glycerol |

| | | | | | | |
|---|---|---|---|---|---|---|
| Cif:VHH222 | 20 mM sodium phosphate, 50 mM NaCl, pH 7.4 | 14.5 | 150 mM ammonium sulfate, 100 mM TRIS pH 8, 15% (*w/v*) PEG 4000 | 0.2 | 1:2 | 20% (*v/v*) glycerol |

Buffer compositions and protein concentrations are listed for each protein complex. Well solution and cryoprotectant compositions are also shown in addition to drop size and volume.

**Supplementary Table 2. VHH Sequences and IMGT Nomenclature.**

| Nanobody | AA | IMGT | Seq Num |
|---|---|---|---|
| VHH101 | M | n/a | 1 |
| VHH101 | A | n/a | 2 |
| VHH101 | E | 1 | 3 |
| VHH101 | V | 2 | 4 |
| VHH101 | Q | 3 | 5 |
| VHH101 | L | 4 | 6 |
| VHH101 | V | 5 | 7 |
| VHH101 | E | 6 | 8 |
| VHH101 | S | 7 | 9 |
| VHH101 | G | 8 | 10 |
| VHH101 | G | 9 | 11 |
| VHH101 | G | 11 | 12 |
| VHH101 | L | 12 | 13 |
| VHH101 | V | 13 | 14 |
| VHH101 | Q | 14 | 15 |
| VHH101 | A | 15 | 16 |
| VHH101 | G | 16 | 17 |
| VHH101 | G | 17 | 18 |
| VHH101 | S | 18 | 19 |
| VHH101 | L | 19 | 20 |
| VHH101 | R | 20 | 21 |
| VHH101 | L | 21 | 22 |
| VHH101 | T | 22 | 23 |
| VHH101 | C | 23 | 24 |
| VHH101 | A | 24 | 25 |
| VHH101 | A | 25 | 26 |
| VHH101 | S | 26 | 27 |
| VHH101 | A | 27 | 28 |
| VHH101 | G | 28 | 29 |
| VHH101 | S | 29 | 30 |
| VHH101 | F | 30 | 31 |
| VHH101 | R | 35 | 32 |
| VHH101 | G | 36 | 33 |
| VHH101 | Y | 37 | 34 |
| VHH101 | A | 38 | 35 |
| VHH101 | M | 39 | 36 |
| VHH101 | G | 40 | 37 |
| VHH101 | W | 41 | 38 |

| Nanobody | AA | IMGT | Seq Num |
|---|---|---|---|
| VHH101 | F | 42 | 39 |
| VHH101 | R | 43 | 40 |
| VHH101 | Q | 44 | 41 |
| VHH101 | A | 45 | 42 |
| VHH101 | P | 46 | 43 |
| VHH101 | G | 47 | 44 |
| VHH101 | K | 48 | 45 |
| VHH101 | E | 49 | 46 |
| VHH101 | R | 50 | 47 |
| VHH101 | E | 51 | 48 |
| VHH101 | F | 52 | 49 |
| VHH101 | V | 53 | 50 |
| VHH101 | A | 54 | 51 |
| VHH101 | A | 55 | 52 |
| VHH101 | V | 56 | 53 |
| VHH101 | S | 57 | 54 |
| VHH101 | V | 58 | 55 |
| VHH101 | L | 59 | 56 |
| VHH101 | T | 60 | 57 |
| VHH101 | W | 61 | 58 |
| VHH101 | S | 62 | 59 |
| VHH101 | G | 63 | 60 |
| VHH101 | D | 64 | 61 |
| VHH101 | S | 65 | 62 |
| VHH101 | T | 66 | 63 |
| VHH101 | N | 67 | 64 |
| VHH101 | I | 68 | 65 |
| VHH101 | A | 69 | 66 |
| VHH101 | D | 70 | 67 |
| VHH101 | S | 71 | 68 |
| VHH101 | V | 72 | 69 |
| VHH101 | K | 73 | 70 |
| VHH101 | G | 74 | 71 |
| VHH101 | R | 75 | 72 |
| VHH101 | F | 76 | 73 |
| VHH101 | T | 77 | 74 |
| VHH101 | I | 78 | 75 |
| VHH101 | F | 79 | 76 |
| VHH101 | R | 80 | 77 |
| VHH101 | D | 81 | 78 |

| Nanobody | AA | IMGT | Seq Num |
|---|---|---|---|
| VHH101 | T | 82 | 79 |
| VHH101 | A | 83 | 80 |
| VHH101 | K | 84 | 81 |
| VHH101 | N | 85 | 82 |
| VHH101 | T | 86 | 83 |
| VHH101 | V | 87 | 84 |
| VHH101 | Y | 88 | 85 |
| VHH101 | L | 89 | 86 |
| VHH101 | Q | 90 | 87 |
| VHH101 | M | 91 | 88 |
| VHH101 | N | 92 | 89 |
| VHH101 | S | 93 | 90 |
| VHH101 | L | 94 | 91 |
| VHH101 | K | 95 | 92 |
| VHH101 | P | 96 | 93 |
| VHH101 | E | 97 | 94 |
| VHH101 | D | 98 | 95 |
| VHH101 | T | 99 | 96 |
| VHH101 | A | 100 | 97 |
| VHH101 | V | 101 | 98 |
| VHH101 | Y | 102 | 99 |
| VHH101 | Y | 103 | 100 |
| VHH101 | C | 104 | 101 |
| VHH101 | N | 105 | 102 |
| VHH101 | G | 106 | 103 |
| VHH101 | A | 107 | 104 |
| VHH101 | S | 108 | 105 |
| VHH101 | E | 109 | 106 |
| VHH101 | I | 110 | 107 |
| VHH101 | G | 111 | 108 |
| VHH101 | A | 111.1 | 109 |
| VHH101 | L | 111.2 | 110 |
| VHH101 | Q | 112.2 | 111 |
| VHH101 | S | 112.1 | 112 |
| VHH101 | G | 112 | 113 |
| VHH101 | A | 113 | 114 |
| VHH101 | S | 114 | 115 |
| VHH101 | L | 115 | 116 |
| VHH101 | W | 116 | 117 |
| VHH101 | S | 117 | 118 |

| Nanobody | AA | IMGT | Seq Num |
|---|---|---|---|
| VHH101 | W | 118 | 119 |
| VHH101 | G | 119 | 120 |
| VHH101 | Q | 120 | 121 |
| VHH101 | G | 121 | 122 |
| VHH101 | T | 122 | 123 |
| VHH101 | Q | 123 | 124 |
| VHH101 | V | 124 | 125 |
| VHH101 | T | 125 | 126 |
| VHH101 | V | 126 | 127 |
| VHH101 | S | 127 | 128 |
| VHH101 | S | 128 | 129 |
| VHH101 | G | n/a | 130 |
| VHH101 | Q | n/a | 131 |
| VHH101 | A | n/a | 132 |
| VHH101 | G | n/a | 133 |
| VHH101 | Q | n/a | 134 |
| VHH113 | M | n/a | 1 |
| VHH113 | A | n/a | 2 |
| VHH113 | E | 1 | 3 |
| VHH113 | V | 2 | 4 |
| VHH113 | Q | 3 | 5 |
| VHH113 | L | 4 | 6 |
| VHH113 | V | 5 | 7 |
| VHH113 | E | 6 | 8 |
| VHH113 | S | 7 | 9 |
| VHH113 | G | 8 | 10 |
| VHH113 | G | 9 | 11 |
| VHH113 | G | 11 | 12 |
| VHH113 | L | 12 | 13 |
| VHH113 | V | 13 | 14 |
| VHH113 | P | 14 | 15 |
| VHH113 | A | 15 | 16 |
| VHH113 | G | 16 | 17 |
| VHH113 | G | 17 | 18 |
| VHH113 | S | 18 | 19 |
| VHH113 | L | 19 | 20 |
| VHH113 | R | 20 | 21 |
| VHH113 | L | 21 | 22 |
| VHH113 | S | 22 | 23 |

| Nanobody | AA | IMGT | Seq Num |
|---|---|---|---|
| VHH113 | C | 23 | 24 |
| VHH113 | T | 24 | 25 |
| VHH113 | T | 25 | 26 |
| VHH113 | S | 26 | 27 |
| VHH113 | E | 27 | 28 |
| VHH113 | R | 28 | 29 |
| VHH113 | A | 29 | 30 |
| VHH113 | F | 30 | 31 |
| VHH113 | R | 35 | 32 |
| VHH113 | S | 36 | 33 |
| VHH113 | N | 37 | 34 |
| VHH113 | A | 38 | 35 |
| VHH113 | M | 39 | 36 |
| VHH113 | G | 40 | 37 |
| VHH113 | W | 41 | 38 |
| VHH113 | F | 42 | 39 |
| VHH113 | R | 43 | 40 |
| VHH113 | Q | 44 | 41 |
| VHH113 | A | 45 | 42 |
| VHH113 | P | 46 | 43 |
| VHH113 | G | 47 | 44 |
| VHH113 | K | 48 | 45 |
| VHH113 | E | 49 | 46 |
| VHH113 | R | 50 | 47 |
| VHH113 | E | 51 | 48 |
| VHH113 | F | 52 | 49 |
| VHH113 | V | 53 | 50 |
| VHH113 | A | 54 | 51 |
| VHH113 | A | 55 | 52 |
| VHH113 | V | 56 | 53 |
| VHH113 | S | 57 | 54 |
| VHH113 | V | 58 | 55 |
| VHH113 | L | 59 | 56 |
| VHH113 | S | 60 | 57 |
| VHH113 | W | 61 | 58 |
| VHH113 | S | 62 | 59 |
| VHH113 | G | 63 | 60 |
| VHH113 | D | 64 | 61 |
| VHH113 | S | 65 | 62 |
| VHH113 | A | 66 | 63 |

| Nanobody | AA | IMGT | Seq Num |
|---|---|---|---|
| VHH113 | V | 67 | 64 |
| VHH113 | V | 68 | 65 |
| VHH113 | A | 69 | 66 |
| VHH113 | D | 70 | 67 |
| VHH113 | S | 71 | 68 |
| VHH113 | V | 72 | 69 |
| VHH113 | A | 73 | 70 |
| VHH113 | G | 74 | 71 |
| VHH113 | R | 75 | 72 |
| VHH113 | F | 76 | 73 |
| VHH113 | T | 77 | 74 |
| VHH113 | I | 78 | 75 |
| VHH113 | F | 79 | 76 |
| VHH113 | R | 80 | 77 |
| VHH113 | D | 81 | 78 |
| VHH113 | N | 82 | 79 |
| VHH113 | A | 83 | 80 |
| VHH113 | K | 84 | 81 |
| VHH113 | N | 85 | 82 |
| VHH113 | T | 86 | 83 |
| VHH113 | V | 87 | 84 |
| VHH113 | Y | 88 | 85 |
| VHH113 | L | 89 | 86 |
| VHH113 | Q | 90 | 87 |
| VHH113 | M | 91 | 88 |
| VHH113 | N | 92 | 89 |
| VHH113 | S | 93 | 90 |
| VHH113 | L | 94 | 91 |
| VHH113 | K | 95 | 92 |
| VHH113 | P | 96 | 93 |
| VHH113 | E | 97 | 94 |
| VHH113 | D | 98 | 95 |
| VHH113 | T | 99 | 96 |
| VHH113 | A | 100 | 97 |
| VHH113 | V | 101 | 98 |
| VHH113 | Y | 102 | 99 |
| VHH113 | Y | 103 | 100 |
| VHH113 | C | 104 | 101 |
| VHH113 | N | 105 | 102 |
| VHH113 | G | 106 | 103 |

| Nanobody | AA | IMGT | Seq Num |
|---|---|---|---|
| VHH113 | A | 107 | 104 |
| VHH113 | S | 108 | 105 |
| VHH113 | D | 109 | 106 |
| VHH113 | I | 110 | 107 |
| VHH113 | G | 111 | 108 |
| VHH113 | A | 111.1 | 109 |
| VHH113 | L | 111.2 | 110 |
| VHH113 | Q | 112.2 | 111 |
| VHH113 | S | 112.1 | 112 |
| VHH113 | G | 112 | 113 |
| VHH113 | A | 113 | 114 |
| VHH113 | S | 114 | 115 |
| VHH113 | S | 115 | 116 |
| VHH113 | W | 116 | 117 |
| VHH113 | S | 117 | 118 |
| VHH113 | W | 118 | 119 |
| VHH113 | G | 119 | 120 |
| VHH113 | H | 120 | 121 |
| VHH113 | G | 121 | 122 |
| VHH113 | T | 122 | 123 |
| VHH113 | Q | 123 | 124 |
| VHH113 | V | 124 | 125 |
| VHH113 | T | 125 | 126 |
| VHH113 | V | 126 | 127 |
| VHH113 | S | 127 | 128 |
| VHH113 | S | 128 | 129 |
| VHH113 | G | n/a | 130 |
| VHH113 | Q | n/a | 131 |
| VHH113 | A | n/a | 132 |
| VHH113 | G | n/a | 133 |
| VHH113 | Q | n/a | 134 |
| VHH114 | M | n/a | 1 |
| VHH114 | A | n/a | 2 |
| VHH114 | Q | 1 | 3 |
| VHH114 | V | 2 | 4 |
| VHH114 | K | 3 | 5 |
| VHH114 | L | 4 | 6 |
| VHH114 | Q | 5 | 7 |
| VHH114 | E | 6 | 8 |

| Nanobody | AA | IMGT | Seq Num |
|---|---|---|---|
| VHH114 | S | 7 | 9 |
| VHH114 | G | 8 | 10 |
| VHH114 | G | 9 | 11 |
| VHH114 | G | 11 | 12 |
| VHH114 | L | 12 | 13 |
| VHH114 | V | 13 | 14 |
| VHH114 | Q | 14 | 15 |
| VHH114 | P | 15 | 16 |
| VHH114 | G | 16 | 17 |
| VHH114 | E | 17 | 18 |
| VHH114 | S | 18 | 19 |
| VHH114 | L | 19 | 20 |
| VHH114 | T | 20 | 21 |
| VHH114 | L | 21 | 22 |
| VHH114 | S | 22 | 23 |
| VHH114 | C | 23 | 24 |
| VHH114 | A | 24 | 25 |
| VHH114 | V | 25 | 26 |
| VHH114 | S | 26 | 27 |
| VHH114 | V | 27 | 28 |
| VHH114 | R | 28 | 29 |
| VHH114 | L | 29 | 30 |
| VHH114 | S | 30 | 31 |
| VHH114 | G | 35 | 32 |
| VHH114 | I | 36 | 33 |
| VHH114 | T | 37 | 34 |
| VHH114 | T | 38 | 35 |
| VHH114 | M | 39 | 36 |
| VHH114 | G | 40 | 37 |
| VHH114 | W | 41 | 38 |
| VHH114 | Y | 42 | 39 |
| VHH114 | R | 43 | 40 |
| VHH114 | Q | 44 | 41 |
| VHH114 | A | 45 | 42 |
| VHH114 | P | 46 | 43 |
| VHH114 | G | 47 | 44 |
| VHH114 | K | 48 | 45 |
| VHH114 | Q | 49 | 46 |
| VHH114 | R | 50 | 47 |
| VHH114 | E | 51 | 48 |

| Nanobody | AA | IMGT | Seq Num |
|---|---|---|---|
| VHH114 | M | 52 | 49 |
| VHH114 | V | 53 | 50 |
| VHH114 | A | 54 | 51 |
| VHH114 | S | 55 | 52 |
| VHH114 | I | 56 | 53 |
| VHH114 | S | 57 | 54 |
| VHH114 | R | 58 | 55 |
| VHH114 | G | 63 | 56 |
| VHH114 | G | 64 | 57 |
| VHH114 | S | 65 | 58 |
| VHH114 | T | 66 | 59 |
| VHH114 | V | 67 | 60 |
| VHH114 | Y | 68 | 61 |
| VHH114 | L | 69 | 62 |
| VHH114 | D | 70 | 63 |
| VHH114 | S | 71 | 64 |
| VHH114 | V | 72 | 65 |
| VHH114 | K | 73 | 66 |
| VHH114 | G | 74 | 67 |
| VHH114 | R | 75 | 68 |
| VHH114 | F | 76 | 69 |
| VHH114 | T | 77 | 70 |
| VHH114 | V | 78 | 71 |
| VHH114 | S | 79 | 72 |
| VHH114 | R | 80 | 73 |
| VHH114 | D | 81 | 74 |
| VHH114 | N | 82 | 75 |
| VHH114 | T | 83 | 76 |
| VHH114 | K | 84 | 77 |
| VHH114 | N | 85 | 78 |
| VHH114 | T | 86 | 79 |
| VHH114 | V | 87 | 80 |
| VHH114 | K | 88 | 81 |
| VHH114 | L | 89 | 82 |
| VHH114 | Q | 90 | 83 |
| VHH114 | M | 91 | 84 |
| VHH114 | N | 92 | 85 |
| VHH114 | S | 93 | 86 |
| VHH114 | L | 94 | 87 |
| VHH114 | K | 95 | 88 |

| Nanobody | AA | IMGT | Seq Num |
|---|---|---|---|
| VHH114 | P | 96 | 89 |
| VHH114 | E | 97 | 90 |
| VHH114 | D | 98 | 91 |
| VHH114 | T | 99 | 92 |
| VHH114 | A | 100 | 93 |
| VHH114 | I | 101 | 94 |
| VHH114 | Y | 102 | 95 |
| VHH114 | Y | 103 | 96 |
| VHH114 | C | 104 | 97 |
| VHH114 | N | 105 | 98 |
| VHH114 | A | 106 | 99 |
| VHH114 | K | 107 | 100 |
| VHH114 | I | 108 | 101 |
| VHH114 | L | 109 | 102 |
| VHH114 | L | 110 | 103 |
| VHH114 | V | 111 | 104 |
| VHH114 | A | 112 | 105 |
| VHH114 | S | 113 | 106 |
| VHH114 | T | 114 | 107 |
| VHH114 | D | 115 | 108 |
| VHH114 | D | 116 | 109 |
| VHH114 | Y | 117 | 110 |
| VHH114 | W | 118 | 111 |
| VHH114 | G | 119 | 112 |
| VHH114 | Q | 120 | 113 |
| VHH114 | G | 121 | 114 |
| VHH114 | T | 122 | 115 |
| VHH114 | Q | 123 | 116 |
| VHH114 | V | 124 | 117 |
| VHH114 | T | 125 | 118 |
| VHH114 | V | 126 | 119 |
| VHH114 | S | 127 | 120 |
| VHH114 | S | 128 | 121 |
| VHH114 | G | n/a | 122 |
| VHH114 | Q | n/a | 123 |
| VHH114 | A | n/a | 124 |
| VHH114 | G | n/a | 125 |
| VHH114 | Q | n/a | 126 |
| VHH219 | M | n/a | 1 |

| Nanobody | AA | IMGT | Seq Num |
|---|---|---|---|
| VHH219 | A | n/a | 2 |
| VHH219 | E | 1 | 3 |
| VHH219 | V | 2 | 4 |
| VHH219 | Q | 3 | 5 |
| VHH219 | L | 4 | 6 |
| VHH219 | V | 5 | 7 |
| VHH219 | E | 6 | 8 |
| VHH219 | S | 7 | 9 |
| VHH219 | G | 8 | 10 |
| VHH219 | G | 9 | 11 |
| VHH219 | G | 11 | 12 |
| VHH219 | L | 12 | 13 |
| VHH219 | V | 13 | 14 |
| VHH219 | Q | 14 | 15 |
| VHH219 | P | 15 | 16 |
| VHH219 | G | 16 | 17 |
| VHH219 | G | 17 | 18 |
| VHH219 | S | 18 | 19 |
| VHH219 | L | 19 | 20 |
| VHH219 | R | 20 | 21 |
| VHH219 | L | 21 | 22 |
| VHH219 | S | 22 | 23 |
| VHH219 | C | 23 | 24 |
| VHH219 | T | 24 | 25 |
| VHH219 | T | 25 | 26 |
| VHH219 | S | 26 | 27 |
| VHH219 | T | 27 | 28 |
| VHH219 | S | 28 | 29 |
| VHH219 | L | 29 | 30 |
| VHH219 | F | 30 | 31 |
| VHH219 | S | 35 | 32 |
| VHH219 | I | 36 | 33 |
| VHH219 | T | 37 | 34 |
| VHH219 | T | 38 | 35 |
| VHH219 | M | 39 | 36 |
| VHH219 | G | 40 | 37 |
| VHH219 | W | 41 | 38 |
| VHH219 | Y | 42 | 39 |
| VHH219 | R | 43 | 40 |
| VHH219 | Q | 44 | 41 |

| Nanobody | AA | IMGT | Seq Num |
|---|---|---|---|
| VHH219 | A | 45 | 42 |
| VHH219 | P | 46 | 43 |
| VHH219 | G | 47 | 44 |
| VHH219 | K | 48 | 45 |
| VHH219 | Q | 49 | 46 |
| VHH219 | R | 50 | 47 |
| VHH219 | E | 51 | 48 |
| VHH219 | L | 52 | 49 |
| VHH219 | V | 53 | 50 |
| VHH219 | A | 54 | 51 |
| VHH219 | S | 55 | 52 |
| VHH219 | I | 56 | 53 |
| VHH219 | K | 57 | 54 |
| VHH219 | R | 58 | 55 |
| VHH219 | G | 63 | 56 |
| VHH219 | G | 64 | 57 |
| VHH219 | G | 65 | 58 |
| VHH219 | T | 66 | 59 |
| VHH219 | N | 67 | 60 |
| VHH219 | Y | 68 | 61 |
| VHH219 | A | 69 | 62 |
| VHH219 | D | 70 | 63 |
| VHH219 | S | 71 | 64 |
| VHH219 | M | 72 | 65 |
| VHH219 | K | 73 | 66 |
| VHH219 | G | 74 | 67 |
| VHH219 | R | 75 | 68 |
| VHH219 | F | 76 | 69 |
| VHH219 | T | 77 | 70 |
| VHH219 | I | 78 | 71 |
| VHH219 | S | 79 | 72 |
| VHH219 | R | 80 | 73 |
| VHH219 | D | 81 | 74 |
| VHH219 | N | 82 | 75 |
| VHH219 | A | 83 | 76 |
| VHH219 | R | 84 | 77 |
| VHH219 | N | 85 | 78 |
| VHH219 | T | 86 | 79 |
| VHH219 | V | 87 | 80 |
| VHH219 | F | 88 | 81 |

| Nanobody | AA | IMGT | Seq Num |
|---|---|---|---|
| VHH219 | L | 89 | 82 |
| VHH219 | E | 90 | 83 |
| VHH219 | M | 91 | 84 |
| VHH219 | N | 92 | 85 |
| VHH219 | N | 93 | 86 |
| VHH219 | L | 94 | 87 |
| VHH219 | T | 95 | 88 |
| VHH219 | T | 96 | 89 |
| VHH219 | E | 97 | 90 |
| VHH219 | D | 98 | 91 |
| VHH219 | T | 99 | 92 |
| VHH219 | A | 100 | 93 |
| VHH219 | V | 101 | 94 |
| VHH219 | Y | 102 | 95 |
| VHH219 | Y | 103 | 96 |
| VHH219 | C | 104 | 97 |
| VHH219 | N | 105 | 98 |
| VHH219 | A | 106 | 99 |
| VHH219 | A | 107 | 100 |
| VHH219 | I | 108 | 101 |
| VHH219 | L | 109 | 102 |
| VHH219 | A | 110 | 103 |
| VHH219 | Y | 111 | 104 |
| VHH219 | T | 112.1 | 105 |
| VHH219 | G | 112 | 106 |
| VHH219 | E | 113 | 107 |
| VHH219 | V | 114 | 108 |
| VHH219 | T | 115 | 109 |
| VHH219 | N | 116 | 110 |
| VHH219 | Y | 117 | 111 |
| VHH219 | W | 118 | 112 |
| VHH219 | G | 119 | 113 |
| VHH219 | Q | 120 | 114 |
| VHH219 | G | 121 | 115 |
| VHH219 | T | 122 | 116 |
| VHH219 | Q | 123 | 117 |
| VHH219 | V | 124 | 118 |
| VHH219 | T | 125 | 119 |
| VHH219 | V | 126 | 120 |
| VHH219 | S | 127 | 121 |

| Nanobody | AA | IMGT | Seq Num |
|---|---|---|---|
| VHH219 | S | 128 | 122 |
| VHH219 | G | n/a | 123 |
| VHH219 | Q | n/a | 124 |
| VHH219 | A | n/a | 125 |
| VHH219 | G | n/a | 126 |
| VHH219 | Q | n/a | 127 |
| VHH222 | M | n/a | 1 |
| VHH222 | A | n/a | 2 |
| VHH222 | Q | 1 | 3 |
| VHH222 | V | 2 | 4 |
| VHH222 | K | 3 | 5 |
| VHH222 | L | 4 | 6 |
| VHH222 | Q | 5 | 7 |
| VHH222 | E | 6 | 8 |
| VHH222 | S | 7 | 9 |
| VHH222 | G | 8 | 10 |
| VHH222 | G | 9 | 11 |
| VHH222 | G | 11 | 12 |
| VHH222 | L | 12 | 13 |
| VHH222 | V | 13 | 14 |
| VHH222 | Q | 14 | 15 |
| VHH222 | P | 15 | 16 |
| VHH222 | G | 16 | 17 |
| VHH222 | G | 17 | 18 |
| VHH222 | S | 18 | 19 |
| VHH222 | L | 19 | 20 |
| VHH222 | R | 20 | 21 |
| VHH222 | L | 21 | 22 |
| VHH222 | S | 22 | 23 |
| VHH222 | C | 23 | 24 |
| VHH222 | A | 24 | 25 |
| VHH222 | S | 25 | 26 |
| VHH222 | S | 26 | 27 |
| VHH222 | V | 27 | 28 |
| VHH222 | P | 28 | 29 |
| VHH222 | I | 29 | 30 |
| VHH222 | F | 30 | 31 |
| VHH222 | A | 35 | 32 |
| VHH222 | I | 36 | 33 |

| Nanobody | AA | IMGT | Seq Num |
|---|---|---|---|
| VHH222 | T | 37 | 34 |
| VHH222 | V | 38 | 35 |
| VHH222 | M | 39 | 36 |
| VHH222 | G | 40 | 37 |
| VHH222 | W | 41 | 38 |
| VHH222 | Y | 42 | 39 |
| VHH222 | R | 43 | 40 |
| VHH222 | Q | 44 | 41 |
| VHH222 | A | 45 | 42 |
| VHH222 | P | 46 | 43 |
| VHH222 | G | 47 | 44 |
| VHH222 | K | 48 | 45 |
| VHH222 | Q | 49 | 46 |
| VHH222 | R | 50 | 47 |
| VHH222 | E | 51 | 48 |
| VHH222 | L | 52 | 49 |
| VHH222 | V | 53 | 50 |
| VHH222 | A | 54 | 51 |
| VHH222 | G | 55 | 52 |
| VHH222 | I | 56 | 53 |
| VHH222 | K | 57 | 54 |
| VHH222 | R | 58 | 55 |
| VHH222 | S | 63 | 56 |
| VHH222 | G | 64 | 57 |
| VHH222 | D | 65 | 58 |
| VHH222 | T | 66 | 59 |
| VHH222 | N | 67 | 60 |
| VHH222 | Y | 68 | 61 |
| VHH222 | A | 69 | 62 |
| VHH222 | D | 70 | 63 |
| VHH222 | S | 71 | 64 |
| VHH222 | V | 72 | 65 |
| VHH222 | K | 73 | 66 |
| VHH222 | G | 74 | 67 |
| VHH222 | R | 75 | 68 |
| VHH222 | F | 76 | 69 |
| VHH222 | T | 77 | 70 |
| VHH222 | I | 78 | 71 |
| VHH222 | S | 79 | 72 |
| VHH222 | R | 80 | 73 |

| Nanobody | AA | IMGT | Seq Num |
|---|---|---|---|
| VHH222 | D | 81 | 74 |
| VHH222 | D | 82 | 75 |
| VHH222 | A | 83 | 76 |
| VHH222 | K | 84 | 77 |
| VHH222 | N | 85 | 78 |
| VHH222 | T | 86 | 79 |
| VHH222 | V | 87 | 80 |
| VHH222 | F | 88 | 81 |
| VHH222 | L | 89 | 82 |
| VHH222 | Q | 90 | 83 |
| VHH222 | M | 91 | 84 |
| VHH222 | N | 92 | 85 |
| VHH222 | S | 93 | 86 |
| VHH222 | L | 94 | 87 |
| VHH222 | T | 95 | 88 |
| VHH222 | T | 96 | 89 |
| VHH222 | E | 97 | 90 |
| VHH222 | D | 98 | 91 |
| VHH222 | T | 99 | 92 |
| VHH222 | A | 100 | 93 |
| VHH222 | V | 101 | 94 |
| VHH222 | Y | 102 | 95 |
| VHH222 | Y | 103 | 96 |
| VHH222 | C | 104 | 97 |
| VHH222 | N | 105 | 98 |
| VHH222 | A | 106 | 99 |
| VHH222 | Q | 107 | 100 |
| VHH222 | I | 108 | 101 |
| VHH222 | L | 109 | 102 |
| VHH222 | S | 110 | 103 |
| VHH222 | W | 111 | 104 |
| VHH222 | M | 112 | 105 |
| VHH222 | G | 113 | 106 |
| VHH222 | G | 114 | 107 |
| VHH222 | T | 115 | 108 |
| VHH222 | D | 116 | 109 |
| VHH222 | Y | 117 | 110 |
| VHH222 | W | 118 | 111 |
| VHH222 | G | 119 | 112 |
| VHH222 | Q | 120 | 113 |

| Nanobody | AA | IMGT | Seq Num |
|---|---|---|---|
| VHH222 | G | 121 | 114 |
| VHH222 | T | 122 | 115 |
| VHH222 | Q | 123 | 116 |
| VHH222 | V | 124 | 117 |
| VHH222 | T | 125 | 118 |
| VHH222 | V | 126 | 119 |
| VHH222 | S | 127 | 120 |
| VHH222 | S | 128 | 121 |
| VHH222 | G | n/a | 122 |
| VHH222 | Q | n/a | 123 |
| VHH222 | A | n/a | 124 |
| VHH222 | G | n/a | 125 |
| VHH222 | Q | n/a | 126 |

Sequences for all VHHs with numbering scheme following IMGT unique numbering. Column headers are as follows: Nanobody, VHH identifier; AA, amino acid one-letter code; IMGT, residue position following IMGT nomenclature; Seq Num, residue position in polypeptide chain.

**Supplementary Table 3. Data collection, reduction, and refinement statistics.**

| | VHH113 | VHH222 | Cif:VHH101 | Cif:VHH113 |
|---|---|---|---|---|
| **Data Collection** | | | | |
| Beamline | NSLS-II 17-ID-2 | NSLS-II 17-ID-2 | NSLS-II 17-ID-2 | NSLS-II 17-ID-2 |
| Wavelength (Å) | 0.979106 | 0.978636 | 0.979339 | 0.979339 |
| Space Group | $P\ 2_12_12_1$ | $P\ 3_121$ | $C\ 2$ | $C\ 222_1$ |
| Unit cell parameters | | | | |
| *a, b, c* (Å) | 51.1, 51.6, 76.0 | 58.0, 58.0, 75.9 | 189.0, 92.0, 151.2 | 115.2, 185.9, 106.8 |
| $\alpha, \beta, \gamma$ (°) | 90, 90, 90 | 90, 90, 120 | 90, 92.5, 90 | 90, 90, 90 |
| Resolution[a] (Å) | 42.4 - 1.10 (1.14 - 1.10) | 41.9 - 1.90 (1.97 - 1.90) | 45.3 - 2.00 (2.07 - 2.00) | 46.5 - 1.70 (1.76 - 1.70) |
| $R_{meas}$[b] (%) | 10.2 (157.8) | 7.9 (137.9) | 9.6 (105.4) | 11.6 (178.8) |
| $R_{p.i.m.}$ (%) | 4.1 (80.8) | 2.5 (48.4) | 4.9 (53.0) | 4.5 (69.1) |
| $CC_{1/2}$[c] (%) | 99.9 (52.4) | 99.9 (83.8) | 99.7 (53.9) | 99.8 (43.9) |
| $I/\sigma_I$ | 8.89 (0.78) | 16.76 (1.58) | 10.57 (2.02) | 11.80 (1.26) |
| Completeness (%) | 99.2 (94.2) | 99.5 (97.4) | 94.5 (97.8) | 99.1 (98.7) |
| Redundancy | 6.1 (3.5) | 9.5 (7.6) | 3.6 (3.6) | 6.7 (6.6) |
| **Refinement** | | | | |
| #Total reflections | 81532 (7645) | 12013 (1139) | 165131 (16961) | 124296 (12261) |
| Test-set reflections | 4114 (388) | 598 (60) | 8270 (797) | 6228 (593) |
| $R_{work}$[d]/$R_{free}$[e] (%) | 19.3/21.3 | 21.2/22.4 | 18.4/20.6 | 17.6/19.9 |
| Number of atoms: | | | | |
| Protein | 2040 | 881 | 19650 | 6620 |
| Water | 320 | 58 | 922 | 784 |
| Ramachandran Plot[f] (%) | 98.5/1.5/0.0 | 99.1/0.9/0.0 | 98.1/1.9/0.0 | 98.1/1.9/0.0 |
| $B_{average}$ (Å$^2$) | 15.34 | 42.92 | 40.94 | 29.61 |
| Protein | 13.67 | 42.67 | 40.94 | 28.45 |
| Water | 26.00 | 46.67 | 40.97 | 39.46 |
| Bond length RMSD (Å) | 0.006 | 0.010 | 0.009 | 0.006 |
| Bond angle RMSD (°) | 0.86 | 0.82 | 0.97 | 0.84 |
| **PDB ID** | pdb_00008e2n | pdb_00008e1c | pdb_00008evd | pdb_00008f6u |

| | Cif:VHH114 | Cif:VHH219 | Cif:VHH222 |
|---|---|---|---|
| **Data Collection** | | | |
| Beamline | NSLS-II 17-ID-1 | NSLS-II 17-ID-2 | NSLS-II 17-ID-2 |
| Wavelength (Å) | 0.920112 | 0.979261 | 0.97933 |
| Space Group | $P\,6_322$ | $P\,2_1$ | $P\,2_1$ |
| Unit cell parameters | | | |
| *a, b, c* (Å) | 163.5, 163.5, 75.0 | 76.2, 162.6, 106.3 | 76.2, 164.3, 107.3 |
| $\alpha, \beta, \gamma$ (°) | 90, 90, 120 | 90, 95.4, 90 | 90, 97.5, 90 |
| Resolution[a] (Å) | 27.1 - 1.85 (1.92 - 1.85) | 41.6 - 2.40 (2.49 - 2.40) | 47.4 - 2.40 (2.49 - 2.40) |
| $R_{meas}$[b] (%) | 14.3 (246.5) | 8.7 (130.9) | 19.1 (118.8) |
| $R_{p.i.m.}$ (%) | 3.2 (53.9) | 4.6 (69.2) | 10.0 (61.7) |
| $CC_{1/2}$[c] (%) | 99.9 (61.8) | 99.8 (50.1) | 98.8 (48.6) |
| $I/\sigma_I$ | 16.31 (1.23) | 11.64 (1.25) | 6.88 (1.38) |
| Completeness (%) | 99.9 (100.0) | 99.6 (99.9) | 99.7 (99.6) |
| Redundancy | 20.2 (20.8) | 3.5 (3.5) | 3.5 (3.6) |
| **Refinement** | | | |
| Total reflections | 50629 (5007) | 99941 (9968) | 101705 (10117) |
| Test-set reflections | 2540 (254) | 5000 (500) | 5080 (508) |
| $R_{work}$[d]/$R_{free}$[e] (%) | 17.02/19.09 | 19.55/22.77 | 19.64/22.88 |
| Number of atoms: | | | |
| Protein | 3241 | 19086 | 19599 |
| Water | 335 | 108 | 616 |
| Ramachandran Plot[f] (%) | 98.3/1.7/0.0 | 98.0/2.0/0.00 | 98.1/1.9/0.0 |
| $B_{average}$ (Å$^2$) | 33.57 | 69.39 | 42.27 |
| Protein | 32.69 | 69.46 | 42.31 |
| Water | 41.97 | 57.17 | 41.19 |
| Bond length RMSD (Å) | 0.008 | 0.009 | 0.007 |
| Bond angle RMSD (°) | 0.89 | 1.01 | 0.85 |
| **PDB ID** | pdb_00008gjr | pdb_00008ee2 | pdb_00008eln |

The data presented here are related to the abbreviated form presented in Table 1.

[a] Values in parentheses correspond to the highest resolution shell.

[b] $R_{meas}$: the redundancy independent R-factor, described in reference [124]

[c] $CC_{½}$: the percentage of correlation between intensities from random half-datasets [125]

[d] $R_{work} = \Sigma_h |F_{obs}(h) - F_{calc}(h)|/\Sigma_h F_{obs}(h)$, $h \in$ {working set}

[e] $R_{free} = \Sigma_h |F_{obs}(h) - F_{calc}(h)|/\Sigma_h F_{obs}(h)$, $h \in$ {test set}

[f] Favored/allowed/outliers
